# The Building Blocks of Early Land Plants: Glycosyltransferases and Cell Wall Architecture in the model liverwort *Marchantia polymorpha*

**DOI:** 10.1101/2025.04.30.651426

**Authors:** H.S. Kang, X. Tong, A. Mariette, M. Leong, C. Beahan, E. Flores-Sandoval, G. B. Pedersen, C. Rautengarten, J.L. Bowman, B. Ebert, A. Bacic, M. Doblin, S. Persson, E.R. Lampugnani

## Abstract

The liverwort *Marchantia polymorpha* has emerged as an important plant model for developmental studies and may become central to elucidate the complex process of cell wall polysaccharide biosynthesis. This study comprehensively analyses the composition and structure of cell wall glycans across eight different *M. polymorpha* tissue types. We show that while the cell walls largely mirror known land plant cell wall composition, they also exhibit some unique characteristics. For example, α-(1,5)-arabinan was prominently present in the sporophyte tissue, which may indicate a specialised role in this life stage. Furthermore, *M. polymorpha* cell walls displayed a remarkably low overall pectin content, yet the abundance of pectic arabinan in sporophytes hint at its putative role in the evolution and complexity of spermatophyte cell walls. Through comparative analyses of glycosyltransferase (GT) families across plant species, we found that *M. polymorpha* has a diversified GT repertoire compared for example, to the model plant *Arabidopsis thaliana*, indicating uniqueness in its cell wall biosynthesis pathways. To support research underpinning cell wall biosynthesis, we developed a Gateway compatible compendium of 93 *M. polymorpha* GTs, providing a valuable resource for genetic and functional studies. Our study thus works as a foundation to drive new insights into cell wall evolution, structure and function across the plant kingdom.

## Introduction

The plant cell wall is an extracellular matrix surrounding the plasma membrane which governs cell growth, and therefore the morphogenesis of plants (Cosgrove, 2016). As the outer-most layer of the cell, the cell wall is an important part of plant immunity and resistance to (a)biotic stresses (Bacete et al., 2018; Houston et al., 2016; Molina et al., 2021; Tenhaken, 2015). The plant cell wall is the most abundant source of biomass on Earth and important as a food, fibre and fuel (Bar-On et al., 2018), and its properties confer many anthropogenically important properties, such as the strength of timber for building materials and as a source of dietary fibre that is essential for gut health (Johnson et al., 2018). Due to its utility, it is of great interest to understand how plant cell wall composition determined at a molecular level.

The plant primary cell wall is composed mostly of polysaccharides and some glycosylated proteins (glycoproteins). Cell wall polysaccharides are further subdivided into three major groups – cellulose, hemicelluloses, and pectins (Lampugnani et al., 2018). Cellulose is a 1,4-β-linked glucose (Glc) homopolymer where 18 glucan chains biosynthesised in parallel form microfibrils, which are the main load bearing structure of the plant cell wall (McFarlane et al., 2014). Hemicelluloses are a diverse group of polysaccharides including xyloglucans, heteromannans and heteroxylans (Scheller & Ulvskov, 2010). They are characterised by the equatorial 1,4-β-linked backbone of Glc, mannose (Man) and xylose (Xyl) respectively, and are substituted further by other sugars and non-glycosyl substituents. Pectins, the other major polysaccharide group, form the hydrated gel-like matrix of the cell wall and are also very important for cell-cell adhesion facilitated by the middle lamella (Atmodjo et al., 2013; Mohnen, 2008). Pectins are also the most heterogeneous plant cell wall polysaccharides, and consist of homogalacturonan, rhamnogalacturonan-I and -II (RG-I and RG-II). Lastly, glycoproteins comprise approx. 10% (w/w) in the primary cell wall and include arabinogalactan proteins (AGPs) and extensins, which are part of the hydroxyproline-rich glycoprotein (HRGPs) superfamily (Nguema-Ona et al., 2014; Strasser et al., 2021). While most cell wall polymers associate through non-covalent interactions (both hydrogen bonds and ionic complexes (e.g. pectins through Ca^2+^)) some components can be covalently attached to one another (Tan et al., 2013, 2022; Doblin et al., 2022).

Despite this overarching understanding of the composition of the cell wall, there are still large gaps in our knowledge about how cell wall polysaccharides and glycoproteins are assembled, and which biosynthetic enzymes and auxiliary proteins underpin their biosynthesis. In part, this is because of the complex network of genes required for the biosynthesis of the many components of the wall. To illustrate, it is estimated that about 15% of genes contribute to the synthesis and remodelling of cell walls in *Arabidopsis* (Carpita et al. 2001), and it is estimated that pectins alone require at least 67 different types of transferase enzymes for their biosynthesis (Mohnen, 2008). Additionally, apart from cellulose, the other classes of polysaccharides display cell/tissue-type, developmental and taxonomic variability (Doblin et al., 2023). For example, xyloglucans in land plants show lineage specific sidechain substitutions (Peña et al., 2008). The heteromannans are further divided into glucomannans, galactomannans and galactoglucomannan depending on their backbone and sidechains (Voiniciuc, 2022). Heteroxylan sub-classes include arabinoxylans, glucuronoxylans and glucouronoarabinoxylans (Curry et al., 2023). Other important cell wall polysaccharides include callose (Doblin et al., 2023; B. Wang et al., 2022), mixed-linkage β-glucans (MLG) (Doblin et al., 2023; S. J. Kim & Brandizzi, 2016) and arabinoglucan (Roberts et al., 2018). Furthermore, hemicelluloses and pectins are commonly substituted with non-glycosyl residues (e.g *O*-acetyl, *O*-methyl, phenolic acids (e.g. ferulic acid)) adding further complexity to the assembly process. The high degree of structural complexity of plant cell wall polysaccharides indicates an equally complex mechanism of biosynthesis.

Plant cell wall polysaccharides are synthesised by glycosyltransferases (GTs), enzymes that catalyse glycosidic bonds between an activated sugar donor and specific acceptors such as proteins, lipids and other sugars (Lairson et al., 2008; Moremen & Haltiwanger, 2019). GTs are catalogued in an online database called CAZy (Carbohydrate Active Enzymes; https://www.cazy.org/), which currently contains over 1 400 000 GTs in more than 130 families (Drula et al., 2022). Plant cell wall biosynthetic GTs belong to 18 well-defined clades which represent subsets of the larger GT superfamily (Table 1). Within these clades, the eudicot model plant *Arabidopsis thaliana* has 288 GT genes. In total, *Arabidopsis* encodes 553 GTs, indicating that over half of its GT genes are candidates in the investigation of cell wall biosynthesis. Plant cell wall biosynthetic GTs use an activated form of monosaccharides, called nucleotide sugars, as donor. Monosaccharides are activated by the addition of a nucleotide, often uridine diphosphate (UDP) or guanine diphosphate (GDP). However, biochemical characterisation of cell wall GT function is made challenging by the high donor and acceptor substrate specificities (Mariette et al., 2023). Furthermore, functional characterization of GTs is hampered by the genetic complexity of *Arabidopsis*, where multigenic families render the investigation of gene function by observing the phenotype of a simple gene knock-out difficult. Additionally, the significant capacity for compensatory wall responses from existing GTs can confuse interpretation, creating a need for a genetically simpler model organism such as the liverwort *Marchantia polymorpha* (*Marchantia*) to explore cell wall biosynthesis.

**Table 1.** Characterised cell wall-associated GT activity across 18 families.

| GT family | Gene ID | Characterised GTs (Polymer, catalytic activity) | References |
| --- | --- | --- | --- |
| 2 | CESA | Cellulose, β-1,4 glucan synthase | (Cho et al., 2015) |
| | CSLC | Xyloglucan, β-1,4 glucan synthase | (Cocuron et al., 2007) |
| | CSLA | Glucomannan, β-1,4 mannan synthase, β-1,4 glucan synthase | (Dhugga et al., 2004; Goubet et al., 2009; Liepman et al., 2005; Voiniciuc et al., 2019) |
| | CSLD | Mannan, β-1,4 mannan synthase | (Verhertbruggen et al., 2011; Yin et al., 2011) |
| | CSLF | Mixed linkage β-glucan, β-1,3; 1,4 glucan synthase | (Burton et al., 2006; Vega-Sánchez et al., 2012; Jobling, 2015; S.-J. Kim et al., 2015; Little et al., 2018) |
| | CSLH | Mixed linkage β-glucan, β-1,3; 1,4 glucan synthase | (Doblin et al., 2009; Little et al., 2018) |
| | CSLJ | Mixed linkage β-glucan, β-1,3; 1,4 glucan synthase | (Little et al., 2018) |
| | CSLF | Xylan, 1,4-β-glucosyltransferase | (Little et al., 2019) |
| 8 | GUX | Xylan, α-1,2 glucuronosyl transferase | (C. Lee et al., 2012; Rennie et al., 2012) |
| | GAUT | Homogalacturonan, α-1,4 galacturonosyl transferase | (Amos et al., 2018; Atmodjo et al., 2011; Biswal et al., 2018; Sterling et al., 2006; Voiniciuc et al., 2018) |
| 14 | GlcAT14 | AGP, β-1,6 Glucuronosyl transferase | (Dilokpimol & Geshi, 2014; Knoch et al., 2013) |
| 29 | GALT29A | AGP, β-1,6 Galactosyl transferase | (Dilokpimol et al., 2014) |
| 30 | KDTA | RG-II, Kdo-transferase | (Delmas et al., 2008; Séveno et al., 2010) |
| 31 | HPGT | AGP, Hydroxyproline-O-galactosyltransferase | (Ogawa-Ohnishi & Matsubayashi, 2015) |
|  | GALT2–6 | AGP. Hydroxyproline-O-galactosyltransferase | (Basu et al., 2013; Basu, Tian, et al., 2015; Basu, Wang, et al., 2015) |
| | GALT31A | AGP, β-1,6 galactosyl transferase | (Geshi et al., 2013) |
| | | AGP, β-1,3 galactosyl transferase | (Ruprecht et al., 2020) |
| | At1g77810 | AGP, β-1,3 galactosyl transferase | (Qu et al., 2008) |
| | KNS4/UPEX1 | AGP/pectin, β-1,3 galactosyl transferase | (Suzuki et al., 2017) |
| | GALT8 | AGP, β-1,3 Galactosyl transferase | (Narciso et al., 2021) |
| 34 | XXT | Xyloglucan, α-1,6 xylosyl transferase | (Faik et al., 2002; Cavalier & Keegstra, 2006; Vuttipongchaikij et al., 2012; Culbertson et al., 2016; Ruprecht et al., 2018) |
| | GMGT | Galactomannan, α-1,6 galactosyl transferase | (Edwards et al., 1994; Voiniciuc et al., 2015) |
| 37 | FUT1 | Xyloglucan, α-1,2 fucosyl transferase | (Cicéron et al., 2016; Faik et al., 2000; Perrin et al., 1999; Rocha et al., 2016; Urbanowicz et al., 2017; Vanzin et al., 2002) |
| | FUT4/6 | AGP, α-1,2 Fucosyl transferase Arabinogalactan | (Wu et al., 2010) |
| 43 | IRX9 | Xylan, Ambiguous | (Mortimer et al., 2015; Urbanowicz et al., 2014) |
|  | IRX14 | Xylan, Ambiguous |  |
| 47 | MUR3 | Xyloglucan, β-1,2 galactosyl transferase | (Madson et al., 2003, 2003) |
| | XGT | Xyloglucan, β-1,2 galactosyl transferase | (Jensen et al., 2012) |
| | XST | Xyloglucan, α-1,2 arabinosyl transferase | (Schultink et al., 2013; Zhu et al., 2018) |
| | XUT | Xyloglucan, β-1,2 galacturonosyl transferase | (Peña et al., 2012) |
| | XGD | Xylogalacturonan, β-1,3 xylosyl transferase | (Jensen et al., 2008) |
| | XYS1/IRX10-L | Xylan, β-1,4 xylosyl transferase | (Jensen et al., 2014, 2018; Jiang et al., 2016; C. Lee et al., 2012; Urbanowicz et al., 2014; H. Wang et al., 2022; Zeng et al., 2010, 2016) |
| | ARAD | Ambiguous, α-1,5 arabinosyl transferase | (Harholt et al., 2006) |
| | ExAD | Extensin, α-1,3 Arabinosyl transferase | (Møller et al., 2017) |
| 48 | GSL | Callose, β-1,3 glucan synthase | (Xie et al., 2011) |
| 61 | XAT1 | Xylan, α-1,3 arabinosyl transferase | (Anders et al., 2012) |
| | XAX1 | Xylan, β-1,2 xylosyl transferase | (Chiniquy et al., 2012) |
| | XAX1 | Xylan, hydroxycinnamoyl-α-1,3-arabinosyl transferase | (Feijao et al., 2022) |
| 77 | RGXT | RG-II, α-1,3 xylosyl transferase | (Egelund et al., 2006, 2008; Liu et al., 2011; Petersen et al., 2009) |
| | RRA | Extensin, β-1,2-arabinosyl transferase | (Velasquez et al., 2011) |
| | XEG113 | β-1,2-Arabinosyl transferase | (Velasquez et al., 2011) |
| | RAY1 | AGP, β-Arabinofuranosyl transferase | (Gille et al., 2013) |
| 92 | GALS | Galactan, β-1,4 galactan synthase | (Ebert et al., 2018; Liwanag et al., 2012) |
| | | Galactan, α-arabinosyl transferase | (Laursen et al., 2018) |
| 95 | HPAT | Extensin, O- β-arabinofuranosyl transferase | (Ogawa-Ohnishi et al., 2013) |
| 96 | SGT1 | Extensin, Serine O-α-galactosyl transferase | (Saito et al., 2014) |
| 106 | MSR | (Gluco)mannan, O-fucosyltransferase | (D. K. Smith et al., 2018; Y. Wang et al., 2013) |
| | RRT | RG-I, α-1,4 rhamnosyl transferase | (Takenaka et al., 2018; Uehara et al., 2017) |
| 116 | RGGAT1 | RG-I, α-1,2-galacturonosyltransferase | (Amos et al., 2022) |

Liverworts are a clade within Bryophytes, a monophyletic division which diverged from the land plant common ancestor around 470 million years ago (Strother & Taylor, 2018). *Marchantia* in particular has gained traction as a popular model organism for the availability and applicability of many genetic tools, the ease of growing it in laboratories and importantly, its low genetic redundancy (Bowman, 2022; Bowman et al., 2017; Ishizaki et al., 2016). For example, *Marchantia* has 130 protein coding genes in the 18 families where cell wall biosynthetic GTs have been identified; less than half compared to *Arabidopsis.* Despite having fewer genes, most cell wall polysaccharides found in land plants are conserved in *Marchantia* (Happ & Classen, 2019; Henry et al., 2020; Kolkas et al., 2023; Pfeifer et al., 2022), as well as the genetics behind cell wall biosynthesis, as exemplified by the functional conservation of the RG-I rhamnosyltransferase *RRT1* in *Marchantia* (Wachananawat et al., 2020). Thus, the seemingly conserved nature of *Marchantia* cell wall polysaccharides and *Marchantia’s* low genetic redundancy makes it an attractive model organism to study the complex processes of plant cell wall polysaccharide biosynthesis. Additionally, investigating the cell wall structure in *Marchantia* sheds light on the lineage-specific innovations or adaptations. For example, the xyloglucans in *Marchantia* and mosses uniquely include the galacturonic acid containing sidechains P and Q (Peña et al., 2008), and the accumulation of secondary metabolites in the cell wall, possibly to subdue environmental stresses related to terrestrialisation (Jibran et al., 2024). In this study, we aim to gain a better understanding and overview of the cell wall biosynthetic precursors, GTs and polysaccharides across different *Marchantia* tissue types and establish *Marchantia* as a model organism for cell wall biosynthesis by creating a cDNA library of GTs which can be used for recombination into a variety of expression vectors to expedite functional characterisation. Furthermore, we have undertaken a comprehensive phylogenetic analysis to infer the degree of conservation and diversification of cell wall biosynthetic proteins in *Marchantia*.

## Methods

### Plant Material

Sporangia from an Australian population of *Marchantia polymorpha* were collected from a southeastern Melbourne field location (37°57’48.36”S, 145° 6’20.41”E), sterilised, suspended in water and plated on petri dishes as previously described (Flores-Sandoval et al., 2015). Plants were grown in a control temperature room for the stipulated period under long day conditions following Lee et al., (2014).

### Glycosidic linkage Analyses

Cell wall preparations (as alcohol-insoluble residue, AIR) and methylation analyses of neutral carbohydrates followed the methods described by Pettolino et al. (2012). Briefly, dried *Marchantia* tissue samples were extracted consecutively with 70% (v/v) ethanol (3x), chloroform:methanol (1:1), 100% methanol and the tissue either stored in 100% ethanol or washed with 100% acetone before air drying to generate alcohol-insoluble residue (AIR). AIR samples were de-starched before performing linkage analyses. AIR preparations were carboxyl reduced to include uronic acid and methyl-esterified uronic acid residues. Monosaccharide linkage analysis was performed by multiple methylation (2x), hydrolysis, reduction and acetylation to generate partially methylated alditol acetates that were separated and identified by GC-MS. Polysaccharide levels were estimated according to Pettolino et al. (2012) by addition of appropriate deduced monosaccharide linkages for each polysaccharide based on knowledge from the wider literature, largely based on angiosperms and gymnosperms.

### Profiling of Nucleotide Sugars

Nucleotide sugar measurements were performed as described in (Rautengarten et al., 2019). In brief, 50 mg of *Marchantia* thallic tissue were ground in liquid nitrogen, resuspended in ice-cold methanol/chloroform (7:3), and incubated at −20 °C for 2 hours. After adding 400 µL of ice-cold water, the samples were centrifuged at 20,000 × g at 4 °C, and the upper phase was collected into 15 mL tubes on ice. The water addition and centrifugation were repeated twice, following that all supernatants were combined. Then the samples were frozen in liquid nitrogen and freeze-dried overnight. Purification was done via solid phase extraction, followed by detection and quantification of nucleotide sugars on a SCIEX 4000 QTRAP system with a TurboIonSpray source and Agilent 1100 Series Capillary LC System. Nucleotide sugar standards included UDP–α-D-xylose, UDP–β-L-arabinopyranose, UDP–α-D-galacturonic acid, UDP–α-D-glucuronic acid, UDP–α-D-glucose, UDP–α-D-galactose, UDP–N-acetyl-α-D-glucosamine, UDP–N-acetyl-α-D-galactosamine, GDP–α-D-mannose, GDP–β-L-fucose, GDP-Glc (Sigma-Aldrich), and UDP–β-L-arabinofuranose (Peptides International) (Rautengarten et al., 2014). Data were analysed using Analyst 1.5.1 and quantified with MultiQuant 2.1 through the linear regression of the peak areas (Rautengarten et al., 2016).

### Immunolabeling of Cell Wall Epitopes and Fluorescence Microscopy

Fresh tissue samples were dissected and fixed and subjected to dehydration through an ethanol series. Sample were then post-processed through an ethanol-HistoClear (National Diagnostics) series in 1 hr intervals (25%, 50%, 75%, 3x 100% HistoClear) and a HistoClear-Paraplast Plus (ProSci Tech) series in half day intervals (50%, 6x 100% Paraplast Plus) with the final embedding in 100% Paraplast Plus. 6-8μm thick sections were cut using a Leica 2040 microtome and then mounted on slides. Wax was subsequently removed from the tissues with xylene washes and samples rehydrated by immersing the slides through an ethanol series with three final washes in deionized water. Immunolabelling using monoclonal antibodies LM6 and LM13, which labels short, branched arabinan and long, unbranched arabinan respectively (Verhertbruggen et al., 2009), LM15 which labels XXXG epitopes of xyloglucan (Marcus et al., 2008,Pedersen et al., 2012) (PlantProbes), BS-400-3 which labels 1-4-β-D-mannan (Pettolino et al., 2001) (Biosupplies Australia) and carbohydrate binding module 3a (CBM3a) (Hernandez-Gomez et al., 2015) (PlantProbes) was carried out as described previously (Lampugnani et al., 2013) except that the samples were counterstained with calcofluor and Pontamine Fast Scarlet 4B (S4B), which binds a range of β-glycans (Anderson et al., 2010). Fluorescence imaging was carried out following Kesten et al. (2019) with a Nikon Eclipse Ti-E inverted Microscope fitted with a CSU-W1 Yokogawa spinning disc head and an Evolve EM-CCD camera (Photometrics Technology). Images were collected using the average of eight optical slices.

### Glycosyltransferase Identification and Phylogenetic Analysis

To identify GTs for the phylogenetic analysis, protein sequences of *M. polymorpha* (MarpolBase, http://marchantia.info, TAK-1 v.7), *A. thaliana* (TAIR, https://www.arabidopsis.org/), *Brachypodium distachyon* (Phytozome, v3.2, The International Brachypodium Initiative, 2010), *Selaginella moellendorffi* (JGI PhycoCosm, v.1, Banks et al., 2011), *Physcomitrella patens* (Phytozome, v3.3, Lang et al., 2018) *Chara braunii* (NCBI, GenBank assembly: GCA_003427395.1, Nishiyama et al., 2018), *Closterium* (NCBI, Genbank assembly: GCA_949281275.1, Kawaguchi et al., 2023), *Mesotaenium endlicherianum* (JGI, SAG 12.97, Cheng et al., 2019) and *Chlamydomonas reinhardtii* (JGI, CC-4532 v6.1, Craig et al., 2023) were acquired from the respective sources. Orthogroups within the nine species were identified using Orthofinder v.2.5.5 (Emms & Kelly, 2019), using DIAMOND as the sequence search method. Orthogroups with GT sequences were identified using *Arabidopsis* and *Marchantia* GT sequences curated in CAZy. The sequences from these orthogroups were used to construct the phylogenetic trees for the respective GT families.

To construct the phylogenetic trees, the GT sequences were aligned with clustal omega (1.2.4; Sievers and Higgins, 2018) and the alignments were trimmed with trimal (1.4.1; Capella-Gutiérrez et al., 2009) using the -automated1 option. These trimmed alignments were used to build the phylogenetic trees with RaXML (8.2.12; Stamatakis, 2014) with 1000 Bootstrap using the following command: raxmlHPC -T 4 -f a -x 12345 -# 1000 -p 12345 -m PROTGAMMAAUTO -s output_trimalignment.fa -n Boot. The phylogenetic trees were visualized with iTOL (Letunic and Bork, 2021) and the figures were manually annotated.

### Cloning Procedures

RNA was isolated from *Marchantia* and first-strand cDNA synthesis was carried out as described previously (Lampugnani et al., 2016). Coding sequences (CDS) of target genes were retrieved from the MarpolBase (Bowman et al., 2017) and gene specific primers designed for cloning into pDONR221 or alternatively pENTR d-TOPO using the Gateway Technology. Sequences of primers used in this study are shown in Table S1. Gene products were amplified using KOD Hot Start (Merck) or Q5 Hot-Start (NEB) and subsequently purified using a QIAquick PCR purification kit (Qiagen) prior to transformation. Cloned products were verified using Sanger sequencing with m13F and m13R primers. Entry clones with Mp3g24900 and Mp5g17760 coding sequences were subcloned into pEARLEYGATE101 by LR Clonase II reaction (Thermo Fisher Scientific).

### *Nicotiana benthamiana* infiltration and co-localisation

Transient transformation in *N. benthamiana* was carried out as described in Lampugnani et al., (2016). The Golgi and Endoplasmic Reticulum (ER) organellar markers used for co-localisation are CD3-967 and CD3-953 from Nelson et al., (2007) respectively.

## Results and Discussion

The identification of plant cell wall biosynthetic enzymes is often hampered by high genetic redundancy in the eudicot model organism, *A. thaliana*. Therefore, the use of *Marchantia* as a genetically simpler model system could expedite the elucidation of these enzymes. To do so, we wanted to investigate the substrate availability for cell wall biosynthesis, perform an in-depth investigation of the distribution of cell wall epitopes across different tissue types in *Marchantia* and conduct a phylogenetic analysis of plant cell wall GT candidates.

### *Marchantia* thallus has most sugar precursors required for the biosynthesis of a typical land plant wall

To examine whether *Marchantia* contains the corresponding substrates for the typical land plant cell wall polysaccharide biosynthesis, we conducted a nucleotide sugar analysis on mature thallus tissue. This revealed that *Marchantia* contains high levels of UDP-Glucose (UDP-Glc), with 50 pmol·mg^-1^ of fresh weight representing about 66% of the total nucleotide sugar content (Table 2). As a comparison, the UDP-Glc pool is typically about 53% in *Arabidopsis* (Rautengarten et al., 2014). Interestingly, UDP-Galacturonic acid (UDP-GalA) contents were relatively low at about 1.5 pmol·mg^−1^ of fresh weight representing about 1.9% of the total nucleotide sugar pool. This is about half of the amount of UDP-GalA content detected in *Arabidopsis* tissues (Rautengarten et al., 2014). Further, there are comparatively low levels of UDP-Rhamnose (UDP-Rha) (1.7 pmol·mg^−1^ of fresh weight) in the *Marchantia* thallus when compared to *Arabidopsis*. As we confirm that the *Marchantia* thallus contains the sugar precursors required for the biosynthesis of a conventional land plant cell wall, albeit in differing proportions, we next delved into the characterisation its cell wall polysaccharide composition.

**Table 2.** Nucleotide sugar analysis of mature *Marchantia* thallus. Measurements are an average of eight technical replicates. Standard deviation values are given in brackets.

| Nucleotide sugar | pmol·mg<sup>-1</sup> fresh weight | % total |
| --- | --- | --- |
| UDP-α-D-Glucose | 50.0 (4.9) | 65.5% |
| UDP-α-D-Galactose | 5.2 (0.4) | 6.8% |
| UDP-β-L-Rhamnose | 1.7 (0.2) | 2.2% |
| UDP-α-D-Glucuronic acid | 3.9 (0.1) | 5.1% |
| UDP-α-D-Galacturonic acid | 1.5 (0.1) | 1.9% |
| UDP-α-D-Xylose | 2.0 (0.3) | 2.6% |
| UDP-β-L-Arabinopyranose | 2.6 (0.2) | 3.4% |
| UDP-β-L-Arabinofuranose | 0.6 (0.1) | 0.8% |
| UDP-α-D-N-Acetylglucosamine | 5.4 (0.6) | 7.1% |
| GDP-β-L-Fucose | 0.1 (0.0) | 0.1% |
| GDP-α-D-Mannose | 2.5 (0.5) | 3.3% |
| GDP-α-L-Galactose | 0.4 (0.1) | 0.5% |
| GDP-α-D-Glucose | 0.4 (0.0) | 0.5% |
| Total | 76.3 |  |

### The cell wall composition of *Marchantia* shows high pectin content in young tissues

We aimed to establish the glycosyl composition of the *Marchantia* cell wall using linkage analysis by methylation in eight different tissue types; to investigate whether there are tissue specific differences that could highlight the importance of some polysaccharides in different biological contexts. We assayed dormant gemmae, eight-day-old gemmaling, mature thallus, antheridiophore heads, archegoniophore stalks, fertilised archegoniophore heads, sporophytes, and seven-day-old sporelings (Table S2). Linkage analysis allows the identification of individual glycosyl linkages of the wall, which can be used to predict the polysaccharide composition of the wall. However, while cell wall polysaccharides can be distinguished by their unique glycosyl linkages, it is challenging to unequivocally assign linkages to particular polysaccharides, especially in species which may have uncharacterised polymers. With this in mind, we assigned linkages largely based on cell wall polymers in other land plants (Table 3). A summary of the calculated monosaccharide content of each of the tissue samples is presented in Figure S1.

**Table 3.** Polysaccharide deductions from the methylation analysis data according to Pettolino et al. (2012). Abundance is colour-coded from white (lowest) to green (highest). tr = trace (<0.1%), n.d. = not detected.

|  | Dormant<br>Gemma | 8d<br>Gemmaling | Mature<br>Thallus | Antheridio-<br>phore<br>Heads | Archegonio<br>-phore<br>Stem | Archegonio<br>-phore<br>Heads | Sporophyte | 7d<br>Sporelings |
| --- | --- | --- | --- | --- | --- | --- | --- | --- |
| Arabinan | 4.8 | 0.7 | 1.9 | 1.5 | 1.2 | 1.1 | 14.5 | 0.9 |
| Type I AG | 2.6 | 6.2 | 8 | 1.5 | 1.5 | 1.4 | 0.9 | 10.1 |
| Type II AG | 4.2 | 12.2 | 5.5 | 1.1 | 1.9 | 1.4 | 1.4 | 6.2 |
| Homogalacturona<br>n | 1 | 0.4 | 2.2 | 1.6 | 1.3 | 1 | 1.9 | 1.3 |
| RG-I | 0.7 | 3.3 | 1.2 | 0.8 | 0.7 | 0.6 | 1.3 | 4.1 |
| GAX | 4.6 | 7.9 | 5.4 | 2.9 | 7.3 | 2.4 | 11.7 | 5.8 |
| Heteromannan | 12.7 | 4.6 | 3.8 | 20.1 | 19.1 | 24.4 | 4.6 | 4.9 |
| Xyloglucan | 15.9 | 11.6 | 13.5 | 16.1 | 17.6 | 11.9 | 19.5 | 12.6 |
| Cellulose | 40.6 | 35.6 | 36.1 | 43.3 | 39.6 | 44.5 | 32.5 | 38.9 |
| Callose | tr | 1.7 | 0.4 | tr | tr | tr | tr | 0.5 |
| Extensin | n.d. | 0.8 | tr | n.d. | n.d. | n.d. | n.d. | 1.1 |
| Unassigned | 12.8 | 14.9 | 21.9 | 11.1 | 9.8 | 11.3 | 11.8 | 13.6 |

The dominant linkage was 1,4-Glc, which made up between ∼39 to ∼57 mol% of the wall in all our samples (Table S2). This is more than what was found in *Arabidopsis* (33.8 mol%) (Pettolino et al., 2012), but similar to what was detected in the moss *P. patens* (48.6 mol%) (Moller et al., 2007). The 1,4-Glc linkages constitute cellulose, the backbone structure of xyloglucans and some heteromannans, such as galactoglucomannans. The β-(1,4)-Glc-linked backbone of xyloglucan is, at a minimum, substituted with α-1,6-Xyl side chains. As such, the assignment of 1,4-Glc to xyloglucan should be proportional to the number of 1,4,6-Glc linkages. Notably, the 1,4,6-Glc linkage was about 4-6 mol% in all tissues (Table S2).

Mannans (β-(1,4)-linked mannan) are distinguished from (galacto)glucomannans by having only β-(1,4)-Man linkages incorporated into their backbone (Voiniciuc et al., 2019; Voiniciuc, 2022). In the polysaccharide deduction method of Pettolino et al (2012), the assignment of the 1,4-Glc linkage to (galacto)glucomannans hinges on the amount of 1,4-Man. In samples with rapid rates of cell division, such as the gemmae and sporelings, we detected 1,4-Man at relatively low levels, i.e. ∼1-2 mol%, compared with ∼40% mol% 1,4-Glc. In older plant samples, such as the mature thallus, antheridiophore and archegoniophore, the proportion of 1,4-Man was significantly higher, reaching up to 9 mol% in the archegoniophore stalk (Table S2). Hence, the major portion of the 1,4-Glc linkage is likely associated with cellulose and xyloglucan in younger tissues with an increasing level associated with (galacto)glucomannas in mature tissues. Assuming that the remaining 1,4-Glc linkages are assigned to cellulose, this would suggest that most *Marchantia* tissues have cell walls composed of 35 to 45 mol% cellulose (Table 3). Another linkage that typically represents a defined polysaccharide is 1,4-Xyl. This linkage is associated with heteroxylans and its derivatives, such as (glucurono)arabinoxylan (GAX). We found that 1,4-Xyl linkages were most abundant in the sporophyte, at around 8.3 mol%, with lesser amounts in other tissue types (Table S2).

Glycosyl linkages associated with arabinogalactans (AG), such as 1,4-Gal, 1,3-Gal and 1,6-Gal are present in higher levels in younger tissue types (Table 3). Type I AG refers to polysaccharides with a β-1,4-Gal backbone (Clarke et al., 1979; Hinz et al., 2005). Type I AGs are found as sidechains of RG-I and ranges in *Marchantia* between 0.9-10.1 mol%, with the highest levels being associated with younger tissue types. Type II AGs possess a β-1,3-Gal backbone branched with β-1,6-Gal residues, and are found on AGPs (Clarke et al., 1979; Showalter, 2001; Seifert & Roberts, 2007; Tan et al., 2012). Like type I AGs, type II AGs range between 1.1-12.2 mol%, with the higher range associated with younger tissue types (Table 3). However, the 1,4-GalA linkage, indicative of the pectic polymer homogalacturonan and RG-I, was found in low levels across all tissue types (Table S2, Table 3). The range of HG is estimated to be 0.4-1.9 mol%. Similarly, RG-I backbone related linkages, 1,2,4-linked Rha and equal amounts of 1,4-GalA only constituted between 0.6-4.1 mol% of the cell wall polysaccharides across the different tissue types; with highest levels associated with dormant gemmaling and cultured sporelings. The sporophyte is particularly interesting because we detected unusually high levels (∼12%) of the linkage 1,5-Ara (arabinose) (Table S2).

Glycosyl linkage analysis of 7-day old sporelings allow us to draw interesting comparisons between *Marchantia* and another model bryophyte organism *P. patens* and its spore-derived tissue (Moller et al., 2007). Linkage analysis of the 6-8 day old protonemal tissue of *P. patens* showed high levels of total GalA (∼14 mol%) compared to our results of 7-day old *Marchantia* sporelings (4.2 mol%), but relative Ara and Xyl levels are higher in *Marchantia* sporelings (7.8 mol%, 7.2 mol% respectively) than in *P. patens* (mol 2.5% and 4.1 mol% respectively) (Moller et al., 2007). Notably, relative Gal (galactose) content is higher in sporelings (23.1 mol%) than *P. patens* protonema (11 mol%) (Moller et al. 2007). The high abundance of Gal is also noticeable in dormant gemmae and 8-day old gemmalings (22.4 mol% and 19.4 mol% respectively), which indicate an important role of Gal-containing cell wall polysaccharides in *Marchantia*.

Despite their shared status as early-diverging lineages of land plants, the difference in the cell wall monosaccharide composition between *Marchantia* and *P. patens* exemplifies the prevalence of taxonomic diversity in cell wall architecture. As such, while the suite of fundamental building blocks of plant cell walls is comparable across land plants, there are indeed notable differences in their relative composition. For example, dicots and non-commelinoid monocots have a cellulose and xyloglucan-dominant framework, embedded in relatively pectin-rich cell walls, also known as ‘Type I’ walls (Carpita & Gibeaut, 1993, Doblin et al., 2023). In contrast, commelinoid monocots, including the members of the Poaceae, have lower pectin content, but higher levels of the hemicellulose GAX, and are called ‘Type II’ cell walls (Carpita and Gibeaut 1993, Scheller and Ulvskov 2010, Doblin et al., 2023). Moreover, grasses have high amounts of MLGs (Doblin et al., 2023; Vega-Sánchez et al., 2013), basal land plants such as eusporangiates are rich in mannan (Silva et al., 2011, Harholt et al. 2012), and Gymnosperm cell walls are characteristic for their abundance of the hemicellulose galactoglucomannans (Voiniciuc et al. 2022). The *Marchantia* cell wall composition appears distinct from other land plant lineages in that its dominant hemicelluloses are xyloglucans and heteromannans and that it has lower levels of pectic polysaccharides and glycoproteins in thallus tissue, which is also consistent with the recent study by Jibran et al., (2024). In addition, the glycosyl linkage analysis across the different life stages revealed that pectin and arabinogalactan polysaccharide content are increased in younger tissue types, and highlights the abundance of Gal-containing cell wall polysaccharides Taken together, our data indicate that *Marchantia* contains polysaccharide linkages representing all cell wall polymers in found in land plant walls, further supporting *Marchantia* as a suitable cell wall model organism for land plants.

An interesting aspect of the *M. polymorpha* cell wall is that it lacks ‘true’ lignin in secondary cell walls (Kremer et al., 2004; Pfeifer et al., 2022). Briefly, *M. polymorpha* possesses the monolignol precursors of lignin subunits *p*-hydroxyphenyl, guaiacyl and syringyl alcohol, and its cell wall contains high level of polyphenols (Espiñeira et al., 2011). A recent study revealed that a type of flavonoid called auronidin is tightly bound to the cell wall, perhaps contributing to cell wall integrity (Jibran et al., 2024). Based on the accumulation of auronidin and possibly other polyphenolic compounds, the authors speculated its role to be similar to that of lignin in tracheophytes, such as anti-microbial effects (Carella et al., 2019; Humphreys et al., 2010), reinforcement of the primary cell wall and protection from excess light (Jibran et al., 2024). In this study, only sugar-containing cell wall polymers have been considered. However, it raises the question of whether sugar-containing polymers may have to ensure certain functionalities in *M. polymorpha,* that in other land plants are mediated by lignin-strengthened cells.

### Immunolocalisation of cell wall epitopes reveals potential tissue specific functions of select cell wall polysaccharides

Our polysaccharide linkage analyses gave us a comprehensive overview of the types and quantities of polysaccharides in different *Marchantia* tissues, however, the overview lacks information on the spatial distribution of the polysaccharides within. To further corroborate the linkage analyses and to assess where certain polysaccharides are deposited, we undertook immunofluorescence labelling experiments with antibodies that bind to specific cell wall polysaccharide epitopes. For better visualisation, sections were counter stained for better visualisation with S4B specific to cellulose (Hoch et al., 2005) and calcofluor white which binds more promiscuously to several cell wall epitopes (Anderson et al., 2010). To ensure that patterns of polysaccharide deposition could be compared between sample types, at least on a relative level, we adjusted the imaging settings to ensure that there were no saturated pixels in the images and maintained constant settings between sample types, which may lead to false negatives for epitopes with low abundance. As a consequence, the lack of signal in a specific tissue does not necessarily mean that the epitope is absent, as the settings may simply have been attenuated below detectable levels or some epitopes might be blocked from detection by other polymers.

When the thallus was probed with either CBM3a, LM15 or BS-400-3 (β-1,4-mannan specific antibody), the corresponding signal was detected throughout the entire cross-section of the thallus, including the upper epidermis, lower epidermis, some parenchymous tissue and rhizoids, while the LM6 and LM13 arabinan antibodies were largely undetected (Figure 1). We used two antibodies binding to arabinans, the LM6 antibody recognising a linear pentasaccharide in (1,5)-α-L-arabinans and the LM13 antibody recognising longer stretches of 1,5-linked arabinosyl residues that are likely to be more abundant in unbranched arabinans (Verhertbruggen et al., 2009). The absence of LM6 and LM13 binding in the rhizoids is intriguing, as LM6 shows binding to bundled rhizoids in the archegoniophore stalk (Figure 2). The rhizoids are structurally intact to allow the binding of probes and antibodies in the thallus section, as shown by the binding of CBM3a and S4B. Whether there is compositional variation between the rhizoids associated with the thallus and rhizoids associated with the stalk requires further investigation. The bundled rhizoids in the stalk of the archegoniophore also showed binding of the CBM3a epitope (Figure 2). LM15 xyloglucan epitopes were detected in the assimilatory filaments of the stalk, and the signal of the mannan antibody was detected extensively throughout the assimilatory filaments, bundled rhizoids and parenchymous tissue (Figure 2E).

**Figure 1.**
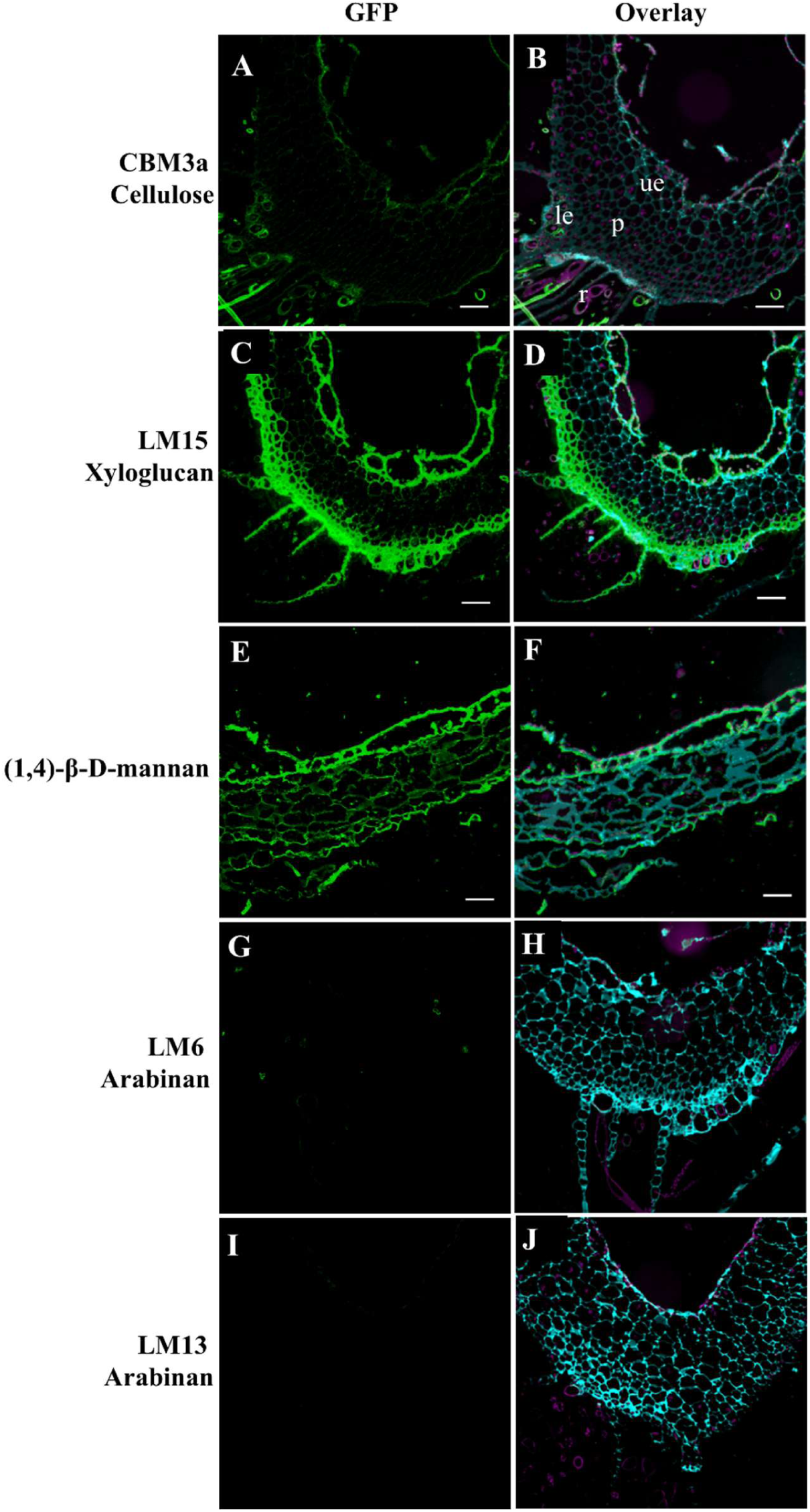
Immunolabelling of the *Marchantia* thallus with A) CBM3a (Cellulose), C) LM15 (Xyloglucan), E) β-1,4-Mannan and G) LM6 (Arabinan) I) LM13 (Arabinan) antibodies, overlayed with counterstains calcofluor white (cyan) and S4B (magenta) in B, D, F H and J respectively. ue = upper epidermis, le = lower epidermis, p = parenchymous tissue. Scale bar = 100 µm

**Figure 2.**
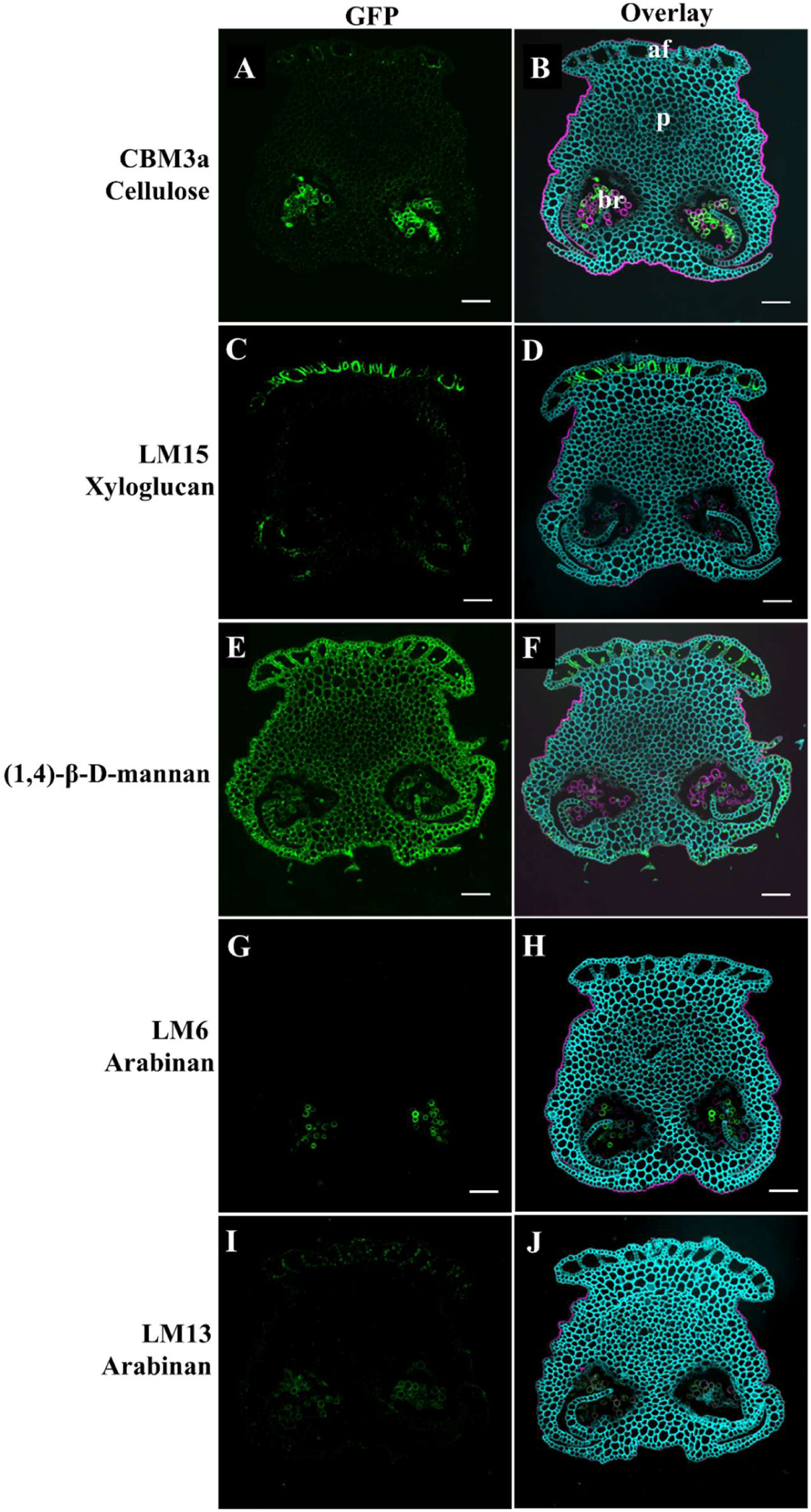
Immunolabelling of the *Marchantia* stalk with the A) CBM3a (Cellulose), C) LM15 (Xyloglucan), E) β-1,4-Mannan, G) LM6 (Arabinan) and I) LM13 (Arabinan) antibodies, overlayed with counterstains calcofluor white (cyan) and propodium iodide (magenta) in B, D, F, H and J, respectively. br = bundled rhizoids, af = assimilatory filaments, p = parenchyma. Scale bar = 100 µm

CBM3a probes of fertilised archegoniophores localised to developing archegonia and the calyptra, a layer which protects the developing sporophyte (Figure 3). LM15 bound to the assimilatory filaments, parts of the stalk and the spore jacket cells. The mannan antibody bound to the archegoniophore extensively, including the assimilatory filaments, spore jacket cells, the calyptra and the perigynium. The LM6 and LM13 antibodies recognizing arabinan epitopes showed binding to the spore capsule, developing spores as well as the foot of the sporophyte. The presence of LM6 epitopes in the sporophytes, which corroborates our glycosyl linkage analysis was also consistent with previous data (Dierschke et al., 2024). A closer inspection of the sporophyte capsule showed binding of the CBM3a probe and the mannan antibody to the helical secondary cell wall structure of elaters (Figure 4). While LM6 and LM13 also recognized arabinan epitopes in the elaters, with a distinct pattern from that of the CBM3a probe and the mannan antibody, in that it does not label the helical patterning of the secondary cell wall but rather co-localises to the tips of the elaters (Figure 4E, G). The CBM3a showed binding to the assimilatory filaments and antheridial chambers of the antheridiophore (Figure 5). The LM15 xyloglucan epitope was more restricted, whereas the mannan epitope was detected extensively in the antheridiophore receptacle. The LM6 and LM13 binding was weak across all cells, except for moderate LM13 binding on the dorsal surface of the antheridiophore overlapping with S4B staining.

**Figure 3.**
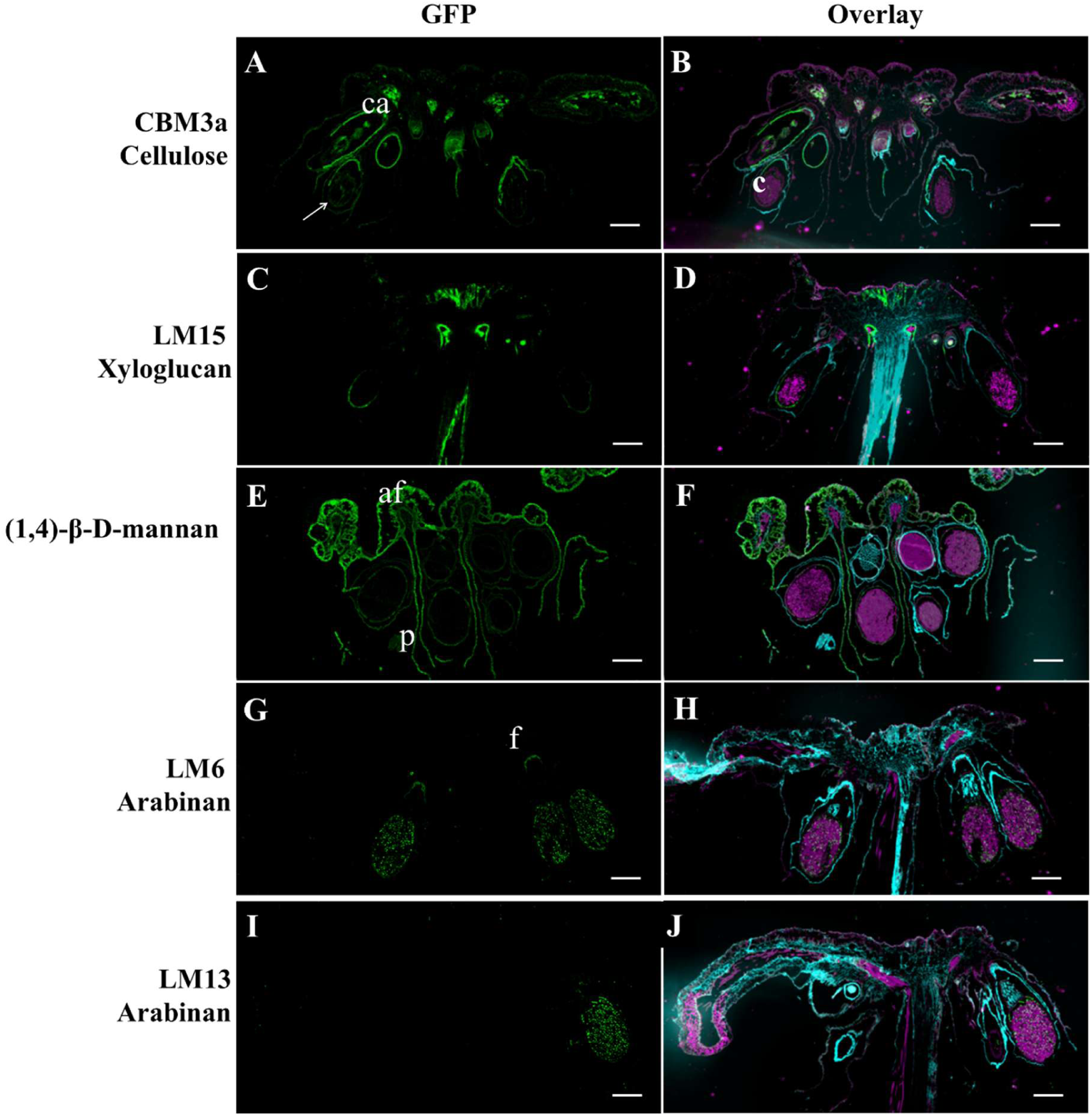
Immunolabelling of the *Marchantia* archegoniophore with the A) CBM3a (Cellulose), C) LM15 (Xyloglucan), E) β-1,4-Mannan and G) LM6 (Arabinan) antibodies, overlayed with counterstains calcofluor white (cyan) and S4B (magenta) in B, D, F and H respectively. c = spore capsule, ca = calyptra, p = perigynium, f = foot, af = assimilartory filaments, arrow = spore jacket cells. Scale bar = 1 mm

**Figure 4.**
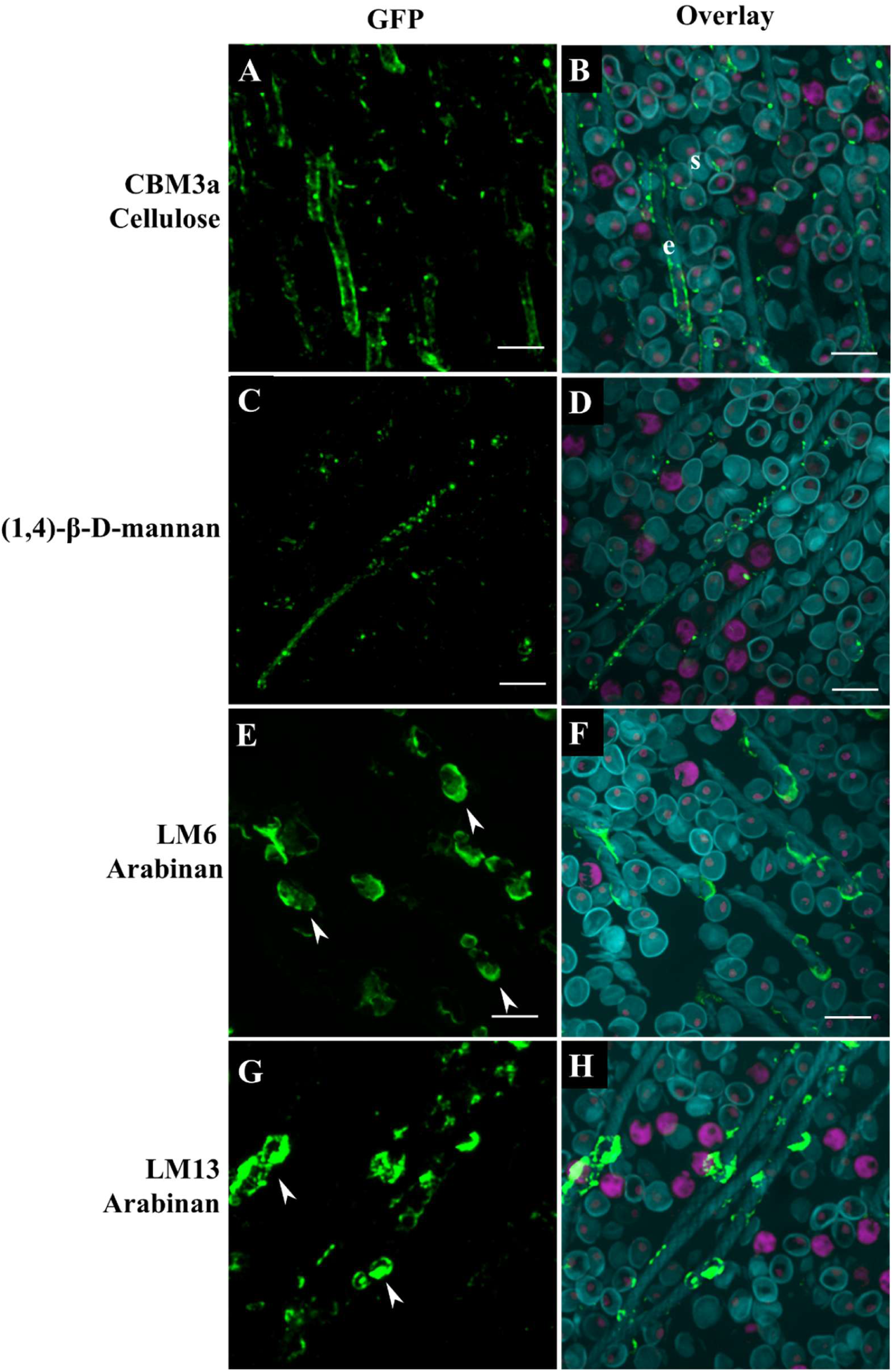
Immunolabelling of the *Marchantia* sporophyte with the A) CBM3a (Cellulose), C) β-1,4-Mannan, E) LM6 (Arabinan) and G) LM13 (Arabinan) antibodies, overlayed with counterstains calcofluor white (cyan) and S4B (magenta) in B, D, F and H respectively. s = spore, e = elaters, arrowheads = tip of elaters. Scale bar = 20 µm

**Figure 5.**
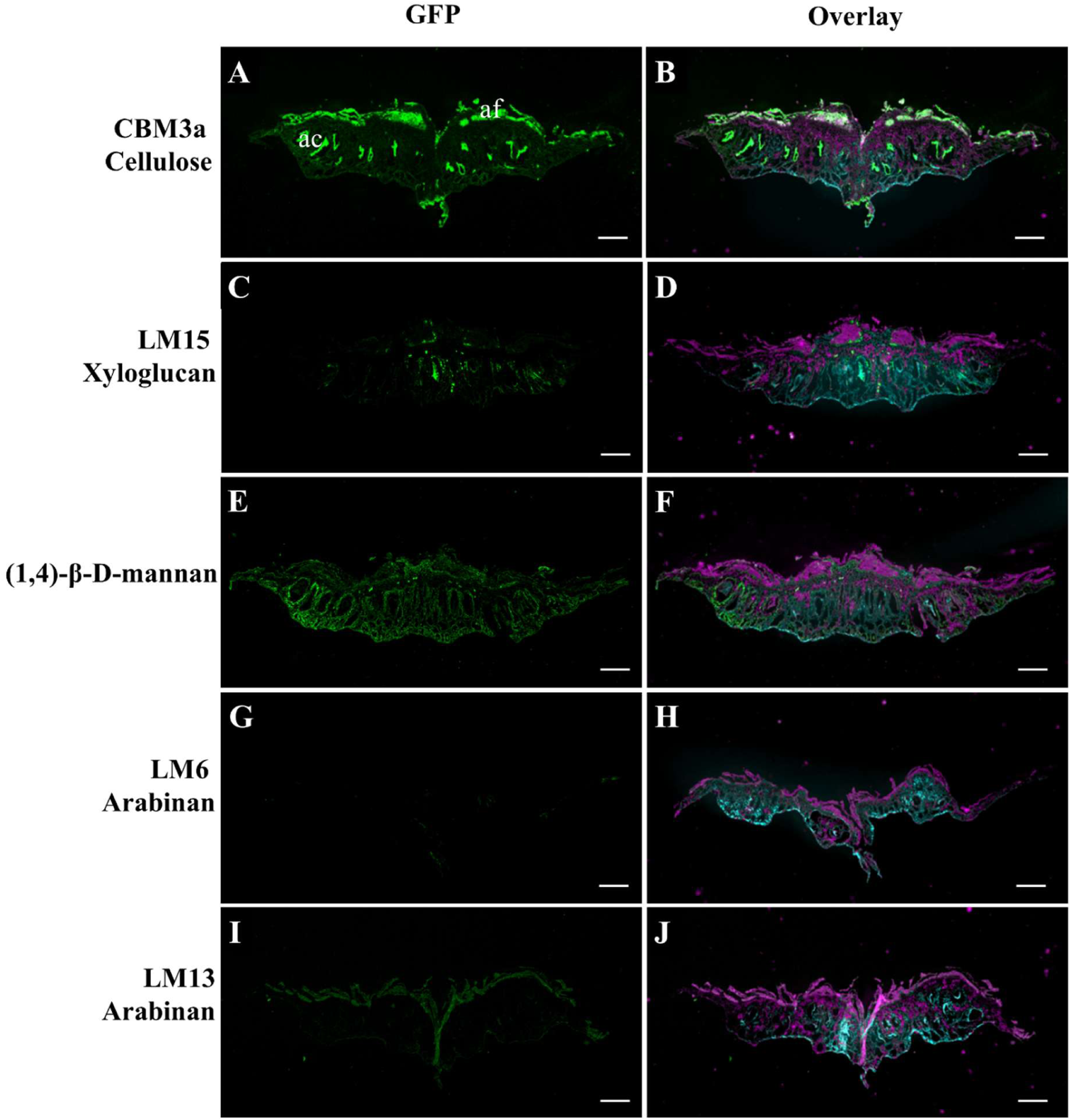
Immunolabelling of the *Marchantia* antheridiophore with the A) CBM3a (Cellulose), C) LM15 (Xyloglucan), E) β-1,4-Mannan, G) LM6 (Arabinan) and I) LM13 (Arabinan) antibodies, overlayed with counterstains calcofluor white (cyan) and S4B (magenta) in B, D, F, H and J respectively. af = assimilatory filaments, ac = antheridial chambers. Scale bar = 1 mm

The tissue-specific deposition of arabinan may indicate a specialised role. Previous studies have associated arabinan with a role in tip-growth, cell wall flexibility and desiccation tolerance (Jones et al., 2003; Parre & Geitmann, 2005; Moore et al., 2008; Carroll et al., 2022). The rhizoids and elaters of *Marchantia* both exhibit tip growth (Shimamura, 2016; Cao et al., 2019), and the detection of the LM6 antibody is consistent with that of other tip-growing cells such as *P. patens* protonemal cells and pollen tubes (K. J. D. Lee et al., 2005; Lampugnani et al., 2016). Both rhizoids (pegged) and elaters also undergo programmed cell death and desiccation (Shimamura, 2016). Due to the hydrophilic properties and ability to function as a physical plasticiser, arabinan has been postulated to act as a buffer to withstand the high mechanical stresses experienced while undergoing desiccation, which is a strategy used by desiccation tolerance plants (Moore et al., 2013). We therefore speculate that the deposition of arabinan in the rhizoids and elaters of *Marchantia* may be related to controlling the extensibility of the cell wall, and maintaining cell integrity during desiccation. However, it is uncertain which polysaccharides, if any, the arabinan is part of. The α-1,5-Ara linkage is associated with the pectic polysaccharide RG-I, AGPs (Happ & Classen, 2019; K. J. D. Lee et al., 2005) or with a linear arabinan in a ‘free’ form that is not part of RG-I (Lampugnani et al., 2016). Interestingly, LM6 does not bind to the α-1,5-linked arabinan sidechains of *Marchantia* AGPs (Happ and Classen, 2018) due to their short length (DP1-2), as LM6 recognizes three or more α-1,5-linked arabinosyl residues (Verhertruggen et al. 2009). Also, our glycosyl linkage analysis indicates a low proportion of predicted RG-I levels in the stalk (0.6 mol%) and sporophytes (1.3 mol%). Thus, more work is needed to conclude whether arabinan detected in the rhizoids exists in either a ‘free’ form, or associated with other polysaccharides.

In combination with the glycosidic linkage results, the immunolabelling data provide a good overview of cell wall polymer distribution in *Marchantia*. Immunolocalization approaches using cell wall epitope directed antibodies or carbohydrate binding modules have previously shown that most polysaccharides like xyloglucan, homogalacturonan, RG-I, mannan, xylans and AGPs are somewhat conserved in *Marchantia* (Happ & Classen, 2019; Henry et al., 2020; Kolkas et al., 2023; Dierschke et al., 2024; summarised in Table 4). Our immunolabelling results further corroborate that *Marchantia* indeed has the cell wall polysaccharide structures comparable to that of other land plants with a ubiquitous deposition of cellulose, xyloglucan and mannan, a targeted deposition of the pectic epitope arabinan in the gametangiophore pegged rhizoids, the sporophyte foot, capsule and elaters. Extensin and MLG antibodies were also used to probe the different structures of *Marchantia*, but no signal was detected (Table 4). However, this does not preclude the presence of these epitopes in *Marchantia*, but perhaps indicate structural differences in the cell wall that hinder extensin detection or specifically in epitopes related to extensins.

**Table 4.** Summary of the antibodies assayed in Marchantia polymorpha, ^1^ This study; Several tissue types; Immunofluorescence, ^2^ Happ and Classen (2019); AGP extract; Enzyme-linked immunosorbent assay (ELISA), ^3^ Kolkas et al., (2023); thallus cell wall extracts; Immunodot-blots, ^4^ Henry et al., (2020); Placental transfer cells of the sporophyte or gametophyte; Immunogold labelling, ^5^Dierschke et al., (2024); Sporophytes; Immunofluorescence.

| Antibody | Target epitope | Tissue/polysaccharide, detected (+) or not detected (-) <sup>reference</sup> |
| --- | --- | --- |
| LM1 | Extensin | Thallus (-)<sup>1</sup>, stalk (-)<sup>1</sup>, antheridiophore (-)<sup>1</sup>, archegoniophore (-)<sup>1</sup> |
| LM2 | AGP β-GlcA | AGPs (+)<sup>2</sup>, Sporophyte (+)<sup>4</sup>, gametophyte (-)<sup>4</sup> |
| LM6 | α-1,5-arabinan | Rhizoids (+)<sup>1</sup>, elaters (+)<sup>1</sup>, AGPs (-)<sup>2</sup>, thallus (+)<sup>3</sup>, sporophyte (-)<sup>4</sup>, gametophyte (-)<sup>4</sup> |
| JIM7 | Partially methylesterified homogalacturonan | Sporophyte (+)<sup>4</sup>, gametophyte (+)<sup>4</sup> |
| JIM13 | AGP: βGlcA-1,3-α-GalA-1,2-Rha | AGPs (+)<sup>2</sup>, Sporophyte (+)<sup>4</sup>, gametophyte (+)<sup>4</sup> |
| KM1 | AGP | AGPs (+)<sup>2</sup> |
| LM16 | Galactosylated RG-I sidechain | Thallus (+)<sup>3</sup> |
| LM26 | β-(1,6)(1,4)-galactan | AGPs (TFA extracted) (+)<sup>2</sup> |
| LM14 | AGPs | AGPs (+)<sup>2</sup> |
| MAC207 | AGP: βGlcA-1,3-α-GalA-1,2-Rha | AGPs (-)<sup>2</sup> |
| LM19 | Unesterified homogalacturonan | Thallus (+)<sup>3</sup>, sporophyte (+)<sup>4</sup>, gametophyte (+)<sup>4</sup> |
| LM20 | Highly methylesterified homogalacturonan | Thallus (+)<sup>3</sup>, sporophyte (+)<sup>4</sup>, gametophyte (+)<sup>4</sup> |
| LM13 | Arabinan | Thallus (+)<sup>3</sup>, sporophyte (-)<sup>4</sup>, gametophyte (-)<sup>4</sup> |
| JIM5 | Partially methylesterified homogalacturonan | Sporophyte (+)<sup>4,5</sup>, gametophyte (+)<sup>4</sup> |
| LM25 | XLLG, XXLG, XXXG (sidechains of xyloglucan) | Thallus (+)<sup>3</sup>, sporophyte (+)<sup>4</sup>, gametophyte (+)<sup>4</sup> |
| LM24 | XXLG (sidechains of xyloglucan) | Thallus (+)<sup>3</sup> |
| LM15 | XXXG (sidechains of xyloglucan) | Thallus (+)<sup>3</sup>, thallus (+)<sup>1</sup>, stalk (+)<sup>1</sup>, antheridiophore (+)<sup>1</sup>, archegoniophore (+)<sup>1</sup>, |
| LM21 | Mannan | Thallus (+)<sup>3</sup>, sporophyte (+)<sup>4</sup>, gametophyte (+)<sup>4</sup> |
| LM22 | Mannan | Thallus (+)<sup>3</sup> |
| LM10 | Xylan | Thallus (+)<sup>3</sup> |
| LM11 | Xylan, arabinoxylan | Thallus (+)<sup>3</sup> |
| CBM3a | Crystalline cellulose | thallus (+)<sup>1</sup>, stalk (+)<sup>1</sup>, antheridiophore (+)<sup>1</sup>, archegoniophore (+)<sup>1</sup>, spore capsule (+)<sup>1,5</sup> |
| CBM4-1 | Amorphous cellulose | Sporophyte (+)<sup>5</sup> |
| LM5 | (1,4)-β-galactan | Sporophyte (+)<sup>4</sup>, gametophyte (+)<sup>4</sup> |
| LM28 | Xylogalacturonan | Sporophyte (-)<sup>4</sup>, gametophyte (-)<sup>4</sup> |
| JIM8 | AGP | Sporophyte (+)<sup>4</sup>, gametophyte (+)<sup>4</sup> |
| JIM12 | AGP | Sporophyte (-)<sup>4</sup>, gametophyte (-)<sup>4</sup> |
| anti-callose | Callose | Sporophyte (-)<sup>4</sup>, gametophyte (-)<sup>4</sup> |
| Aniline blue | Callose | Sporophyte (+)<sup>5</sup> |
| BS-400-3 | Mixed linkage β- glucan | thallus (-)<sup>1</sup>, stalk (-)<sup>1</sup>, antheridiophore (-)<sup>1</sup>, archegoniophore (-)<sup>1</sup> |
| NBGM C6 | β-1,4-mannan | thallus (+)<sup>1</sup>, stalk (+)<sup>1</sup>, antheridiophore (+)<sup>1</sup>, archegoniophore (+)<sup>1</sup>, spore capsule (+)<sup>1</sup> |

### Glycosyltransferase repertoire of *Marchantia*

To further investigate the distribution of *Marchantia* GT genes within each family, we conducted an Orthofinder analysis using whole proteomes of *A. thaliana* (Dicot), *B. distachyon* (Monocot), *S. moellendorffi* (Lycophyte), *M. polymorpha* (Liverwort), and *P. patens* (Moss), *C. braunii* (Charales), *Closterium* (Desmidiales) *M. endlicherianum* (Zygnematales) and *C. reinhardtii* (Chlorophyta) (Table S3). We compiled sequences for each of the 18-cell wall-related families using the *Arabidopsis* and *Marchantia* GT list on CAZy as reference (Table S3, Figure 6). It should be noted that the *Marchantia* CAZy GT list of the 18-cell wall-related families had two gene entries duplicated (Mp1g18380 and Mp3g22290), and one gene deleted in the latest version of the *Marchantia* genome (TAK-1 ver.7.1). In the *Arabidopsis* CAZy GT list of the 18-cell wall-related families, one gene entry was triplicated (At1g53040), three gene entries were duplicated (At2g02910, At4g20170 and At4g38500. Using the sequences, we constructed a phylogenetic tree for each cell wall-related GT family, which uncovered interesting insights into the putative cell wall biosynthetic machinery in *Marchantia*. It should be noted that splice variants were excluded from the count only where possible (*A. thaliana, B. distachyon, P. patens and M. polymorpha*), which may have led to an overestimation of the GT numbers in the other organisms. Furthermore, reciprocal blastp analysis (a two-way blastp showing reciprocal top hits for a *Marchantia* gene against representative chlorophyte, streptophyte, bryophyte and tracheophyte proteomes) was conducted to infer the ancestry of GT genes within the families of interest.

**Figure 6.**
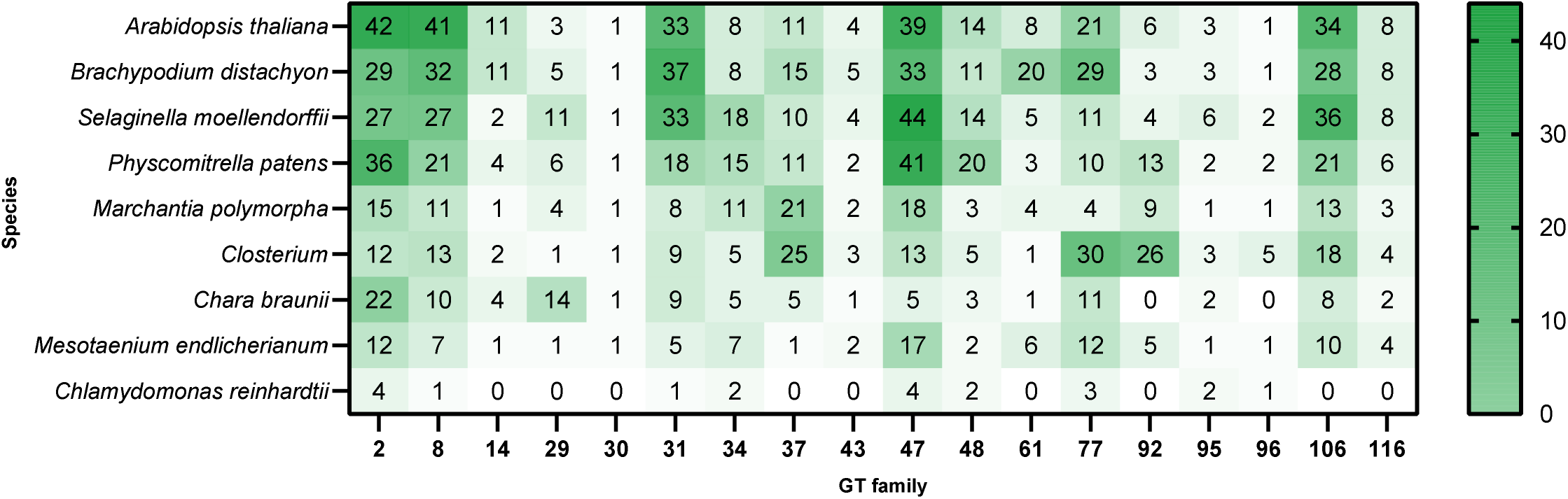
Number of genes in cell-wall related GT families in the species according to Orthofinder. Colour of the boxes correspond to the bar on the right-hand side.

Our Orthofinder analysis identified three more orthologs in GT family 37 of *Marchantia* which was not listed in CAZy (Mp6g02880, Mp6g02910 and Mp6g03040). For *Arabidopsis*, Orthofinder identified 17 additional orthologs across families GT8 (At1g03520 and At5g30500), GT14 (At1g53100, At1g71070, At2g37585, At3g03690, At3g15350, At3g24040, At4g03340, At4g27480, At5g15050 and At5g39990), GT37 (At2g15360), GT77 (At1g14600, At2g02061 and At2g42660) and GT92 (At3g60990). Thus, when we compare the GTs in the 18-cell wall-related families, *Marchantia* has 130, and *Arabidopsis* has 288 GTs. *Marchantia* has fewer members than *Arabidopsis* in most families including major families like GT2, GT8 and GT47. In GT families 29, 34, 37 and 92, *Marchantia* has more members than *Arabidopsis*, which may indicate novel catalytic activity not present in other land plant clades rather than gene duplication with conservation of function, given the overall low genetic redundancy in *Marchantia*.

### Cellulose

*Marchantia* only has two orthologs in the CELLULOSE SYNTHASE (CESA) subclade from GT family 2 (Figure S2). In *Arabidopsis*, CESAs operate in 18-subunit multimeric complexes (Cellulose Synthase Complex; CSC) (Nixon et al., 2016; Purushotham et al., 2020). The multimeric complex consists of equimolar amounts of three different CESA homologs, which are different between primary and secondary cell wall CSCs (Pedersen et al., 2023). In *P. patens*, there are seven CESA orthologs distributed in two subclades and both homo-oligomerisation or hetero-oligomerisation can take place (Li et al., 2022). Studies of the very few orthologs in *Marchantia* present in the CESA clade of GT family 2 may further provide insight into the basal form of the CSC in the land plant common ancestor (Lampugnani et al., 2019). Reciprocal blast shows that Mp7g07800 and Mp2g07690 showing streptophytic ancestry with relatively high support despite losses in some embryophyte lineages (Figure S3).

### Hemicelluloses

Members of the *CELLULOSE SYNTHASE-LIKE D* (*CSLD*) clade in GT family 2 exhibit β-1,4-mannan synthase and β-1,3-glucan synthase activity (Verhertbruggen et al., 2011; Yang et al., 2020). In our phylogenetic analysis, all *Marchantia* members as well as the orthologs in *P. patens* and *S. moellendorffi* in the *CSLD* subclade are closely related to *AtCSLD5*, a β-1,4-mannan synthase, and exhibit diversification within each species (Figure S2). A single *Marchantia* ortholog exists in the *CSLA* clade, where members exhibit β-1,4-mannan synthase activity (Dhugga et al., 2004; Goubet et al., 2009; Liepman et al., 2005; Voiniciuc et al., 2019). Reciprocal blast shows that Mp2g05640 and Mp5g24530 are respectively classified as having phragmoplastidae and post-phragmoplastidae ancestry, but have been lost in all other species except for Chara (evalue = 0) and Spirogolea (evalue = 0) (Table S4). Mp5g24550 and Mp5g24550 are of embryophyte ancestry with orthologue losses in some lineages (Figure S3). Mp2g05660 is classified as bryophyte-specific but not present in *Ricciocarpos*. Finally, Mp2g05670 and Mp2g02440 seem to be liverwort-specific (Figure S3).

The xyloglucan β-1,4-glucan synthases have been identified in *CELLULOSE SYNTHASE-LIKE C (CSLC)* clade in GT family 2 (Cocuron et al., 2007; S.-J. Kim et al., 2020). *Marchantia* has a single ortholog in this clade (Mp8g17530), which the reciprocal blast shows is robustly of viridiplantae ancestry (Figure S3). GT34 includes xyloglucan 1,6 xylosyltransferases (XXT1-5), where XXT1 and 2 preferentially catalyse the first two Xyl substitutions of XXGG and XXXG-type xyloglucans and XXT3-5 adds the third Xyl substitution onto XXXG-type xyloglucans (Culbertson et al., 2016; Ruprecht et al., 2018; N. Zhang et al., 2023). In GT34 and GT37, *Marchantia* and *P. patens* genes show extensive diversification (Figure S4, S5 respectively). The *At*XXT1 and 2 form a cluster distinct from *At*XXT3, 4 and 5. Consistent with only the XXGG branching type detected in *Marchantia* (Peña et al., 2008), the diversified members of *Marchantia* are more closely related to the clade containing *At*XXT1 and 2 (Figure S4). Within GT34 genes, Mp3g11650 is of viridiplantae origin with orthologues in chlorophytes and streptophytes. Mp3g03380 could be of viridiplantae origin as it is found in *Chlamydomonas* and *Ricciocarpos* but has been lost in all other lineages (Table S4, Figure S6). Mp8g11830 is of streptophytic origin although it’s been lost in tracheophytes, mosses and hornworts but has orthologues in *Klebsormidium* and *Chara*. Mp2g07920 dates back to the phragmoplastidae ancestor as it is robustly found in Chara, zygnematophytes and embryophytes. Mp2g00460 is found in *Ceratopteris* and *Ricciocarpos* suggesting embryophyte origin, and Mp5g04120, Mp3g22350, Mp3g03520, Mp3g03490, Mp2g18730 seem to have orthologues only in *Ricciocarpos*, suggesting liverwort or marchantophyta-origin.

The GT37 family contains xyloglucan α-1,2-fucosyltransferase (FUT1) which also has galactosyltransferase activity in *Arabidopsis* (Cicéron et al., 2016; Rocha et al., 2016; Urbanowicz et al., 2017). Interestingly, *P. patens* and *Marchantia* xyloglucans do not possess the fucosylated ‘F’ sidechain (Peña et al. 2008). Thus, it could be speculated that the dual function of FUT1 evolved after the split of the moss and liverwort common ancestor. On the other hand, *P. patens* and *Marchantia* xyloglucan have GalA sidechains structurally distinct from sidechains found in other land plant clades tested, including hornworts, monilophytes and lycopodiophytes (Peña et al. 2008). Thus, the diversified GTs of *Marchantia* family 37 pose as interesting candidates to explore whether they have also independently evolved a dual function mechanism, perhaps forming the linkages in GalA-containing xyloglucan sidechains. Consistently, Mp8g06690, Mp7g18170, Mp7g07390, Mp7g06430, Mp6g10590, Mp6g02870, Mp2g22100, Mp2g21110 and Mp2g06140 are likely liverwort-specific as reciprocal blast shows only orthologues in *Ricciocarpos* (Table S4). Meanwhile, Mp2g16920, Mp1g01370 and Mp2g20270 have respectively streptophytic, post-phragmoplastidae (common ancestor of zygnematophycean algae and land plants) and embryophyte ancestry (Figure S7). Mp4g10040 has an orthologue only in *Anthoceros* suggesting bryophyte ancestry. Mp7g07400 and Mp6g19640 have orthologues in *Physcomitrium* but losses in *Ricciocarpos* suggesting setaphyte ancestry. Finally, Mp2g12440 has clear orthologues in *Selaginella*, *Ricciocarpos* and *Anthoceros* with a *Chlamydomonas* gene showing reciprocity at a high evalue threshold (evalue = 0.000733), suggesting embryophyte ancestry but possibly viridiplantae ancestry with losses in streptophytic algae.

### Pectic polysaccharides

GT family 29 includes *At*GALT29A (At1g08280), a 1,3-galactan galactosyltransferase (Dilokpimol et al. 2014) and the RG-II cytidine 5’-monophospho-3-deoxy-D-manno-2-octulosonic acid (CMP-Kdo) transferase (*At*RCKT1; At1g08660) (Y. Zhang et al., 2024) (Figure S8). Phylogenetic analysis identified two putative *Marchantia* orthologs of *At*GALT29A, but none corresponding to *At*RCKT1. Instead, two genes, Mp5g13750 and Mp2g00070 form a distinct clade. Ancestry analysis based on reciprocal blasts, shows that Mp1g03590 is of streptophytic origin as putative orthologues are found in *Klebsormidium* and *Chara* but not in chlorophytes (Figure S9). Mp1g19260 is likely embryophyte-specific as no putative orthologues are found in chlorophytes and streptophytic algae. Mp5g13750 and Mp2g00070 are likely liverwort-specific as blastp only shows reciprocal top hits in *Ricciocarpos*.

GT family 92 includes *Arabidopsis* β-1,4-galactan synthases (GALS) that are also capable of adding terminal arabinofuranose (Liwanag et al., 2012; Ebert et al., 2018; Laursen et al., 2018). A clade with highly diversified *Marchantia* and moss sequences is present in addition to the clade containing the *Arabidopsis* GALS orthologs (Figure S10). Considering *Marchantia* has a relatively high cell wall Gal content, its members in GT92 poses an opportunity to probe for potentially new polysaccharide structures that are not present in angiosperms like *Arabidopsis*. Reciprocal blast suggests that Mp3g00230 and Mp1g21980 are of viridiplantae ancestry although their putative *Chlamydomonas*, *Klebsormidium* and *Chara* orthologues have evalues above 1xE-10 (Table S4, Figure S11). Mp6g19260 and Mp2g14970 may be of streptophyte origin although blastp evalues against *Klebsormidium* orthologues are respectively 6.09E-10 and 1.20E-09. Mp8g00660 is supported as of phragmoplastidae origin with Chara orthologue blastp evalues (1.07E-07) showing moderate support for this classification. Mp3g21560 and Mp2g21390 are highly supported as having post-phragmoplastidae origin with robust evalues against *Spirogolea* and other embryophyte proteomes but presenting losses in *Ceratopteris* and *Arabidopsis*. Mp7g09030 is likely of bryophyte origin with orthologues in *Anthoceros* and *Physcomitrium* but losses in *Ricciocarpos* (Figure S11). Finally Mp1g18380 seems to be of liverwort origin.

The phylogenetic analysis also allows insight into the evolution of RG-II. While *Marchantia* has orthologs of most characterised plant cell wall related GTs, orthologs of some RG-II biosynthetic genes have not yet been identified. Specifically, members of the GT family 29 clade containing the RG-II (Kdo) transferase (RCKT1) (Y. Zhang et al., 2024) (Figure S8), or the RG-II xylosyltransferases (RGXT) from GT family 77 (Egelund et al., 2006, 2008; Liu et al., 2011; Petersen et al., 2009) do not have orthologs in *Marchantia* (Table S3). This is somewhat puzzling as orthologs of RGXT and RCKT are present in *P. patens, C. braunii, Closterum* and *M. endlicherianum* (Table S3). Furthermore, *Marchantia* does not have orthologs of *At*CDI, the putative RG-II galactosyltransferase (Peng et al., 2021). Reciprocal blastp shows that Mp7g00150, Mp3g12220, and Mp1g13560 are robustly of Viridiplantae ancestry (Table S4, Figure S12). Meanwhile, Mp6g19260 and Mp2g14970 show borderline streptophytic ancestry with top hit evalues slightly above the 1xE-10 threshold in *Klebsormidium*. Mp8g00660 has a Chara orthologue with mild support (evalue = 1.07E-07) suggesting phragmosplastidae origin. Mp3g21560 and Mp2g21390 have clear orthologues in Spirogolea but possible losses in *Arabidopsis* and other embryophyte lineages, suggesting post-phragmoplastideae ancestry. Finally, Mp7g09030 and Mp1g18380 are of bryophyte and liverwort ancestry, respectively (Table S4, Figure S12).

RG-II is a heterogenous pectic polysaccharide characteristic with unique sidechains which include apiose, aceric acid, 3-deoxy-D-manno-octulosonic acid (Kdo) and 3-deoxy-D-lyxo-2-heptulosaric acid (Dha) on a 4-linked GalA backbone (Bar-Peled et al., 2012). The timing at which RG-II emerged during land plant evolution is unclear, but it is to be a land plant specific trait (Bar-Peled et al. 2012, Mikkelsen et al. 2014). Some charophycean green algae (CGA) contain Kdo and Dha either in their cell walls or associated with the cell surface (Becker et al. 1998, Domozych et al., 1991) and RG-II xylosyltransferase (RGXT) orthologs have also been identified in CGAs (Mikkelsen et al., 2014). However, whether the Kdo and Dha identified in these species is associated with RG-II, and whether the biochemical function is conserved is still unclear (Mikkelsen et al. 2014). The presence of RG-II in liverworts was speculated by the detection of, albeit in low amounts, borate-crosslinked RG-II (Matsunaga et al., 2004). Furthermore, putative orthologs of the UDP-Xylose/UDP-Apiose synthase (UAS; also called AXS) in *Arabidopsis* are present in bryophytes (Smith et al., 2016, Table S4). The overexpression of the duckweed AXS1/UAS1 in the moss *P. patens* led to an increase in apiose content in the metabolite fraction, but was not incorporated into the cell wall, leading the authors to speculate the lack of cell wall apiosyltransferases in bryophytes and algae (J. Smith et al., 2016). (Berardini et al., 2015)Indeed, the absence of orthologs in RG-II biosynthetic GTs in *Marchantia* further corroborate that RG-II may be absent, or structurally distinct to RG-II characterised in spermatophytes. Despite not having orthologs of the RG-II biosynthetic GTs identified thus far, other genes that are involved in RG-II substrate biosynthesis are present in *Marchantia*. For example, according to TAIR (https://www.arabidopsis.org/; Berardini et al., 2015) the GDP-D-mannose-4,6-dehydratase *MURUS 1* (*MUR1*) (Voxeur et al., 2017) has four orthologs in *Marchantia* and one ortholog of the *CMP-Kdo Synthase* (*CKS1*) (Kobayashi et al., 2011). Therefore, *Marchantia* may be useful for gain-of-function studies for RG-II biosynthetic GTs.

The reciprocal blastp analysis of *Marchantia* GTs reveal that the *Marchantia* genome has a GT repertoire that comprises ancestral streptophyte-specific, embryophyte-specific, and liverwort-specific proteins. The highest proportion of *Marchantia* GT-proteins show viridiplantae (N = 25/141), streptophytic (N=27), embryophytic (N = 21) and liverwort (N = 23) ancestry (Figure S13). It appears that GT-proteins have diversified across all major evolutionary transitions and they are not associated with a specific transition, perhaps suggesting the importance of consistently repurposing GT proteins to adapt to the changing environmental conditions.

### Building a *Marchantia* cell wall related GT cDNA library

Given the similarities in cell wall composition and GT inventories of *Marchantia* and other land plants, we argued that a comprehensive *Marchantia* GT compendium may be a valuable resource to explore GT functions and to accelerate plant cell wall research. To this end, we attempted to clone the coding sequences (CDS) of all annotated *Marchantia* GTs into Gateway compatible entry vectors pDONR221 or pENTR. The use of the Gateway technology allows efficient and high throughput transfer of DNA-fragments between plasmids (Katzen, 2007). All CDS sequences were amplified from mixed organ and developmental cDNA libraries using primers that placed the gene of interest in-frame with the *att* region and excluded the stop codon. Out of the 130 *Marchantia* GTs in cell-wall related families, we successfully generated 87 Gateway compatible entry vectors (Table S1). Using such an approach is advantageous as it allows entry vectors containing specific GOIs to be recombined with a range of destination vectors including those which could generate C-terminal translational fusions (Ishizaki et al., 2015). To exemplify the utility of the *Marchantia* cell wall GT library, YFP-fusion proteins were generated with entry clones harbouring the coding sequence for the two GTs from family 47 clade G (Mp3g24900 and Mp5g17760), which were subcloned into the pEARLEYGATE101 destination vector. The two members of GT47 clade G were chosen as they appear to lack orthologs in the angiosperm model organisms in our analysis. The constructs were transiently co-expressed with Golgi apparatus and ER organelle markers in *N. benthamiana.* As expected for type II GTs, *Mp*GT47G1 (Mp3g24900) and *Mp*GT47G2 (Mp5g17760.1) both exhibited punctate patterns that co-localised with the Golgi-marker (Figure 7).

**Figure 7.**
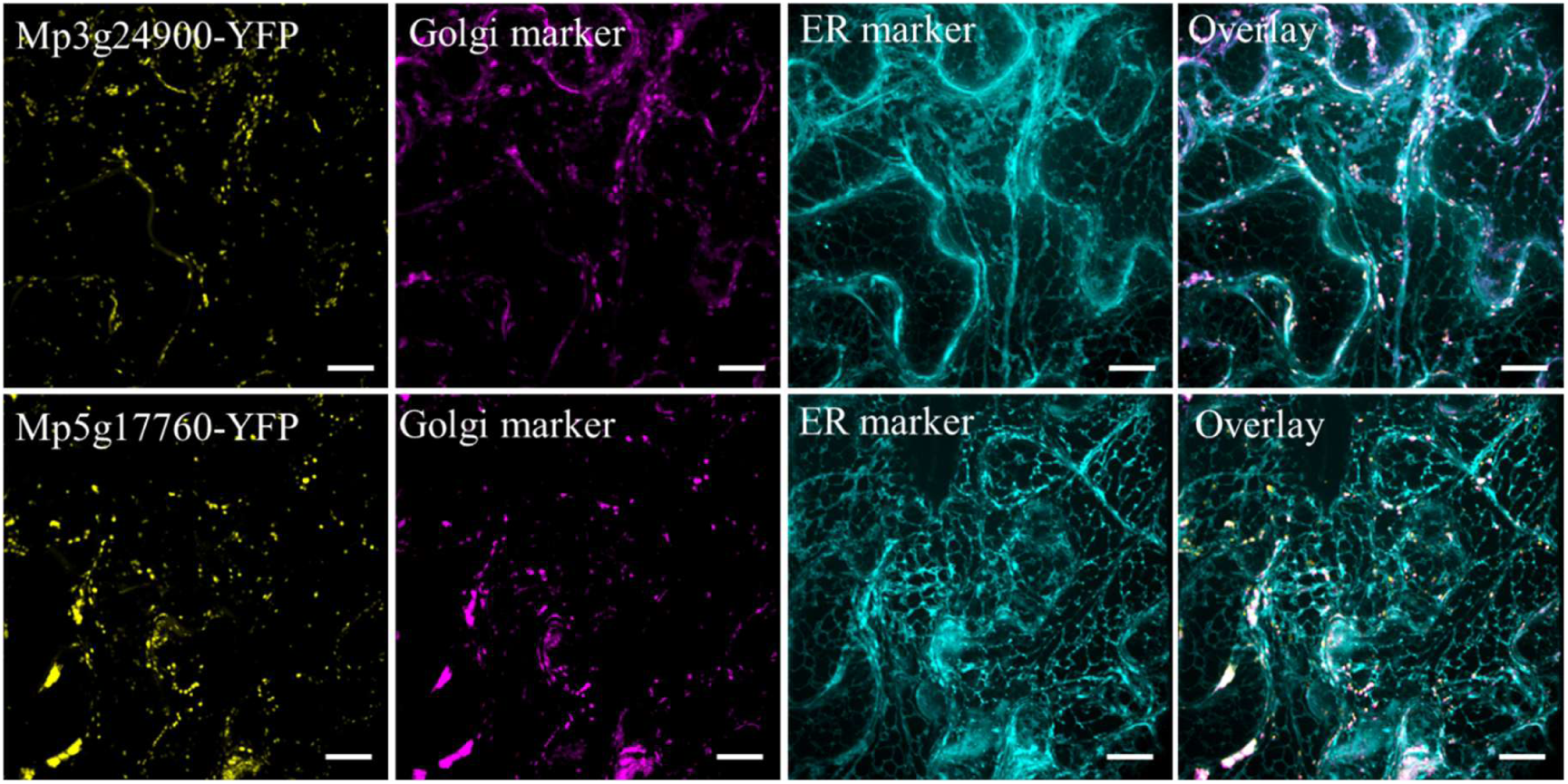
Subcellular co-localisation of Mp3g24900 and Mp5g17760.1 in *N. benthamiana* epidermal cells with Golgi- and ER-specific markers. Scale bar = 20 µm

In conjunction with the liverwort model system that can be transformed relatively easily and grown in the laboratory, the GT library has the potential to streamline and expedite the *in vivo* characterisation of cell wall related GTs. Further, it will be exciting to explore the members in GT families that are highly diversified in *Marchantia* compared to other land plants and GTs that are specific to *Marchantia*. Extending from GT studies, using *Marchantia* as a model organism to study plant cell wall biosynthesis and delivery could be useful considering the utility of plants also for glycoengineering of complex substrates like nucleotide sugars (Tang et al., 2023).

## Conclusion

*Marchantia* offers a simple genetic model system to study the complex mechanisms of cell wall biosynthesis. Here, we provide a detailed characterisation of the *Marchantia* cell wall across different tissue types. *Marchantia* has a cellulose-rich cell wall with mannan and xyloglucan also being abundant but a generally low level of pectic polysaccharides in mature stages. However, increased abundance of pectic polysaccharides is observed during early developmental stages. Phylogenetic analyses of cell wall related GTs in different species across the land plant lineage reveals conservation and diversification of genes in the *Marchantia* family, which may allude to novel linkages that are not found in other plants. With the development of a *Marchantia* cell wall GT cDNA library, we hope to further accelerate the identification and characterisation of GTs involved in cell wall assembly.

## Acknowledgements

This research was supported by The University of Melbourne’s Research Computing Services and the Petascale Campus Initiative. HSK was supported by the Australian Research Training Program (RTP) scholarship and the Ruhr University Bochum PhD Exchange Scholarship. ERL and HSK are grateful for support from the University of Melbourne Botany Foundation, Vassilios Sarafis research grant and The Australia & Pacific Science Foundation. ERL acknowledges funding from the Australian Academy of Science Thomas Davies Research. BE was supported by an Australian Research Council Discovery Project (DP180102630) and the 2020 Inaugural Botany Foundation Fellowship Award during this work. SP acknowledges a Villum Investigator (Project ID: 25915), DNRF Chair (DNRF155), Novo Nordisk Laureate (NNF19OC0056076), Novo Nordisk Emerging Investigator (NNF20OC0060564), and Novo Nordisk Data Science (NNF0068884) grants. JLB and EF-S were supported by funding from the Australian Research Council (CE200100015 to JLB). The authors wish to thank Dr Uli Felzmann, The University of Melbourne’s Research Computing Services and the Petascale Campus Initiative for assistance in accessing High Performance Computing (HPC) facilities. AB, CB and MSD acknowledge the support of La Trobe University to LISAF. AB and MSD acknowledge the support of the ARC Centre in Excellence for Plant Cell Walls (CE1101007).

## Supplementary Figure captions S2 – S13

Figure S2. Phylogenetic tree of GT family 2. Bootstrap values above 70 are indicated with lilac circles. Splice variants are shown in black. Labelled subclades are defined based on annotated function and OrthoFinder analysis (Table S3).

Figure S3. Reciprocal BLASTp analysis of *Marchantia* members in GT family 2. Support values represent the ratio of (sum_match/max_match), providing a measure for support for the ancestry classification (Table S4). The numbers within the tiles represent BLASTp e-values with -log10 transformation (Table S4).

Figure S4. Phylogenetic tree of GT family 34. Bootstrap values above 70 are indicated with lilac circles. Splice variants are shown in black. Labelled subclades are defined based on annotated function and OrthoFinder analysis (Table S3).

Figure S5. Phylogenetic tree of GT family 37. Bootstrap values above 70 are indicated with lilac circles. Splice variants are shown in black. Labelled subclades are defined based on annotated function and OrthoFinder analysis (Table S3).

Figure S6. Reciprocal BLASTp analysis of *Marchantia* members in GT family 34. Support values represent the ratio of (sum_match/max_match), providing a measure for support for the ancestry classification (Table S4). The numbers within the tiles represent BLASTp e-values with -log10 transformation (Table S4).

Figure S7. Reciprocal BLASTp analysis of *Marchantia* members in GT family 37. Support values represent the ratio of (sum_match/max_match), providing a measure for support for the ancestry classification (Table S4). The numbers within the tiles represent BLASTp e-values with -log10 transformation (Table S4).

Figure S8. Phylogenetic tree of GT family 29. Bootstrap values above 70 are indicated with lilac circles. Splice variants are shown in black. Labelled subclades are defined based on annotated function and OrthoFinder analysis (Table S3).

Figure S9. Reciprocal BLASTp analysis of *Marchantia* members in GT family 29. Support values represent the ratio of (sum_match/max_match), providing a measure for support for the ancestry classification (Table S4). The numbers within the tiles represent BLASTp e-values with -log10 transformation (Table S4).

Figure S10. Phylogenetic tree of GT family 92. Bootstrap values above 70 are indicated with lilac circles. Splice variants are shown in black. Labelled subclades are defined based on annotated function and OrthoFinder analysis (Table S3).

Figure S11. Reciprocal BLASTp analysis of *Marchantia* members in GT family 92. Support values represent the ratio of (sum_match/max_match), providing a measure for support for the ancestry classification (Table S4). The numbers within the tiles represent BLASTp e-values with -log10 transformation (Table S4).

Figure S12. Reciprocal BLASTp analysis of *Marchantia* members in GT family 77. Support values represent the ratio of (sum_match/max_match), providing a measure for support for the ancestry classification (Table S4). The numbers within the tiles represent BLASTp e-values with -log10 transformation (Table S4).

Figure S13. Count data from the reciprocal BLASTp analysis of *Marchantia* GTs. The largest proportion of *Marchantia* GT proteins show ancestry in Viridiplantae (N = 25/141), Streptophyta (N = 27), Embryophyta (N = 21), and liverworts (N = 23).

## References

1. Amos, R. A., Atmodjo, M. A., Huang, C., Gao, Z., Venkat, A., Taujale, R., Kannan, N., Moremen, K. W., & Mohnen, D. (2022). Polymerization of the backbone of the pectic polysaccharide rhamnogalacturonan I. Nature Plants, 8(11), 1289–1303. 10.1038/s41477-022-01270-3

2. Amos, R. A., Pattathil, S., Yang, J. Y., Atmodjo, M. A., Urbanowicz, B. R., Moremen, K. W., & Mohnen, D. (2018). A two-phase model for the non-processive biosynthesis of homogalacturonan polysaccharides by the GAUT1:GAUT7 complex. Journal of Biological Chemistry, 293(49), 19047–19063. 10.1074/jbc.RA118.004463

3. Anders, N., Wilkinson, M. D., Lovegrove, A., Freeman, J., Tryfona, T., Pellny, T. K., Weimar, T., Mortimer, J. C., Stott, K., Baker, J. M., Defoin-Platel, M., Shewry, P. R., Dupree, P., & Mitchell, R. A. C. (2012). Glycosyl transferases in family 61 mediate arabinofuranosyl transfer onto xylan in grasses. Proceedings of the National Academy of Sciences of the United States of America, 109(3), 989–993. 10.1073/pnas.1115858109

4. Anderson, C. T., Carroll, A., Akhmetova, L., & Somerville, C. (2010). Real-time imaging of cellulose reorientation during cell wall expansion in Arabidopsis roots. Plant Physiology, 152(2), 787–796. 10.1104/pp.109.150128

5. Atmodjo, M. A., Hao, Z., & Mohnen, D. (2013). Evolving Views of Pectin Biosynthesis. Annual Review of Plant Biology, 64(1), 747–779. 10.1146/annurev-arplant-042811-105534

6. Atmodjo, M. A., Sakuragi, Y., Zhu, X., Burrell, A. J., Mohanty, S. S., Atwood, J. A., Orlando, R., Scheller, H. V., & Mohnen, D. (2011). Galacturonosyltransferase (GAUT)1 and GAUT7 are the core of a plant cell wall pectin biosynthetic homogalacturonan:galacturonosyltransferase complex. Proceedings of the National Academy of Sciences of the United States of America, 108(50), 20225–20230. 10.1073/pnas.1112816108

7. Bacete, L., Mélida, H., Miedes, E., & Molina, A. (2018). Plant cell wall-mediated immunity: Cell wall changes trigger disease resistance responses. The Plant Journal, 93(4), 614–636. 10.1111/tpj.13807

8. Banks, J. A., Nishiyama, T., Hasebe, M., Bowman, J. L., Gribskov, M., dePamphilis, C., Albert, V. A., Aono, N., Aoyama, T., Ambrose, B. A., Ashton, N. W., Axtell, M. J., Barker, E., Barker, M. S., Bennetzen, J. L., Bonawitz, N. D., Chapple, C., Cheng, C., Correa, L. G. G., … Grigoriev, I. V. (2011). The compact Selaginella genome identifies changes in gene content associated with the evolution of vascular plants. Science (New York, N.Y.), 332(6032), 960–963. 10.1126/science.1203810

9. Bar-On, Y. M., Phillips, R., & Milo, R. (2018). The biomass distribution on Earth. Proceedings of the National Academy of Sciences, 115(25), 6506–6511. 10.1073/pnas.1711842115

10. Basu, D., Liang, Y., Liu, X., Himmeldirk, K., Faik, A., Kieliszewski, M., Held, M., & Showalter, A. M. (2013). Functional Identification of a Hydroxyproline-O-galactosyltransferase Specific for Arabinogalactan Protein Biosynthesis in Arabidopsis*. Journal of Biological Chemistry, 288(14), 10132–10143. 10.1074/jbc.M112.432609

11. Basu, D., Tian, L., Wang, W., Bobbs, S., Herock, H., Travers, A., & Showalter, A. M. (2015). A small multigene hydroxyproline-O-galactosyltransferase family functions in arabinogalactan-protein glycosylation, growth and development in Arabidopsis. BMC Plant Biology, 15(1), 295. 10.1186/s12870-015-0670-7

12. Basu, D., Wang, W., Ma, S., DeBrosse, T., Poirier, E., Emch, K., Soukup, E., Tian, L., & Showalter, A. M. (2015). Two Hydroxyproline Galactosyltransferases, GALT5 and GALT2, Function in Arabinogalactan-Protein Glycosylation, Growth and Development in Arabidopsis. PLOS ONE, 10(5), e0125624. 10.1371/journal.pone.0125624

13. Berardini, T. Z., Reiser, L., Li, D., Mezheritsky, Y., Muller, R., Strait, E., & Huala, E. (2015). The arabidopsis information resource: Making and mining the “gold standard” annotated reference plant genome. Genesis, 53(8), 474–485. 10.1002/dvg.22877

14. Biswal, A. K., Atmodjo, M. A., Pattathil, S., Amos, R. A., Yang, X., Winkeler, K., Collins, C., Mohanty, S. S., Ryno, D., Tan, L., Gelineo-Albersheim, I., Hunt, K., Sykes, R. W., Turner, G. B., Ziebell, A., Davis, M. F., Decker, S. R., Hahn, M. G., & Mohnen, D. (2018). Working towards recalcitrance mechanisms: Increased xylan and homogalacturonan production by overexpression of GAlactUronosylTransferase12 (GAUT12) causes increased recalcitrance and decreased growth in Populus. Biotechnology for Biofuels, 11(1), 9. 10.1186/s13068-017-1002-y

15. Bowman, J. L. (2022). Chapter One—The liverwort Marchantia polymorpha, a model for all ages. In B. Goldstein & M. Srivastava (Eds.), Current Topics in Developmental Biology (Vol. 147, pp. 1–32). Academic Press. 10.1016/bs.ctdb.2021.12.009

16. Bowman, J. L., Takayuki, K., & Yamato, K. T. (2017). Insights into Land Plant Evolution Garnered from the Marchantia polymorpha Genome. Cell, 171, 287–304.

17. Burton, R. A., Wilson, S. M., Hrmova, M., Harvey, A. J., Shirley, N. J., Medhurst, A., Stone, B. A., Newbigin, E. J., Bacic, A., & Fincher, G. B. (2006). Cellulose Synthase-Like CslF Genes Mediate the Synthesis of Cell Wall (1,3;1,4)-ß-d-Glucans. Science, 311(5769), 1940–1942. 10.1126/science.1122975

18. Cao, J., Dai, X., Zou, H., & Wang, Q. (2019). Formation and development of rhizoids of the liverwort Marchantía polymorpha Author (s): Jian-Guo Cao, Xi-Ling Dai, Hong-Mei Zou and Quan-Xi Wang Source: The Journal of the Torrey Botanical Society, Vol. 141, No. 2 (APRIL – JUNE 2014), Publishe. 141(2), 126–134.

19. Carella, P., Gogleva, A., Hoey, D. J., Bridgen, A. J., Stolze, S. C., Nakagami, H., & Schornack, S. (2019). Conserved Biochemical Defenses Underpin Host Responses to Oomycete Infection in an Early-Divergent Land Plant Lineage. Current Biology, 29(14), 2282–2294.e5. 10.1016/j.cub.2019.05.078

20. Carpita, N. C., & Gibeaut, D. M. (1993). Structural models of primary cell walls in flowering plants: Consistency of molecular structure with the physical properties of the walls during growth. Plant Journal, 3(1), 1–30. 10.1111/j.1365-313X.1993.tb00007.x

21. Carroll, S., Amsbury, S., Durney, C. H., Smith, R. S., Morris, R. J., Gray, J. E., & Fleming, A. J. (2022). Altering arabinans increases Arabidopsis guard cell flexibility and stomatal opening. Current Biology, 32(14), 3170–3179.e4. 10.1016/j.cub.2022.05.042

22. Cavalier, D. M., & Keegstra, K. (2006). Two Xyloglucan Xylosyltransferases Catalyze the Addition of Multiple Xylosyl Residues to Cellohexaose *. Journal of Biological Chemistry, 281(45), 34197–34207. 10.1074/jbc.M606379200

23. Cheng, S., Xian, W., Fu, Y., Marin, B., Keller, J., Wu, T., Sun, W., Li, X., Xu, Y., Zhang, Y., Wittek, S., Reder, T., Günther, G., Gontcharov, A., Wang, S., Li, L., Liu, X., Wang, J., Yang, H., … Melkonian, M. (2019). Genomes of Subaerial Zygnematophyceae Provide Insights into Land Plant Evolution. Cell, 179(5), 1057–1067.e14. 10.1016/j.cell.2019.10.019

24. Chiniquy, D., Sharma, V., Schultink, A., Baidoo, E. E., Rautengarten, C., Cheng, K., Carroll, A., Ulvskov, P., Harholt, J., Keasling, J. D., Pauly, M., Scheller, H. V., & Ronald, P. C. (2012). XAX1 from glycosyltransferase family 61 mediates xylosyltransfer to rice xylan. Proceedings of the National Academy of Sciences, 109(42), 17117–17122. 10.1073/pnas.1202079109

25. Cho, S. H., Du, J., Sines, I., Poosarla, V. G., Vepachedu, V., Kafle, K., Park, Y. B., Kim, S. H., Kumar, M., & Nixon, B. T. (2015). In vitro synthesis of cellulose microfibrils by a membrane protein from protoplasts of the non-vascular plant Physcomitrella patens. Biochemical Journal, 470(2), 195–205. 10.1042/BJ20141391

26. Cicéron, F., Rocha, J., Kousar, S., Hansen, S. F., Chazalet, V., Gillon, E., Breton, C., & Lerouxel, O. (2016). Expression, purification and biochemical characterization of AtFUT1, a xyloglucan-specific fucosyltransferase from *Arabidopsis thaliana*. Biochimie, 128–129, 183–192. 10.1016/j.biochi.2016.08.012

27. Clarke, A. E., Anderson, R. L., & Stone, B. A. (1979). Form and function of arabinogalactans and arabinogalactan-proteins. Phytochemistry, 18(4), 521–540. 10.1016/S0031-9422(00)84255-7

28. Cocuron, J.-C., Lerouxel, O., Drakakaki, G., Alonso, A. P., Liepman, A. H., Keegstra, K., Raikhel, N., & Wilkerson, C. G. (2007). A gene from the cellulose synthase-like C family encodes a β-1,4 glucan synthase. Proceedings of the National Academy of Sciences, 104(20), 8550–8555. 10.1073/pnas.0703133104

29. Cosgrove, D. J. (2016). Plant cell wall extensibility: Connecting plant cell growth with cell wall structure, mechanics, and the action of wall-modifying enzymes. Journal of Experimental Botany, 67(2), 463–476. 10.1093/jxb/erv511

30. Craig, R. J., Gallaher, S. D., Shu, S., Salomé, P. A., Jenkins, J. W., Blaby-Haas, C. E., Purvine, S. O., O’Donnell, S., Barry, K., Grimwood, J., Strenkert, D., Kropat, J., Daum, C., Yoshinaga, Y., Goodstein, D. M., Vallon, O., Schmutz, J., & Merchant, S. S. (2023). The Chlamydomonas Genome Project, version 6: Reference assemblies for mating-type plus and minus strains reveal extensive structural mutation in the laboratory. The Plant Cell, 35(2), 644–672. 10.1093/plcell/koac347

31. Culbertson, A. T., Chou, Y.-H., Smith, A. L., Young, Z. T., Tietze, A. A., Cottaz, S., Fauré, R., & Zabotina, O. A. (2016). Enzymatic Activity of Xyloglucan Xylosyltransferase 5. Plant Physiology, 171(3), 1893–1904. 10.1104/pp.16.00361

32. Curry, T. M., Peña, M. J., & Urbanowicz, B. R. (2023). An update on xylan structure, biosynthesis, and potential commercial applications. The Cell Surface, 9, 100101. 10.1016/j.tcsw.2023.100101

33. Delmas, F., Séveno, M., Northey, J. G. B., Hernould, M., Lerouge, P., McCourt, P., & Chevalier, C. (2008). The synthesis of the rhamnogalacturonan II component 3-deoxy-D-manno-2-octulosonic acid (Kdo) is required for pollen tube growth and elongation. Journal of Experimental Botany, 59(10), 2639–2647. 10.1093/jxb/ern118

34. Dhugga, K. S., Barreiro, R., Whitten, B., Stecca, K., Hazebroek, J., Randhawa, G. S., Dolan, M., Kinney, A. J., Tomes, D., Nichols, S., & Anderson, P. (2004). Guar Seed b-Mannan Synthase Is a Member of the Cellulose Synthase Super Gene Family. Science, 303(5656), 363–366.

35. Dierschke, T., Levins, J., Lampugnani, E. R., Ebert, B., Zachgo, S., & Bowman, J. L. (2024). Control of sporophyte secondary cell wall development in Marchantia by a Class II KNOX gene. *Current Biology*, S0960982224013290. 10.1016/j.cub.2024.09.061

36. Dilokpimol, A., & Geshi, N. (2014). Arabidopsis thaliana glucuronosyltransferase in family GT14. Plant Signaling & Behavior, 9, e28891. 10.4161/psb.28891

37. Dilokpimol, A., Poulsen, C. P., Vereb, G., Kaneko, S., Schulz, A., & Geshi, N. (2014). Galactosyltransferases from arabidopsis thaliana in the biosynthesis of type II arabinogalactan: Molecular interaction enhances enzyme activity. BMC Plant Biology, 14(1), 1–14. 10.1186/1471-2229-14-90

38. Doblin, M. S., Ma, Y., Novaković, L., Bacic, A., & Johnson, K. L. (2023). Plant Cell Walls – Past, Present, and Future. In A. Geitmann, Plant Cell Walls (1st ed., pp. 1–28). CRC Press. 10.1201/9781003178309-1

39. Doblin, M. S., Pettolino, F. A., Wilson, S. M., Campbell, R., Burton, R. A., Fincher, G. B., Newbigin, E., & Bacic, A. (2009). A barley cellulose synthase-like CSLH gene mediates (1,3;1,4)-β-d-glucan synthesis in transgenic Arabidopsis. Proceedings of the National Academy of Sciences, 106(14), 5996–6001. 10.1073/pnas.0902019106

40. Drula, E., Garron, M.-L., Dogan, S., Lombard, V., Henrissat, B., & Terrapon, N. (2022). The carbohydrate-active enzyme database: Functions and literature. Nucleic Acids Research, 50(D1), D571–D577. 10.1093/nar/gkab1045

41. Ebert, B., Birdseye, D., Liwanag, A. J. M., Laursen, T., Rennie, E. A., Guo, X., Catena, M., Rautengarten, C., Stonebloom, S. H., Gluza, P., Pidatala, V. R., Andersen, M. C. F., Cheetamun, R., Mortimer, J. C., Heazlewood, J. L., Bacic, A., Clausen, M. H., Willats, W. G. T., & Scheller, H. V. (2018). The three members of the Arabidopsis Glycosyltransferase Family 92 are functional B-1,4-galactan synthases. Plant and Cell Physiology, 59(12), 2624–2636. 10.1093/pcp/pcy180

42. Edwards, D., Richardsont, D. B. J. B., & Boron, W. F. (1994). Hepatic characters in the earliest land plants Out-of-equilibrium C02lHCO ; solutions and their use in characterizing a new K / HC0 3 cotransporter. In Situ, 3540(1987), 1994–1995.

43. Egelund, J., Damager, I., Faber, K., Olsen, C.-E., Ulvskov, P., & Petersen, B. L. (2008). Functional characterisation of a putative rhamnogalacturonan II specific xylosyltransferase. FEBS Letters, 582(21–22), 3217–3222. 10.1016/j.febslet.2008.08.015

44. Egelund, J., Petersen, B. L., Motawia, M. S., Damager, I., Faik, A., Olsen, C. E., Ishii, T., Clausen, H., Ulvskov, P., & Geshi, N. (2006). Arabidopsis thaliana RGXT1 and RGXT2 Encode Golgi-Localized (1,3)-α-d-Xylosyltransferases Involved in the Synthesis of Pectic Rhamnogalacturonan-II. The Plant Cell, 18(10), 2593–2607. 10.1105/tpc.105.036566

45. Emms, D. M., & Kelly, S. (2019). OrthoFinder: Phylogenetic orthology inference for comparative genomics. Genome Biology, 20(1), 238. 10.1186/s13059-019-1832-y

46. Espiñeira, J. M., Novo Uzal, E., Gómez Ros, L. V., Carrión, J. S., Merino, F., Ros Barceló, A., & Pomar, F. (2011). Distribution of lignin monomers and the evolution of lignification among lower plants. Plant Biology, 13(1), 59–68. 10.1111/j.1438-8677.2010.00345.x

47. Faik, A., Bar-Peled, M., DeRocher, A. E., Zeng, W., Perrin, R. M., Wilkerson, C., Raikhel, N. V., & Keegstra, K. (2000). Biochemical characterization and molecular cloning of an alpha-1,2-fucosyltransferase that catalyzes the last step of cell wall xyloglucan biosynthesis in pea. The Journal of Biological Chemistry, 275(20), 15082–15089. 10.1074/jbc.M000677200

48. Faik, A., Price, N. J., Raikhel, N. V., & Keegstra, K. (2002). An Arabidopsis gene encoding an α-xylosyltransferase involved in xyloglucan biosynthesis. Proceedings of the National Academy of Sciences, 99(11), 7797–7802. 10.1073/pnas.102644799

49. Feijao, C., Morreel, K., Anders, N., Tryfona, T., Busse-Wicher, M., Kotake, T., Boerjan, W., & Dupree, P. (2022). Hydroxycinnamic acid-modified xylan side chains and their cross-linking products in rice cell walls are reduced in the Xylosyl arabinosyl substitution of xylan 1 mutant. The Plant Journal: For Cell and Molecular Biology, 109(5), 1152– 1167. 10.1111/tpj.15620

50. Geshi, N., Johansen, J. N., Dilokpimol, A., Rolland, A., Belcram, K., Verger, S., Kotake, T., Tsumuraya, Y., Kaneko, S., Tryfona, T., Dupree, P., Scheller, H. V., Höfte, H., & Mouille, G. (2013). A galactosyltransferase acting on arabinogalactan protein glycans is essential for embryo development in Arabidopsis. The Plant Journal, 76(1), 128–137. 10.1111/tpj.12281

51. Gille, S., Sharma, V., Baidoo, E. E. K., Keasling, J. D., Scheller, H. V., & Pauly, M. (2013). Arabinosylation of a Yariv-precipitable cell wall polymer impacts plant growth as exemplified by the arabidopsis glycosyltransferase mutant ray1. Molecular Plant, 6(4), 1369–1372. 10.1093/mp/sst029

52. Goubet, F., Barton, C. J., Mortimer, J. C., Yu, X., Zhang, Z., Miles, G. P., Richens, J., Liepman, A. H., Seffen, K., & Dupree, P. (2009). Cell wall glucomannan in Arabidopsis is synthesised by CSLA glycosyltransferases, and influences the progression of embryogenesis. The Plant Journal, 60(3), 527–538. 10.1111/j.1365-313X.2009.03977.x

53. Happ, K., & Classen, B. (2019). Arabinogalactan-proteins from the liverwort marchantia polymorpha L., a member of a basal land plant lineage, are structurally different to those of angiosperms. Plants, 8(11). 10.3390/plants8110460

54. Harholt, J., Jensen. J.K, Sørensen, S., O., Orfila, C., Pauly, M., & Scheller, H. V. (2006). ARABINAN DEFICIENT 1 Is a Putative Arabinosyltransferase Involved in Biosynthesis of Pectic Arabinan in Arabidopsis. Plant Physiology, 140(1), 49–58. 10.1104/pp.105.072744

55. Henry, J. S., Lopez, R. A., & Renzaglia, K. S. (2020). Differential localization of cell wall polymers across generations in the placenta of Marchantia polymorpha. Journal of Plant Research, 133(6), 911–924. 10.1007/s10265-020-01232-w

56. Hernandez-Gomez, M. C., Rydahl, M. G., Rogowski, A., Morland, C., Cartmell, A., Crouch, L., Labourel, A., Fontes, C. M. G. A., Willats, W. G. T., Gilbert, H. J., & Knox, J. P. (2015). Recognition of xyloglucan by the crystalline cellulose-binding site of a family 3a carbohydrate-binding module. FEBS Letters, 589(18), 2297–2303. 10.1016/j.febslet.2015.07.009

57. Hinz, S. W. A., Verhoef, R., Schols, H. A., Vincken, J. P., & Voragen, A. G. J. (2005). Type I arabinogalactan contains β-D-Galp-(1→3)-β-D-Galp structural elements. Carbohydrate Research, 340(13), 2135–2143. 10.1016/j.carres.2005.07.003

58. Hoch, H. C., Galvani, C. D., Szarowski, D. H., & Turner, J. N. (2005). Two new fluorescent dyes applicable for visualization of fungal cell walls. Mycologia, 97(3), 580–588. 10.3852/mycologia.97.3.580

59. Houston, K., Tucker, M. R., Chowdhury, J., Shirley, N., & Little, A. (2016). The Plant Cell Wall: A Complex and Dynamic Structure As Revealed by the Responses of Genes under Stress Conditions. Frontiers in Plant Science, 7, 984. 10.3389/fpls.2016.00984

60. Humphreys, C. P., Franks, P. J., Rees, M., Bidartondo, M. I., Leake, J. R., & Beerling, D. J. (2010). Mutualistic mycorrhiza-like symbiosis in the most ancient group of land plants. Nature Communications, 1(1), 103. 10.1038/ncomms1105

61. Ishizaki, K., Nishihama, R., Ueda, M., Inoue, K., Ishida, S., Nishimura, Y., Shikanai, T., & Kohchi, T. (2015). Development of gateway binary vector series with four different selection markers for the liverwort marchantia polymorpha. PLoS ONE, 10(9), 1–13. 10.1371/journal.pone.0138876

62. Ishizaki, K., Nishihama, R., Yamato, K. T., & Kohchi, T. (2016). Molecular genetic tools and techniques for marchantia polymorpha research. Plant and Cell Physiology, 57(2). 10.1093/pcp/pcv097

63. Jensen, J. K., Busse-Wicher, M., Poulsen, C. P., Fangel, J. U., Smith, P. J., Yang, J.-Y., Peña, M.-J., Dinesen, M. H., Martens, H. J., Melkonian, M., Wong, G. K.-S., Moremen, K. W., Wilkerson, C. G., Scheller, H. V., Dupree, P., Ulvskov, P., Urbanowicz, B. R., & Harholt, J. (2018). Identification of an algal xylan synthase indicates that there is functional orthology between algal and plant cell wall biosynthesis. New Phytologist, 218(3), 1049–1060. 10.1111/nph.15050

64. Jensen, J. K., Johnson, N. R., & Wilkerson, C. G. (2014). Arabidopsis thaliana IRX10 and two related proteins from psyllium and Physcomitrella patens are xylan xylosyltransferases. The Plant Journal, 80(2), 207–215. 10.1111/tpj.12641

65. Jensen, J. K., Schultink, A., Keegstra, K., Wilkerson, C. G., & Pauly, M. (2012). RNA-seq analysis of developing nasturtium seeds (Tropaeolum majus): Identification and characterization of an additional galactosyltransferase involved in xyloglucan biosynthesis. Molecular Plant, 5(5), 984–992. 10.1093/mp/sss032

66. Jensen, J. K., Sørensen, S. O., Harholt, J., Geshi, N., Sakuragi, Y., Møller, I., Zandleven, J., Bernal, A. J., Jensen, N. B., Sørensen, C., Pauly, M., Beldman, G., Willats, W. G. T., & Scheller, H. V. (2008). Identification of a Xylogalacturonan Xylosyltransferase Involved in Pectin Biosynthesis in Arabidopsis. The Plant Cell, 20(5), 1289–1302. 10.1105/tpc.107.050906

67. Jiang, N., Wiemels, R. E., Soya, A., Whitley, R., Held, M., & Faik, A. (2016). Composition, Assembly, and Trafficking of a Wheat Xylan Synthase Complex. Plant Physiology, 170(4), 1999–2023. 10.1104/pp.15.01777

68. Jibran, R., Hill, S. J., Lampugnani, E. R., Hao, P., Doblin, M. S., Bacic, A., Vaidya, A. A., O’Donoghue, E. M., McGhie, T. K., Albert, N. W., Zhou, Y., Raymond, L. G., Schwinn, K. E., Jordan, B. R., Bowman, J. L., Davies, K. M., & Brummell, D. A. (2024). The auronidin flavonoid pigments of the liverwort Marchantia polymorpha form polymers that modify cell wall properties. The Plant Journal, 120(3), 1159–1175. 10.1111/tpj.17045

69. Jobling, S. A. (2015). Membrane pore architecture of the CslF6 protein controls (1-3,1-4)-β-glucan structure. Science Advances. 10.1126/sciadv.1500069

70. Johnson, K. L., Gidley, M. J., Bacic, A., & Doblin, M. S. (2018). Cell wall biomechanics: A tractable challenge in manipulating plant cell walls ‘fit for purpose’! Current Opinion in Biotechnology, 49, 163–171. 10.1016/j.copbio.2017.08.013

71. Jones, L., Milne, J. L., Ashford, D., & McQueen-Mason, S. J. (2003). Cell wall arabinan is essential for guard cell function. Proceedings of the National Academy of Sciences, 100(20), 11783–11788. 10.1073/pnas.1832434100

72. Katzen, F. (2007). Gateway® recombinational cloning: A biological operating system. Expert Opinion on Drug Discovery, 2(4), 571–589. 10.1517/17460441.2.4.571

73. Kawaguchi, Y. W., Tsuchikane, Y., Tanaka, K., Taji, T., Suzuki, Y., Toyoda, A., Ito, M., Watano, Y., Nishiyama, T., Sekimoto, H., & Tsuchimatsu, T. (2023). Extensive Copy Number Variation Explains Genome Size Variation in the Unicellular Zygnematophycean Alga, Closterium peracerosum–strigosum–littorale Complex. Genome Biology and Evolution, 15(8), evad115. 10.1093/gbe/evad115

74. Kim, S. J., & Brandizzi, F. (2016). The plant secretory pathway for the trafficking of cell wall polysaccharides and glycoproteins. Glycobiology, 26(9), 940–949. 10.1093/glycob/cww044

75. Kim, S.-J., Chandrasekar, B., Rea, A. C., Danhof, L., Zemelis-Durfee, S., Thrower, N., Shepard, Z. S., Pauly, M., Brandizzi, F., & Keegstra, K. (2020). The synthesis of xyloglucan, an abundant plant cell wall polysaccharide, requires CSLC function. Proceedings of the National Academy of Sciences, 117(33), 20316–20324. 10.1073/pnas.2007245117

76. Kim, S.-J., Zemelis, S., Keegstra, K., & Brandizzi, F. (2015). The cytoplasmic localization of the catalytic site of CSLF6 supports a channeling model for the biosynthesis of mixed-linkage glucan. 10.1111/tpj.12748

77. Knoch, E., Dilokpimol, A., Tryfona, T., Poulsen, C. P., Xiong, G., Harholt, J., Petersen, B. L., Ulvskov, P., Hadi, M. Z., Kotake, T., Tsumuraya, Y., Pauly, M., Dupree, P., & Geshi, N. (2013). A β–glucuronosyltransferase from Arabidopsis thaliana involved in biosynthesis of type II arabinogalactan has a role in cell elongation during seedling growth. The Plant Journal, 76(6), 1016–1029. 10.1111/tpj.12353

78. Kobayashi, M., Kouzu, N., Inami, A., Toyooka, K., Konishi, Y., Matsuoka, K., & Matoh, T. (2011). Characterization of Arabidopsis CTP:3-Deoxy-d-manno-2-Octulosonate Cytidylyltransferase (CMP-KDO synthetase), the Enzyme that Activates KDO During Rhamnogalacturonan II Biosynthesis. Plant and Cell Physiology, 52(10), 1832–1843. 10.1093/pcp/pcr120

79. Kolkas, H., Burlat, V., & Jamet, E. (2023). Immunochemical Identification of the Main Cell Wall Polysaccharides of the Early Land Plant Marchantia polymorpha. Cells, 12(14), Article 14. 10.3390/cells12141833

80. Kremer, C., Pettolino, F., Bacic, A., & Drinnan, A. (2004). Distribution of cell wall components in Sphagnum hyaline cells and in liverwort and hornwort elaters. Planta, 219(6), 1023–1035. 10.1007/s00425-004-1308-4

81. Lairson, L. L., Henrissat, B., Davies, G. J., & Withers, S. G. (2008). Glycosyltransferases: Structures, Functions, and Mechanisms. Annual Review of Biochemistry, 77(1), 521–555. 10.1146/annurev.biochem.76.061005.092322

82. Lampugnani, E. R., Ho, Y. Y., Moller, I. E., Koh, P.-L., Golz, J. F., Bacic, A., & Newbigin, E. (2016). A Glycosyltransferase from Nicotiana alata Pollen Mediates Synthesis of a Linear (1,5)-α-L-Arabinan When Expressed in Arabidopsis. Plant Physiology, 170(4), 1962–1974. 10.1104/pp.15.02005

83. Lampugnani, E. R., Khan, G. A., Somssich, M., & Persson, S. (2018). Building a plant cell wall at a glance. Journal of Cell Science, 131(2), jcs207373–jcs207373. 10.1242/jcs.207373

84. Lang, D., Ullrich, K. K., Murat, F., Fuchs, J., Jenkins, J., Haas, F. B., Piednoel, M., Gundlach, H., Van Bel, M., Meyberg, R., Vives, C., Morata, J., Symeonidi, A., Hiss, M., Muchero, W., Kamisugi, Y., Saleh, O., Blanc, G., Decker, E. L., … Rensing, S. A. (2018). The Physcomitrella patens chromosome-scale assembly reveals moss genome structure and evolution. The Plant Journal, 93(3), 515–533. 10.1111/tpj.13801

85. Laursen, T., Stonebloom, S. H., Pidatala, V. R., Birdseye, D. S., Clausen, M. H., Mortimer, J. C., & Scheller, H. V. (2018). Bifunctional glycosyltransferases catalyze both extension and termination of pectic galactan oligosaccharides. Plant Journal, 94(2), 340–351. 10.1111/tpj.13860

86. Lee, C., Zhong, R., & Ye, Z.-H. (2012). Arabidopsis Family GT43 Members are Xylan Xylosyltransferases Required for the Elongation of the Xylan Backbone. Plant and Cell Physiology, 53(1), 135–143. 10.1093/pcp/pcr158

87. Lee, K. J. D., Sakata, Y., Mau, S.-L., Pettolino, F., Bacic, A., Quatrano, R. S., Knight, C. D., & Knox, J. P. (2005). Arabinogalactan Proteins Are Required for Apical Cell Extension in the Moss Physcomitrella patens. The Plant Cell Online, 17(11), 3051–3065. 10.1105/tpc.105.034413

88. Li, X., Chaves, A. M., Dees, D. C. T., Mansoori, N., Yuan, K., Speicher, T. L., Norris, J. H., Wallace, I. S., Trindade, L. M., & Roberts, A. W. (2022). Cellulose synthesis complexes are homo-oligomeric and hetero-oligomeric in Physcomitrium patens. Plant Physiology, 188(4), 2115–2130. 10.1093/plphys/kiac003

89. Liepman, A. H., Wilkerson, C. G., & Keegstra, K. (2005). Expression of cellulose synthase-like (Csl) genes in insect cells reveals that CslA family members encode mannan synthases. Proceedings of the National Academy of Sciences of the United States of America, 102(6), 2221–2226. 10.1073/pnas.0409179102

90. Little, A., Lahnstein, J., Jeffery, D. W., Khor, S. F., Schwerdt, J. G., Shirley, N. J., Hooi, M., Xing, X., Burton, R. A., & Bulone, V. (2019). A Novel (1,4)-β-Linked Glucoxylan Is Synthesized by Members of the Cellulose Synthase-Like F Gene Family in Land Plants. ACS Central Science. 10.1021/acscentsci.8b00568

91. Little, A., Schwerdt, J. G., Shirley, N. J., Khor, S. F., Neumann, K., O’Donovan, L. A., Lahnstein, J., Collins, H. M., Henderson, M., Fincher, G. B., & Burton, R. A. (2018). Revised Phylogeny of the Cellulose Synthase Gene Superfamily: Insights into Cell Wall Evolution1[OPEN]. Plant Physiology, 177(3), 1124–1141. 10.1104/pp.17.01718

92. Liu, X.-L., Liu, L., Niu, Q.-K., Xia, C., Yang, K.-Z., Li, R., Chen, L.-Q., Zhang, X.-Q., Zhou, Y., & Ye, D. (2011). MALE GAMETOPHYTE DEFECTIVE 4 encodes a rhamnogalacturonan II xylosyltransferase and is important for growth of pollen tubes and roots in Arabidopsis. The Plant Journal, 65(4), 647–660. 10.1111/j.1365-313X.2010.04452.x

93. Liwanag, A. J. M., Ebert, B., Verhertbruggen, Y., Rennie, E. A., Rautengarten, C., Oikawa, A., Andersen, M. C. F., Clausen, M. H., & Scheller, H. V. (2012). Pectin Biosynthesis: GALS1 in Arabidopsis thaliana Is a b -1, 4-Galactan b -1, 4-Galactosyltransferase. 24(December), 5024–5036. 10.1105/tpc.112.106625

94. Madson, M., Dunand, C., Li, X., Verma, R., Vanzin, G. F., Caplan, J., Shoue, D. A., Carpita, N. C., & Reiter, W.-D. (2003). The MUR3 gene of Arabidopsis encodes a xyloglucan galactosyltransferase that is evolutionarily related to animal exostosins. The Plant Cell, 15(7), 1662–1670. 10.1105/tpc.009837

95. Marcus, S. E., Verhertbruggen, Y., Hervé, C., Ordaz-Ortiz, J. J., Farkas, V., Pedersen, H. L., Willats, W. G., & Knox, J. P. (2008). Pectic homogalacturonan masks abundant sets of xyloglucan epitopes in plant cell walls. BMC Plant Biology, 8(1), 60. 10.1186/1471-2229-8-60

96. Mariette, A., Curry, T. M., Urbanowicz, B. R., & Ebert, B. (2023). Plant Cell Wall Glycosyltransferases: From Sequence to Structure and Function. In Plant Cell Walls. CRC Press.

97. Matsunaga, T., Ishii, T., Matsumoto, S., Higuchi, M., Darvill, A., Albersheim, P., & O’Neill, M. A. (2004). Occurrence of the Primary Cell Wall Polysaccharide Rhamnogalacturonan II in Pteridophytes, Lycophytes, and Bryophytes. Implications for the Evolution of Vascular Plants. Plant Physiology, 134(1), 339–351. 10.1104/pp.103.030072

98. McFarlane, H. E., Döring, A., & Persson, S. (2014). The Cell Biology of Cellulose Synthesis. Annual Review of Plant Biology, 65(1), 69–94. 10.1146/annurev-arplant-050213-040240

99. Mohnen, D. (2008). Pectin structure and biosynthesis. Current Opinion in Plant Biology, 11(3), 266–277. 10.1016/j.pbi.2008.03.006

100. Molina, A., Miedes, E., Bacete, L., Rodríguez, T., Mélida, H., Denancé, N., Sánchez-Vallet, A., Rivière, M.-P., López, G., Freydier, A., Barlet, X., Pattathil, S., Hahn, M., & Goffner, D. (2021). Arabidopsis cell wall composition determines disease resistance specificity and fitness. Proceedings of the National Academy of Sciences, 118(5), e2010243118. 10.1073/pnas.2010243118

101. Moller, I., Sørensen, I., Bernal, A. J., Blaukopf, C., Lee, K., Øbro, J., Pettolino, F., Roberts, A., Mikkelsen, J. D., Knox, J. P., Bacic, A., & Willats, W. G. T. (2007). High-throughput mapping of cell-wall polymers within and between plants using novel microarrays. The Plant Journal, 50(6), 1118–1128. 10.1111/j.1365-313X.2007.03114.x

102. Møller, S. R., Yi, X., Velásquez, S. M., Gille, S., Hansen, P. L. M., Poulsen, C. P., Olsen, C. E., Rejzek, M., Parsons, H., Zhang, Y., Wandall, H. H., Clausen, H., Field, R. A., Pauly, M., Estevez, J. M., Harholt, J., Ulvskov, P., & Petersen, B. L. (2017). Identification and evolution of a plant cell wall specific glycoprotein glycosyl transferase, ExAD. Scientific Reports, 7(March), 1–16. 10.1038/srep45341

103. Moore, J. P., Farrant, J. M., & Driouich, A. (2008). A role for pectin-associated arabinans in maintaining the flexibility of the plant cell wall during water deficit stress. Plant Signaling and Behavior, 3(2), 102–104. 10.4161/psb.3.2.4959

104. Moore, J. P., Nguema-Ona, E. E., Vicré-Gibouin, M., Sørensen, I., Willats, W. G. T., Driouich, A., & Farrant, J. M. (2013). Arabinose-rich polymers as an evolutionary strategy to plasticize resurrection plant cell walls against desiccation. Planta, 237(3), 739–754. 10.1007/s00425-012-1785-9

105. Moremen, K. W., & Haltiwanger, R. S. (2019). Emerging structural insights into glycosyltransferase-mediated synthesis of glycans. Nature Chemical Biology, 15(9), 853–864. 10.1038/s41589-019-0350-2

106. Mortimer, J. C., Faria-Blanc, N., Yu, X., Tryfona, T., Sorieul, M., Ng, Y. Z., Zhang, Z., Stott, K., Anders, N., & Dupree, P. (2015). An unusual xylan in Arabidopsis primary cell walls is synthesised by GUX3, IRX9L, IRX10L and IRX14. The Plant Journal, 83(3), 413–426. 10.1111/tpj.12898

107. Narciso, J. O., Zeng, W., Ford, K., Lampugnani, E. R., Humphries, J., Austarheim, I., van de Meene, A., Bacic, A., & Doblin, M. S. (2021). Biochemical and Functional Characterization of GALT8, an Arabidopsis GT31 β-(1,3)-Galactosyltransferase That Influences Seedling Development. Frontiers in Plant Science, 12. https://www.frontiersin.org/articles/10.3389/fpls.2021.678564

108. Nelson, B. K., Cai, X., & Nebenführ, A. (2007). A multicolored set of in vivo organelle markers for co-localization studies in Arabidopsis and other plants. Plant Journal, 51(6), 1126–1136. 10.1111/j.1365-313X.2007.03212.x

109. Nguema-Ona, E., Vicré-Gibouin, M., Gotté, M., Plancot, B., Lerouge, P., Bardor, M., & Driouich, A. (2014). Cell wall O-glycoproteins and N-glycoproteins: Aspects of biosynthesis and function. Frontiers in Plant Science, 5(OCT), 1–12. 10.3389/fpls.2014.00499

110. Nishiyama, T., Sakayama, H., Vries, J. de, Buschmann, H., Saint-Marcoux, D., Ullrich, K. K., Haas, F. B., Vanderstraeten, L., Becker, D., Lang, D., Vosolsobě, S., Rombauts, S., Wilhelmsson, P. K. I., Janitza, P., Kern, R., Heyl, A., Rümpler, F., Villalobos, L. I. A. C., Clay, J. M., … Rensing, S. A. (2018). The Chara Genome: Secondary Complexity and Implications for Plant Terrestrialization. Cell, 174(2), 448–464.e24. 10.1016/j.cell.2018.06.033

111. Nixon, B. T., Mansouri, K., Singh, A., Du, J., Davis, J. K., Lee, J.-G., Slabaugh, E., Vandavasi, V. G., O’Neill, H., Roberts, E. M., Roberts, A. W., Yingling, Y. G., & Haigler, C. H. (2016). Comparative Structural and Computational Analysis Supports Eighteen Cellulose Synthases in the Plant Cellulose Synthesis Complex. Scientific Reports, 6, 28696. 10.1038/srep28696

112. Ogawa-Ohnishi, M., & Matsubayashi, Y. (2015). Identification of three potent hydroxyproline O-galactosyltransferases in Arabidopsis. The Plant Journal, 81(5), 736–746. 10.1111/tpj.12764

113. Ogawa-Ohnishi, M., Matsushita, W., & Matsubayashi, Y. (2013). Identification of three hydroxyproline O-arabinosyltransferases in Arabidopsis thaliana. Nature Chemical Biology, 9(11), 726–730. 10.1038/nchembio.1351

114. Parre, E., & Geitmann, A. (2005). Pectin and the role of the physical properties of the cell wall in pollen tube growth of Solanum chacoense. Planta, 220(4), 582–592. 10.1007/s00425-004-1368-5

115. Pedersen, G. B., Blaschek, L., Frandsen, K. E. H., Noack, L. C., & Persson, S. (2023). Cellulose synthesis in land plants. Molecular Plant, 16(1), 206–231. 10.1016/j.molp.2022.12.015

116. Pedersen, H. L., Fangel, J. U., McCleary, B., Ruzanski, C., Rydahl, M. G., Ralet, M.-C., Farkas, V., von Schantz, L., Marcus, S. E., Andersen, M. C. F., Field, R., Ohlin, M., Knox, J. P., Clausen, M. H., & Willats, W. G. T. (2012). Versatile High Resolution Oligosaccharide Microarrays for Plant Glycobiology and Cell Wall Research. Journal of Biological Chemistry, 287(47), 39429–39438. 10.1074/jbc.M112.396598

117. Peña, M. J., Darvill, A. G., Eberhard, S., York, W. S., & O’Neill, M. A. (2008). Moss and liverwort xyloglucans contain galacturonic acid and are structurally distinct from the xyloglucans synthesized by hornworts and vascular plants. Glycobiology, 18(11), 891–904. 10.1093/glycob/cwn078

118. Peña, M. J., Kong, Y., York, W. S., & O’Neill, M. A. (2012). A galacturonic acid-containing xyloglucan is involved in arabidopsis root hair tip growthw. Plant Cell, 24(11), 4511– 4524. 10.1105/tpc.112.103390

119. Peng, J.-S., Zhang, B.-C., Chen, H., Wang, M.-Q., Wang, Y.-T., Li, H.-M., Cao, S.-X., Yi, H.-Y., Wang, H., Zhou, Y.-H., & Gong, J.-M. (2021). Galactosylation of rhamnogalacturonan-II for cell wall pectin biosynthesis is critical for root apoplastic iron reallocation in Arabidopsis. Molecular Plant, 14(10), 1640–1651. 10.1016/j.molp.2021.06.016

120. Perrin, R. M., DeRocher, A. E., Bar-Peled, M., Zeng, W., Norambuena, L., Orellana, A., Raikhel, N. V., & Keegstra, K. (1999). Xyloglucan Fucosyltransferase, an Enzyme Involved in Plant Cell Wall Biosynthesis. Science, 284(5422), 1976–1979. 10.1126/science.284.5422.1976

121. Petersen, B. L., Egelund, J., Damager, I., Faber, K., Krüger Jensen, J., Yang, Z., Bennett, E. P., Scheller, H. V., & Ulvskov, P. (2009). Assay and heterologous expression in Pichia pastoris of plant cell wall type-II membrane anchored glycosyltransferases. Glycoconjugate Journal, 26(9), 1235–1246. 10.1007/s10719-009-9242-0

122. Pettolino, F. A., Hoogenraad, N. J., Ferguson, C., Bacic, A., Johnson, E., & Stone, B. A. (2001). A (1→4)-β-mannan-specific monoclonal antibody and its use in the immunocytochemical location of galactomannans. Planta, 214(2), 235–242. 10.1007/s004250100606

123. Pettolino, F. A., Walsh, C., Fincher, G. B., & Bacic, A. (2012). Determining the polysaccharide composition of plant cell walls. Nature Protocols, 7(9), 1590–1607. 10.1038/nprot.2012.081

124. Pfeifer, L., Mueller, K.-K., & Classen, B. (2022). The cell wall of hornworts and liverworts: Innovations in early land plant evolution? Journal of Experimental Botany, 73(13), 4454–4472. 10.1093/jxb/erac157

125. Purushotham, P., Ho, R., & Zimmer, J. (2020). Architecture of a catalytically active homotrimeric plant cellulose synthase complex. Science, 369(6507), 1089–1094. 10.1126/science.abb2978

126. Qu, Y., Egelund, J., Gilson, P. R., Houghton, F., Gleeson, P. A., Schultz, C. J., & Bacic, A. (2008). Identification of a novel group of putative Arabidopsis thaliana β-(1,3)-galactosyltransferases. Plant Molecular Biology, 68(1), 43–59. 10.1007/s11103-008-9351-3

127. Rautengarten, C., Ebert, B., Liu, L., Stonebloom, S., Smith-Moritz, A. M., Pauly, M., Orellana, A., Scheller, H. V., & Heazlewood, J. L. (2016). The Arabidopsis Golgi-localized GDP-L-fucose transporter is required for plant development. Nature Communications, 7(1), Article 1. 10.1038/ncomms12119

128. Rautengarten, C., Ebert, B., Moreno, I., Temple, H., Herter, T., Link, B., Donas-Cofre, D., Moreno, A., Saez-Aguayo, S., Blanco, F., Mortimer, J. C., Schultink, A., Reiter, W.-D., Dupree, P., Pauly, M., Heazlewood, J. L., Scheller, H. V., & Orellana, A. (2014). The Golgi localized bifunctional UDP-rhamnose/UDP-galactose transporter family of Arabidopsis. Proceedings of the National Academy of Sciences, 111(31), 11563– 11568. 10.1073/pnas.1406073111

129. Rautengarten, C., Heazlewood, J. L., & Ebert, B. (2019). Profiling Cell Wall Monosaccharides and Nucleotide-Sugars from Plants. Current Protocols in Plant Biology, 4(2), e20092–e20092. 10.1002/cppb.20092

130. Rennie, E. A., Hansen, S. F., Baidoo, E. E. K., Hadi, M. Z., Keasling, J. D., & Scheller, H. V. (2012). Three Members of the Arabidopsis Glycosyltransferase Family 8 Are Xylan Glucuronosyltransferases. Plant Physiology, 159(4), 1408–1417. 10.1104/pp.112.200964

131. Roberts, A. W., Lahnstein, J., Hsieh, Y. S. Y., Xing, X., Yap, K., Chaves, A. M., Scavuzzo-Duggan, T. R., Dimitroff, G., Lonsdale, A., Roberts, E., Bulone, V., Fincher, G. B., Doblin, M. S., Bacic, A., & Burton, R. A. (2018). Functional characterization of a glycosyltransferase from the moss physcomitrella patens involved in the biosynthesis of a novel cell wall arabinoglucan. Plant Cell, 30(6), 1293–1308. 10.1105/tpc.18.00082

132. Rocha, J., Cicéron, F., de Sanctis, D., Lelimousin, M., Chazalet, V., Lerouxel, O., & Breton, C. (2016). Structure of Arabidopsis thaliana FUT1 Reveals a Variant of the GT-B Class Fold and Provides Insight into Xyloglucan Fucosylation. The Plant Cell, 28(10), 2352–2364. 10.1105/tpc.16.00519

133. Ruprecht, C., Bartetzko, M. P., Senf, D., Lakhina, A., Smith, P. J., Soto, M. J., Oh, H., Yang, J. Y., Chapla, D., Varon Silva, D., Clausen, M. H., Hahn, M. G., Moremen, K. W., Urbanowicz, B. R., & Pfrengle, F. (2020). A Glycan Array-Based Assay for the Identification and Characterization of Plant Glycosyltransferases. Angewandte Chemie - International Edition, 59(30), 12493–12498. 10.1002/anie.202003105

134. Ruprecht, C., Dallabernardina, P., Smith, P. J., Urbanowicz, B. R., & Pfrengle, F. (2018). Analyzing Xyloglucan Endotransglycosylases by Incorporating Synthetic Oligosaccharides into Plant Cell Walls. ChemBioChem, 19(8), 793–798. 10.1002/cbic.201700638

135. Saito, F., Suyama, A., Oka, T., Yoko-O, T., Matsuoka, K., Jigami, Y., & Shimma, Y.-I. (2014). Identification of Novel Peptidyl Serine α-Galactosyltransferase Gene Family in Plants. The Journal of Biological Chemistry, 289(30), 20405–20420. 10.1074/jbc.M114.553933

136. Scheller, H. V., & Ulvskov, P. (2010). Hemicelluloses. Annual Review of Plant Biology, 61(1), 263–289. 10.1146/annurev-arplant-042809-112315

137. Schultink, A., Cheng, K., Park, Y. B., Cosgrove, D. J., & Pauly, M. (2013). The identification of two arabinosyltransferases from tomato reveals functional equivalency of xyloglucan side chain substituents. Plant Physiology, 163(1), 86–94. 10.1104/pp.113.221788

138. Seifert, G. J., & Roberts, K. (2007). The Biology of Arabinogalactan Proteins. Annual Review of Plant Biology, 58(1), 137–161. 10.1146/annurev.arplant.58.032806.103801

139. Séveno, M., Séveno-Carpentier, E., Voxeur, A., Menu-Bouaouiche, L., Rihouey, C., Delmas, F., Chevalier, C., Driouich, A., & Lerouge, P. (2010). Characterization of a putative 3-deoxy-d-manno-2-octulosonic acid (Kdo) transferase gene from Arabidopsisthaliana. Glycobiology, 20(5), 617–628. 10.1093/glycob/cwq011

140. Shimamura, M. (2016). Marchantia polymorpha: Taxonomy, phylogeny and morphology of a model system. Plant and Cell Physiology, 57(2), 230–256. 10.1093/pcp/pcv192

141. Showalter, A. M. (2001). Arabinogalactan-proteins: Structure, expression and function. Cellular and Molecular Life Sciences, 58, 1399–1417.

142. Silva, G. B., Ionashiro, M., Carrara, T. B., Crivellari, A. C., Tiné, M. A. S., Prado, J., Carpita, N. C., & Buckeridge, M. S. (2011). Cell wall polysaccharides from fern leaves: Evidence for a mannan-rich Type III cell wall in Adiantum raddianum. Phytochemistry, 72(18), 2352–2360. 10.1016/j.phytochem.2011.08.020

143. Smith, D. K., Jones, D. M., Lau, J. B. R., Cruz, E. R., Brown, E., Harper, J. F., & Wallace, I. S. (2018). A Putative Protein O-Fucosyltransferase Facilitates Pollen Tube Penetration through the Stigma–Style Interface. Plant Physiology, 176(4), 2804– 2818. 10.1104/pp.17.01577

144. Smith, J., Yang, Y., Levy, S., Adelusi, O. O., Hahn, M. G., O’Neill, M. A., & Bar-Peled, M. (2016). Functional Characterization of UDP-apiose Synthases from Bryophytes and Green Algae Provides Insight into the Appearance of Apiose-containing Glycans during Plant Evolution. The Journal of Biological Chemistry, 291(41), 21434–21447. 10.1074/jbc.M116.749069

145. Sterling, J. D., Atmodjo, M. A., Inwood, S. E., Kolli, V. S. K., Quigley, H. F., Hahn, M. G., & Mohnen, D. (2006). Functional identification of an Arabidopsis pectin biosynthetic homogalacturonan galacturonosyltransferase. Proceedings of the National Academy of Sciences of the United States of America, 103(13), 5236–5241. 10.1073/pnas.0600120103

146. Strasser, R., Seifert, G., Doblin, M. S., Johnson, K. L., Ruprecht, C., Pfrengle, F., Bacic, A., & Estevez, J. M. (2021). Cracking the “Sugar Code”: A Snapshot of N- and O-Glycosylation Pathways and Functions in Plants Cells. Frontiers in Plant Science, 12. https://www.frontiersin.org/articles/10.3389/fpls.2021.640919

147. Strother, P. K., & Taylor, W. A. (2018). Chapter 1—The Evolutionary Origin of the Plant Spore in Relation to the Antithetic Origin of the Plant Sporophyte. In M. Krings, C. J. Harper, N. R. Cúneo, & G. W. Rothwell (Eds.), Transformative Paleobotany (pp. 3–20). Academic Press. 10.1016/B978-0-12-813012-4.00001-2

148. Suzuki, T., Narciso, J. O., Zeng, W., van de Meene, A., Yasutomi, M., Takemura, S., Lampugnani, E. R., Doblin, M. S., Bacic, A., & Ishiguro, S. (2017). KNS4/UPEX1: A Type II Arabinogalactan β-(1,3)-Galactosyltransferase Required for Pollen Exine Development. Plant Physiology, 173(1), 183–205. 10.1104/pp.16.01385

149. Takenaka, Y., Kato, K., Ogawa-Ohnishi, M., Tsuruhama, K., Kajiura, H., Yagyu, K., Takeda, A., Takeda, Y., Kunieda, T., Hara-Nishimura, I., Kuroha, T., Nishitani, K., Matsubayashi, Y., & Ishimizu, T. (2018). Pectin RG-I rhamnosyltransferases represent a novel plant-specific glycosyltransferase family. Nature Plants, 4(9), 669– 676. 10.1038/s41477-018-0217-7

150. Tan, L., Eberhard, S., Pattathil, S., Warder, C., Glushka, J., Yuan, C., Hao, Z., Zhu, X., Avci, U., Miller, J. S., Baldwin, D., Pham, C., Orlando, R., Darvill, A., Hahn, M. G., Kieliszewski, M. J., & Mohnen, D. (2013). An Arabidopsis Cell Wall Proteoglycan Consists of Pectin and Arabinoxylan Covalently Linked to an Arabinogalactan Protein. The Plant Cell, 25(1), 270–287. 10.1105/tpc.112.107334

151. Tan, L., Showalter, A. M., Egelund, J., Hernandez-Sanchez, A., Doblin, M. S., & Bacic, A. (2012). Arabinogalactan-proteins and the research challenges for these enigmatic plant cell surface proteoglycans. Frontiers in Plant Science, 3. 10.3389/fpls.2012.00140

152. Tan, L., Zhang, L., Black, I., Glushka, J., Urbanowicz, B., Heiss, C., & Azadi, P. (2022). Most of the rhamnogalacturonan-I from cultured Arabidopsis cell walls is covalently linked to arabinogalactan-protein. Carbohydrate Polymers, 301, 120340. 10.1016/j.carbpol.2022.120340

153. Tang, S. N., Barnum, C. R., Szarzanowicz, M. J., Sirirungruang, S., & Shih, P. M. (2023). Harnessing Plant Sugar Metabolism for Glycoengineering. Biology, 12(12), Article 12. 10.3390/biology12121505

154. Tenhaken, R. (2015). Cell wall remodeling under abiotic stress. Frontiers in Plant Science, 5(JAN), 1–9. 10.3389/fpls.2014.00771

155. The International Brachypodium Initiative. (n.d.). Genome sequencing and analysis of the model grass Brachypodium distachyon | Nature. Retrieved 18 February 2025, from https://www.nature.com/articles/nature08747

156. Uehara, Y., Tamura, S., Maki, Y., Yagyu, K., Mizoguchi, T., Tamiaki, H., Imai, T., Ishii, T., Ohashi, T., Fujiyama, K., & Ishimizu, T. (2017). Biochemical characterization of rhamnosyltransferase involved in biosynthesis of pectic rhamnogalacturonan I in plant cell wall. Biochemical and Biophysical Research Communications, 486(1), 130–136. 10.1016/j.bbrc.2017.03.012

157. Urbanowicz, B. R., Bharadwaj, V. S., Alahuhta, M., Peña, M. J., Lunin, V. V., Bomble, Y. J., Wang, S., Yang, J.-Y., Tuomivaara, S. T., Himmel, M. E., Moremen, K. W., York, W. S., & Crowley, M. F. (2017). Structural, mutagenic and in silico studies of xyloglucan fucosylation in Arabidopsis thaliana suggest a water-mediated mechanism. The Plant Journal, 91(6), 931–949. 10.1111/tpj.13628

158. Urbanowicz, B. R., Peña, M. J., Moniz, H. A., Moremen, K. W., & York, W. S. (2014). Two Arabidopsis proteins synthesize acetylated xylan in vitro. The Plant Journal, 80(2), 197–206. 10.1111/tpj.12643

159. Vanzin, G. F., Madson, M., Carpita, N. C., Raikhel, N. V., Keegstra, K., & Reiter, W.-D. (2002). The mur2 mutant of Arabidopsis thaliana lacks fucosylated xyloglucan because of a lesion in fucosyltransferase AtFUT1. Proceedings of the National Academy of Sciences of the United States of America, 99(5), 3340–3345. 10.1073/pnas.052450699

160. Vega-Sánchez, M. E., Verhertbruggen, Y., Christensen, U., Chen, X., Sharma, V., Varanasi, P., Jobling, S. A., Talbot, M., White, R. G., Joo, M., Singh, S., Auer, M., Scheller, H. V., & Ronald, P. C. (2012). Loss of Cellulose synthase-like F6 function affects mixed-linkage glucan deposition, cell wall mechanical properties, and defense responses in vegetative tissues of rice. Plant Physiology, 159(1), 56–69. 10.1104/pp.112.195495

161. Vega-Sánchez, M. E., Verhertbruggen, Y., Scheller, H. V., & Ronald, P. C. (2013). Abundance of mixed linkage glucan in mature tissues and secondary cell walls of grasses. Plant Signaling & Behavior, 8(2), e23143. 10.4161/psb.23143

162. Velasquez, S. M., Ricardi, M. M., Dorosz, J. G., Fernandez, P. V., Nadra, A. D., Pol-Fachin, L., Egelund, J., Gille, S., Harholt, J., Ciancia, M., Verli, H., Pauly, M., Bacic, A., Olsen, C. E., Ulvskov, P., Petersen, B. L., Somerville, C., Iusem, N. D., & Estevez, J. M. (2011). O-glycosylated cell wall proteins are essential in root hair growth. Science, 332(6036), 1401–1403. 10.1126/science.1206657

163. Verhertbruggen, Y., Yin, L., Oikawa, A., & Scheller, H. V. (2011). Mannan synthase activity in the CSLD family. Plant Signaling & Behavior, 6(10), 1620–1623. 10.4161/psb.6.10.17989

164. Voiniciuc, C. (2022). Modern mannan: A hemicellulose’s journey. New Phytologist, 234(4), 1175–1184. 10.1111/nph.18091

165. Voiniciuc, C., Dama, M., Gawenda, N., Stritt, F., & Pauly, M. (2019). Mechanistic insights from plant heteromannan synthesis in yeast. Proceedings of the National Academy of Sciences, 116(2), 522–527. 10.1073/pnas.1814003116

166. Voiniciuc, C., Engle, K. A., Günl, M., Dieluweit, S., Schmidt, M. H.-W., Yang, J.-Y., Moremen, K. W., Mohnen, D., & Usadel, B. (2018). Identification of Key Enzymes for Pectin Synthesis in Seed Mucilage. Plant Physiology, 178(3), 1045–1064. 10.1104/pp.18.00584

167. Voiniciuc, C., Schmidt, M. H.-W., Berger, A., Yang, B., Ebert, B., Scheller, H. V., North, H. M., Usadel, B., & Günl, M. (2015). MUCILAGE-RELATED10 Produces Galactoglucomannan That Maintains Pectin and Cellulose Architecture in Arabidopsis Seed Mucilage. Plant Physiology, 169(1), 403–420. 10.1104/pp.15.00851

168. Voxeur, A., Soubigou-Taconnat, L., Legée, F., Sakai, K., Antelme, S., Durand-Tardif, M., Lapierre, C., & Sibout, R. (2017). Altered lignification in mur1-1 a mutant deficient in GDP-L-fucose synthesis with reduced RG-II cross linking. PLOS ONE, 12(9), e0184820. 10.1371/journal.pone.0184820

169. Vuttipongchaikij, S., Brocklehurst, D., Steele-King, C., Ashford, D. A., Gomez, L. D., & McQueen-Mason, S. J. (2012). Arabidopsis GT34 family contains five xyloglucan α-1,6-xylosyltransferases. The New Phytologist, 195(3), 585–595. 10.1111/j.1469-8137.2012.04196.x

170. Wachananawat, B., Kuroha, T., Takenaka, Y., Kajiura, H., Naramoto, S., Yokoyama, R., Ishizaki, K., Nishitani, K., & Ishimizu, T. (2020). Diversity of Pectin Rhamnogalacturonan I Rhamnosyltransferases in Glycosyltransferase Family 106. Frontiers in Plant Science, 11(July), 1–12. 10.3389/fpls.2020.00997

171. Wang, B., Andargie, M., & Fang, R. (2022). The function and biosynthesis of callose in high plants. Heliyon, 8(4), e09248. 10.1016/j.heliyon.2022.e09248

172. Wang, H., Yang, H., Wen, Z., Gao, C., Gao, Y., Tian, Y., Xu, Z., Liu, X., Persson, S., Zhang, B., & Zhou, Y. (2022). Xylan-based nanocompartments orchestrate plant vessel wall patterning. Nature Plants, 8(3), 295–306. 10.1038/s41477-022-01113-1

173. Wang, Y., Mortimer, J. C., Davis, J., Dupree, P., & Keegstra, K. (2013). Identification of an additional protein involved in mannan biosynthesis. The Plant Journal, 73(1), 105–117. 10.1111/tpj.12019

174. Wu, Y., Williams, M., Bernard, S., Driouich, A., Showalter, A. M., & Faik, A. (2010). Functional identification of two nonredundant Arabidopsis α(1,2)fucosyltransferases specific to arabinogalactan proteins. Journal of Biological Chemistry, 285(18), 13638–13645. 10.1074/jbc.M110.102715

175. Xie, B., Wang, X., Zhu, M., Zhang, Z., & Hong, Z. (2011). CalS7 encodes a callose synthase responsible for callose deposition in the phloem. The Plant Journal, 65(1), 1–14. 10.1111/j.1365-313X.2010.04399.x

176. Yang, J., Bak, G., Burgin, T., Barnes, W. J., Mayes, H. B., Peña, M. J., Urbanowicz, B. R., & Nielsen, E. (2020). Biochemical and Genetic Analysis Identify CSLD3 as a beta-1,4-Glucan Synthase That Functions during Plant Cell Wall Synthesis[OPEN]. The Plant Cell, 32(5), 1749–1767. 10.1105/tpc.19.00637

177. Yin, L., Verhertbruggen, Y., Oikawa, A., Manisseri, C., Knierim, B., Prak, L., Jensen, J. K., Knox, J. P., Auer, M., Willats, W. G. T., & Scheller, H. V. (2011). The Cooperative Activities of CSLD2, CSLD3, and CSLD5 Are Required for Normal *Arabidopsis* Development. Molecular Plant, 4(6), 1024–1037. 10.1093/mp/ssr026

178. Zeng, W., Jiang, N., Nadella, R., Killen, T. L., Nadella, V., & Faik, A. (2010). A Glucurono(arabino)xylan synthase complex from wheat contains members of the GT43, GT47, and GT75 families and functions cooperatively. Plant Physiology, 154(1), 78–97. 10.1104/pp.110.159749

179. Zeng, W., Lampugnani, E. R., Picard, K. L., Song, L., Wu, A.-M., Farion, I. M., Zhao, J., Ford, K., Doblin, M. S., & Bacic, A. (2016). Asparagus IRX9, IRX10, and IRX14A Are Components of an Active Xylan Backbone Synthase Complex that Forms in the Golgi Apparatus. Plant Physiology, 171(1), 93–109. 10.1104/pp.15.01919

180. Zhang, N., Julian, J. D., Yap, C. E., Swaminathan, S., & Zabotina, O. A. (2023). The Arabidopsis xylosyltransferases, XXT3, XXT4, and XXT5, are essential to complete the fully xylosylated glucan backbone XXXG-type structure of xyloglucans. New Phytologist, 238(5), 1986–1999. 10.1111/nph.18851

181. Zhang, Y., Sharma, D., Liang, Y., Downs, N., Dolman, F., Thorne, K., Black, I. M., Pereira, J. H., Adams, P., Scheller, H. V., O’Neill, M., Urbanowicz, B., & Mortimer, J. C. (2024). Putative rhamnogalacturonan-II glycosyltransferase identified through callus gene editing which bypasses embryo lethality. *Plant Physiology*, kiae259. 10.1093/plphys/kiae259

182. Zhu, L., Dama, M., & Pauly, M. (2018). Identification of an arabinopyranosyltransferase from Physcomitrella patens involved in the synthesis of the hemicellulose xyloglucan. Plant Direct, 2(3). 10.1002/pld3.46

