## Supplementary figures and images for "The Building Blocks of Early Land Plants: Glycosyltransferases and Cell Wall Architecture in the model liverwort *Marchantia polymorpha*"

### Figure S2

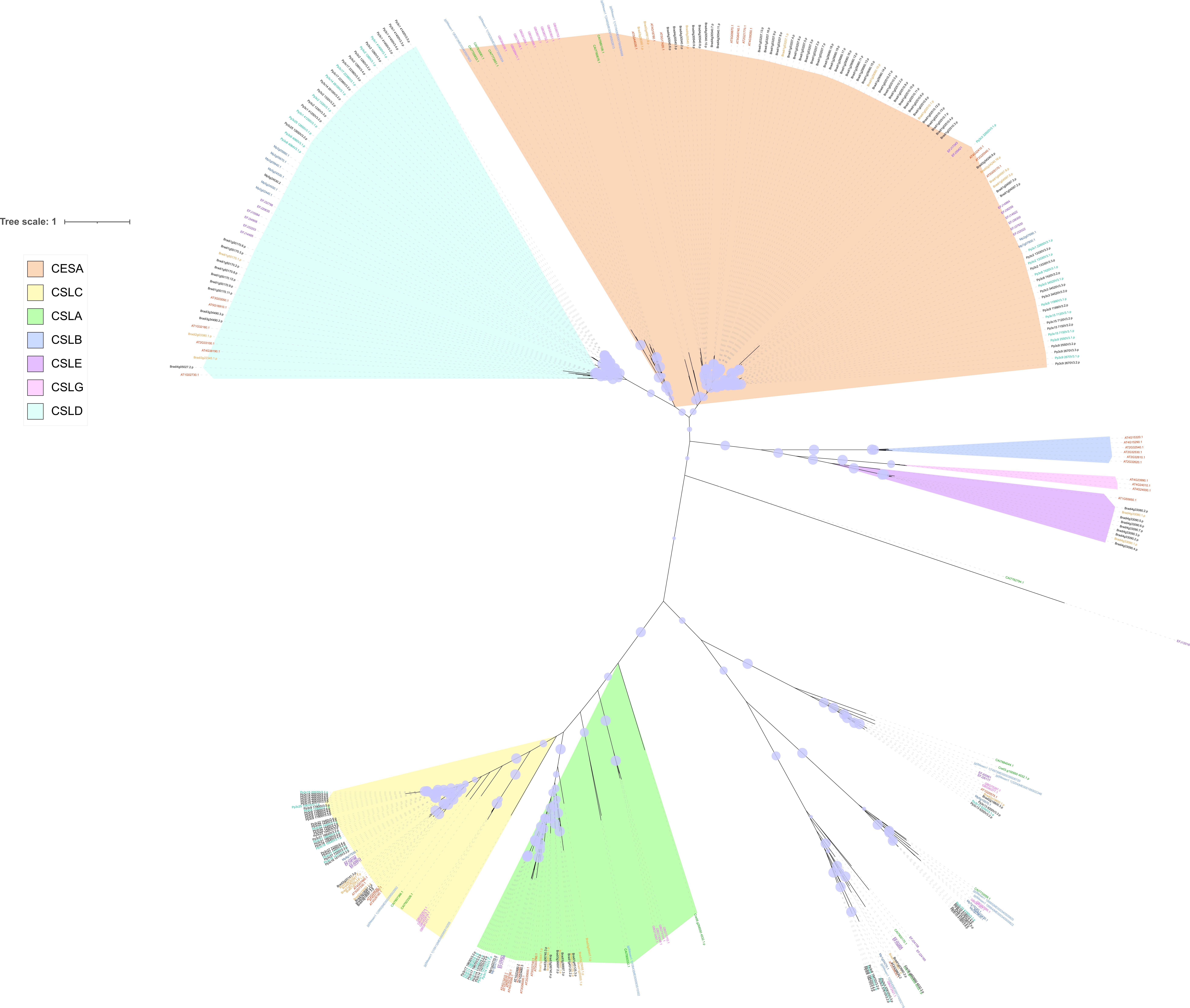

### Figure S5

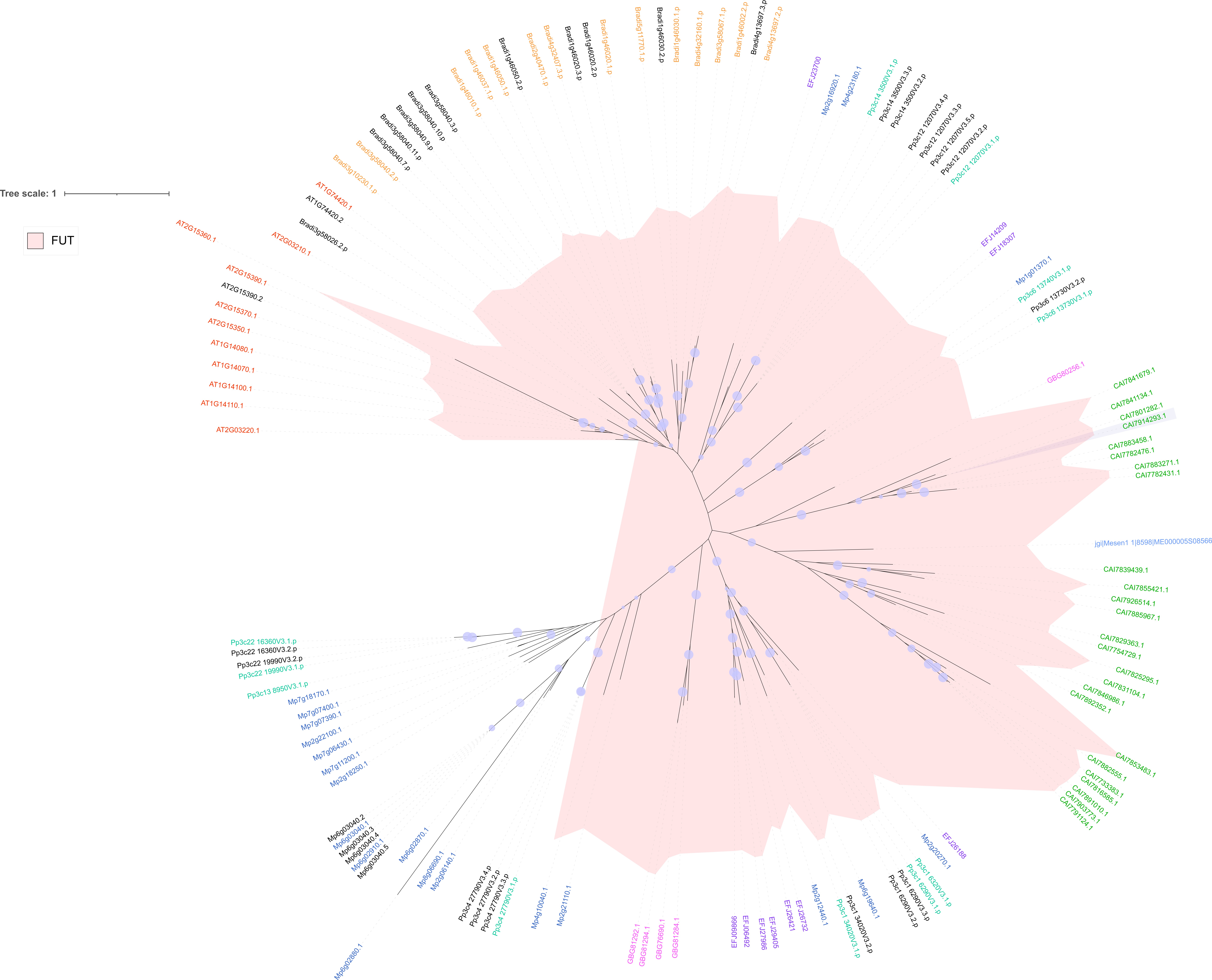

### Figure S6

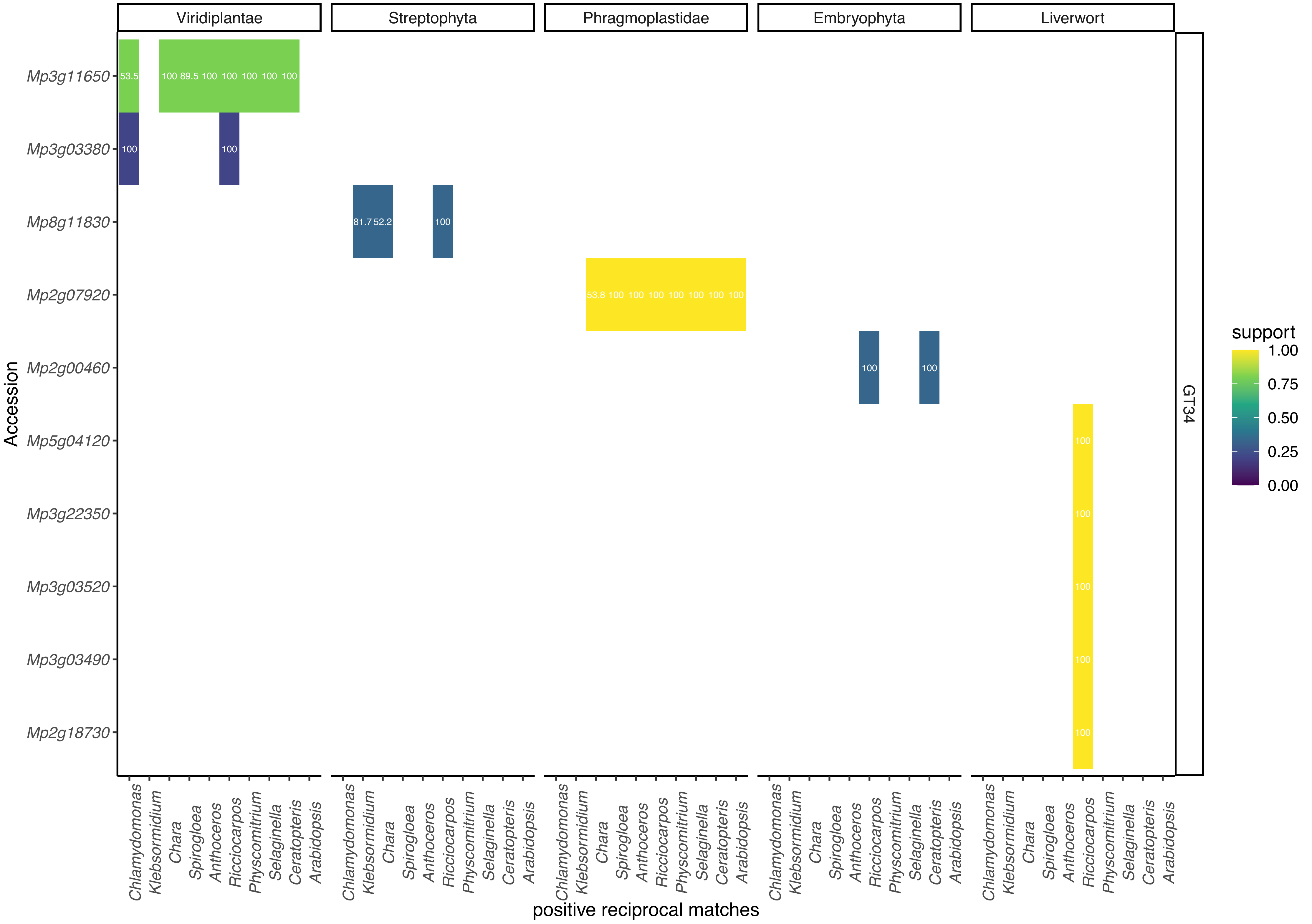

### Figure S7

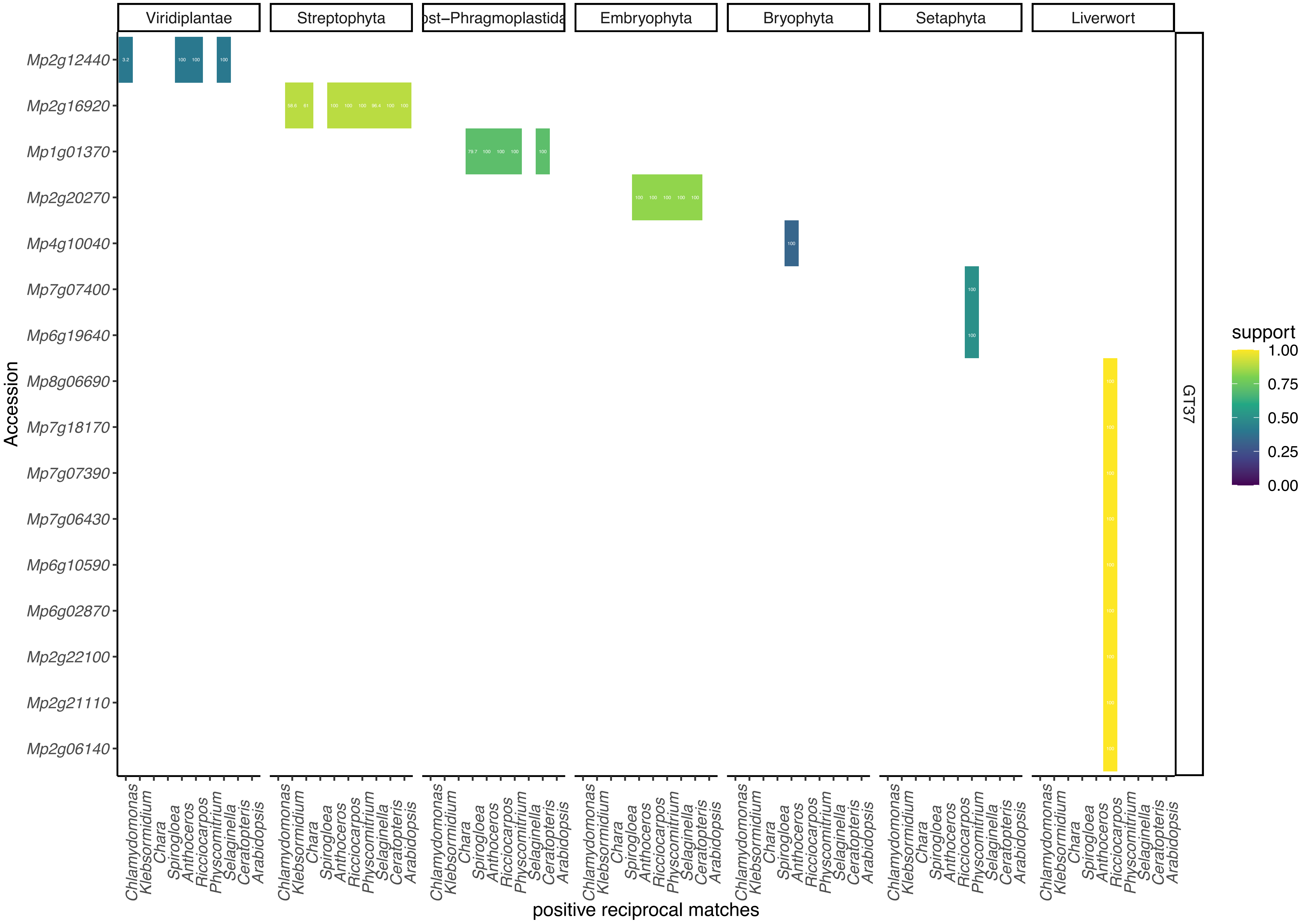

### Figure S8

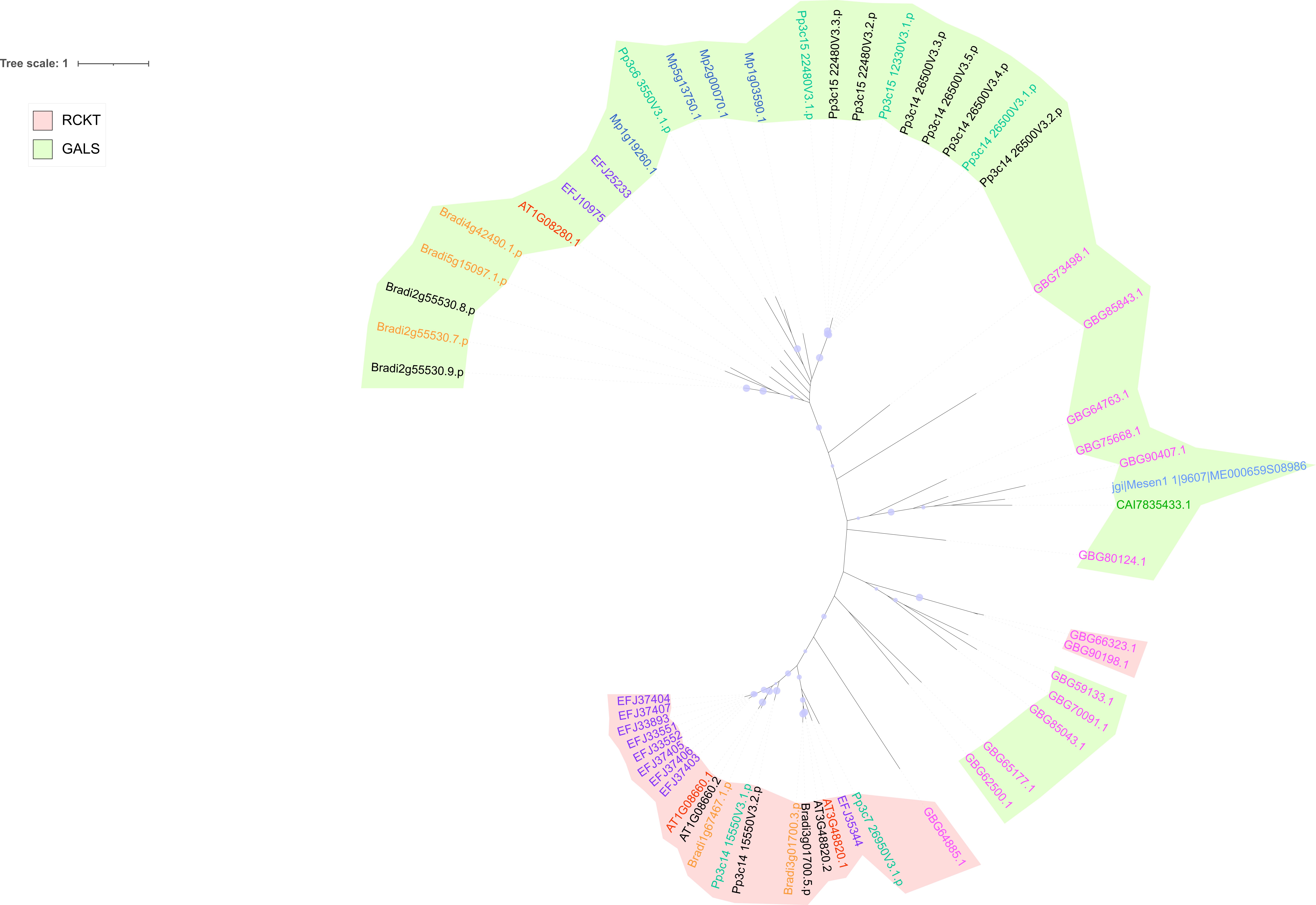

### Figure S9

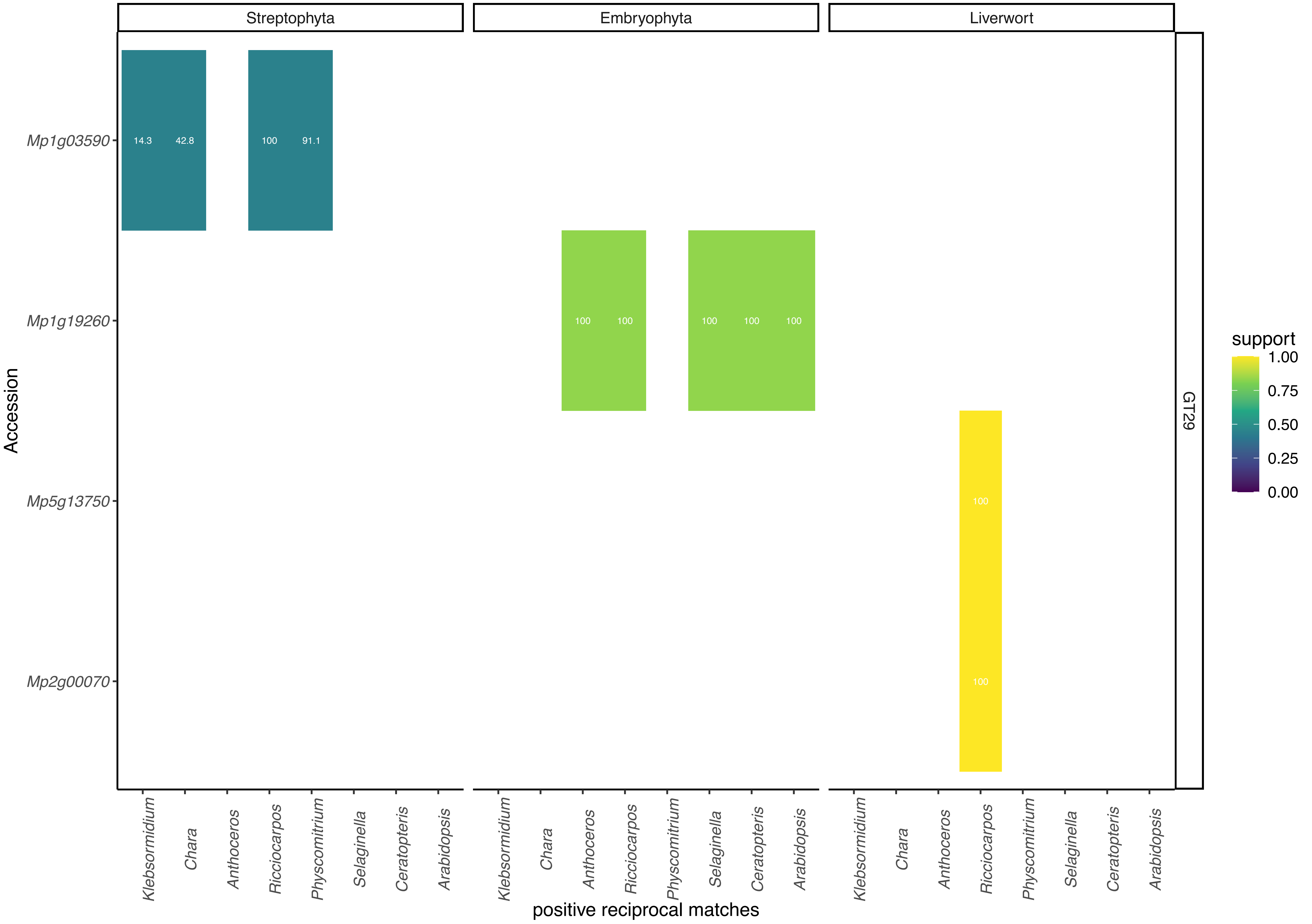

### Figure S10

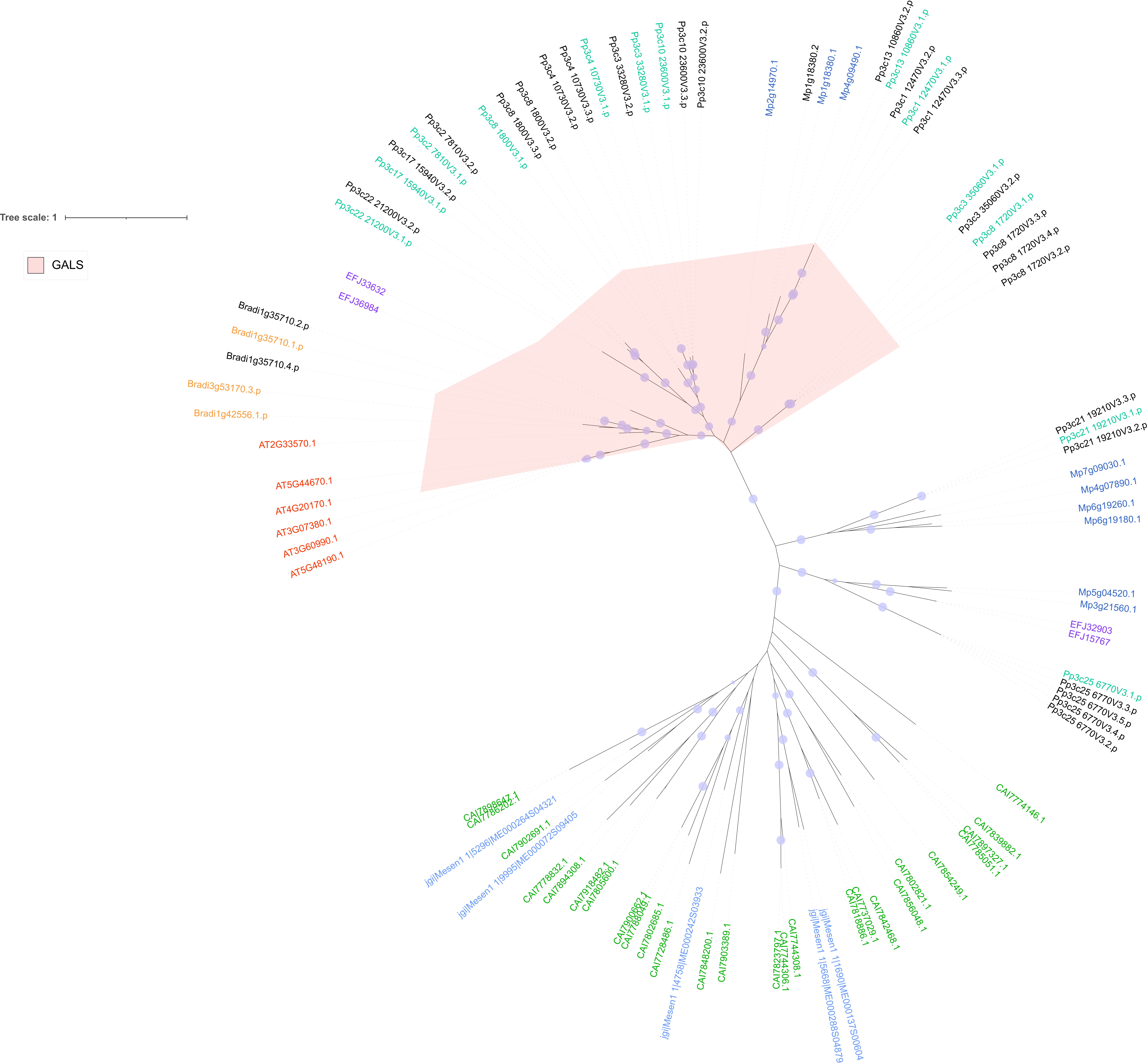

### Figure S11

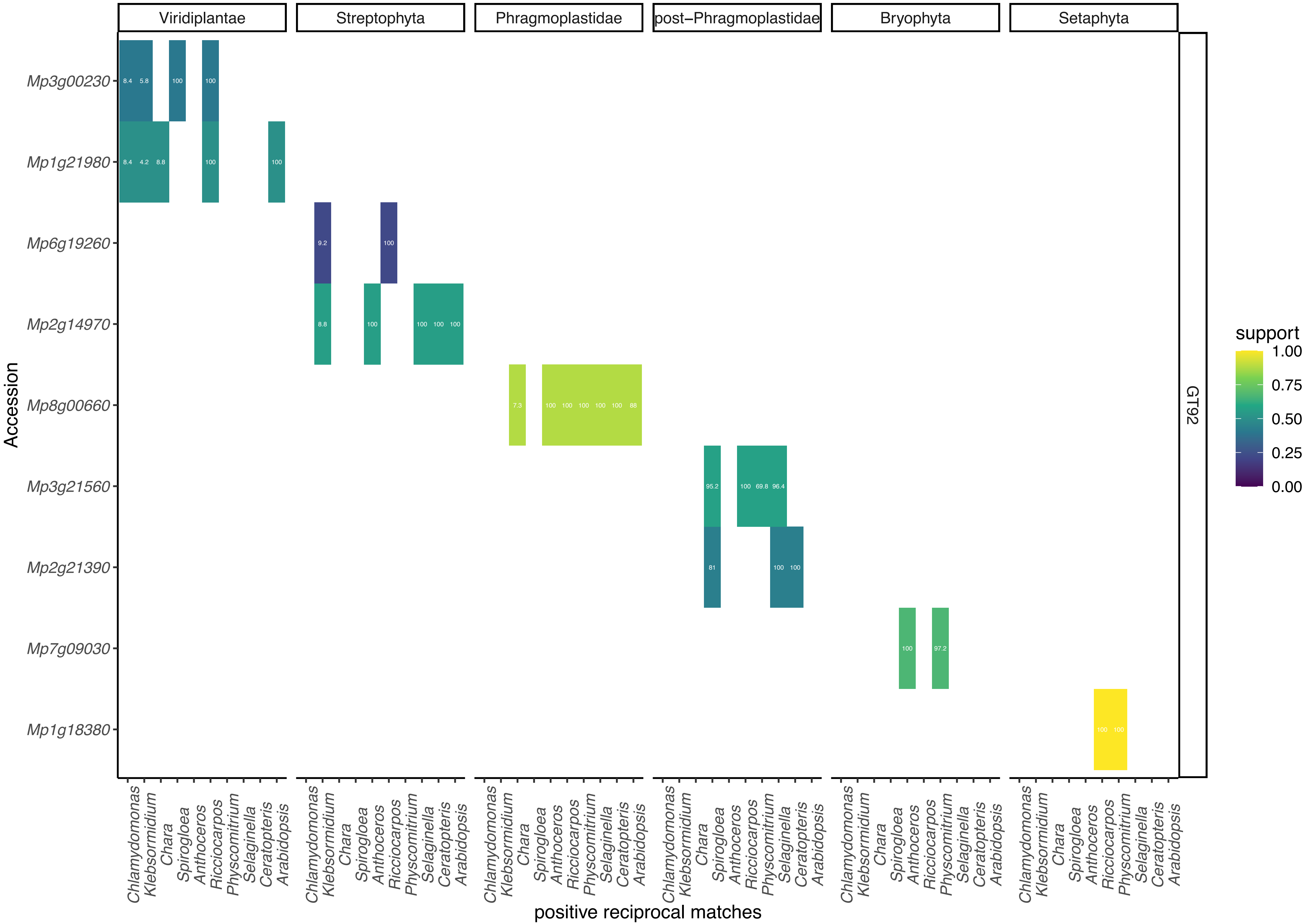

### Figure S12

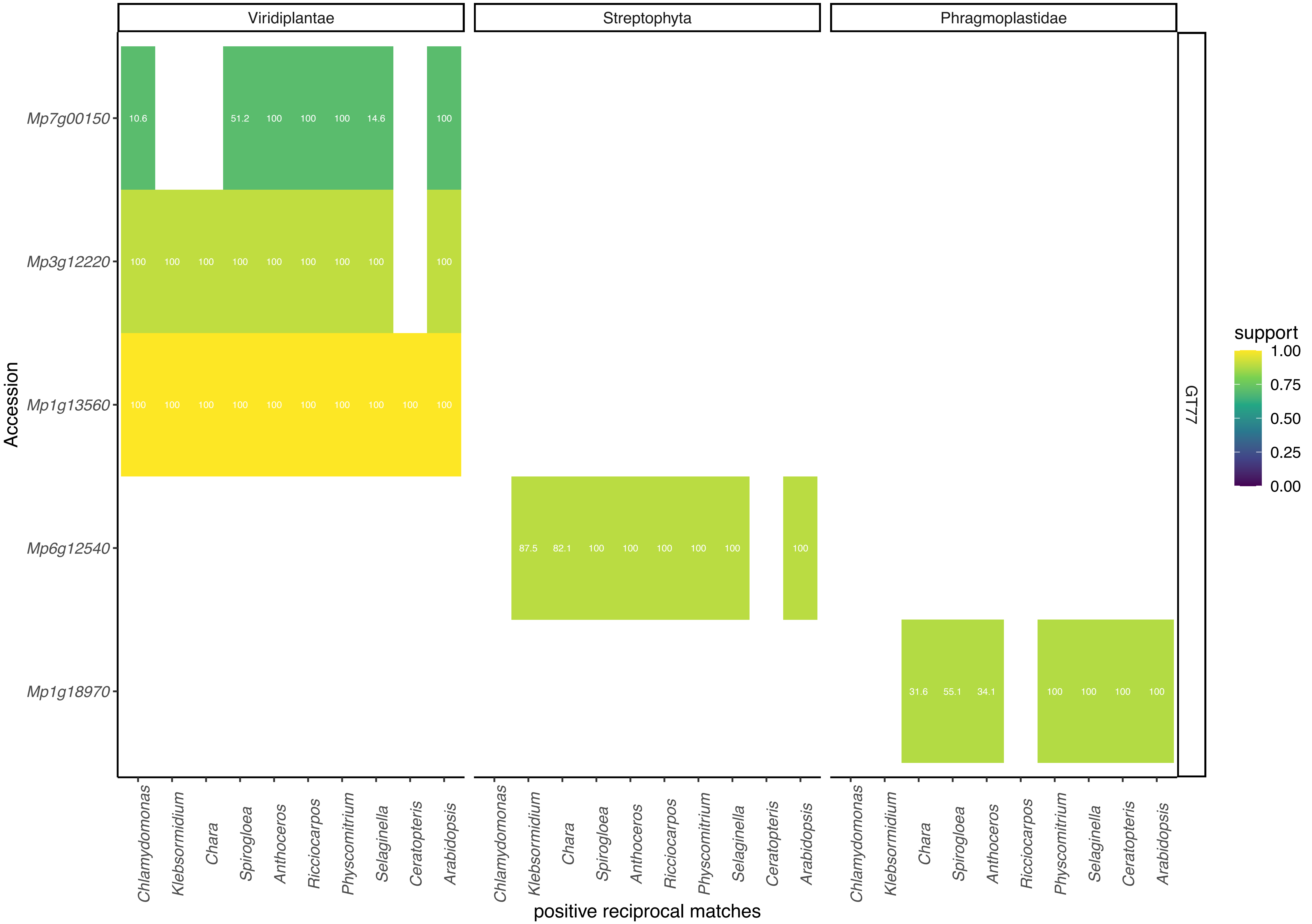

### Figure S13

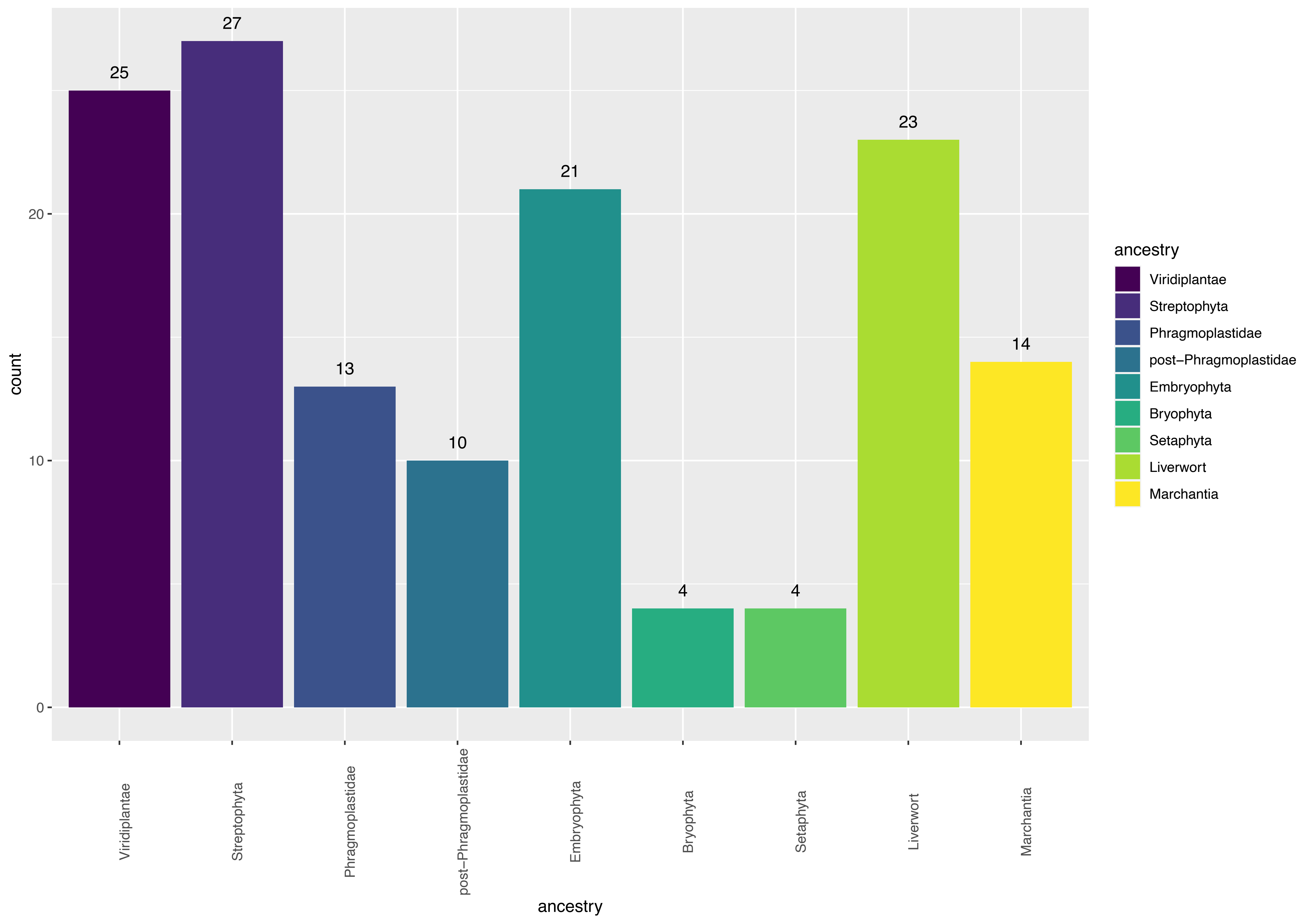

### Figures S3

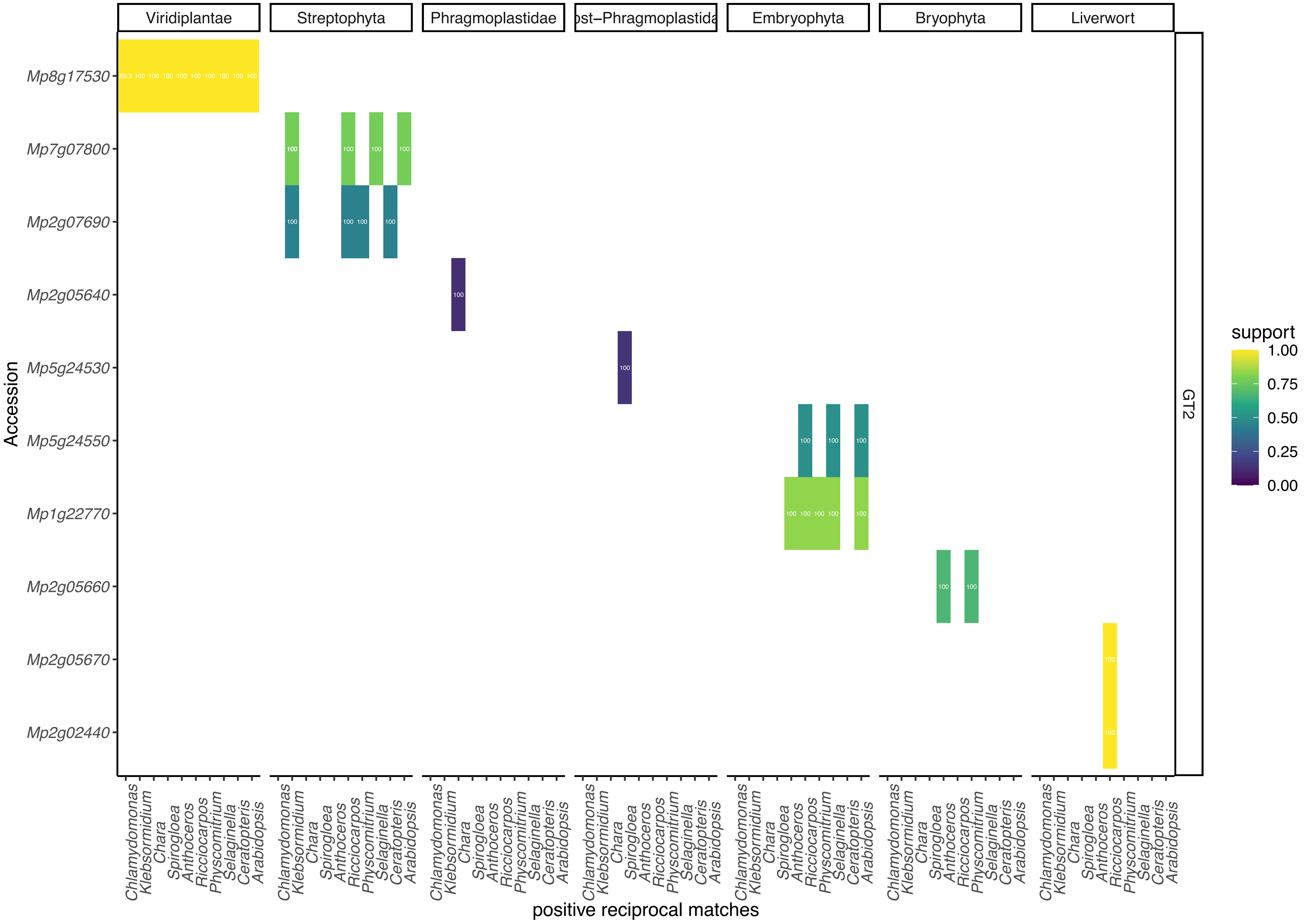

### Figures S4

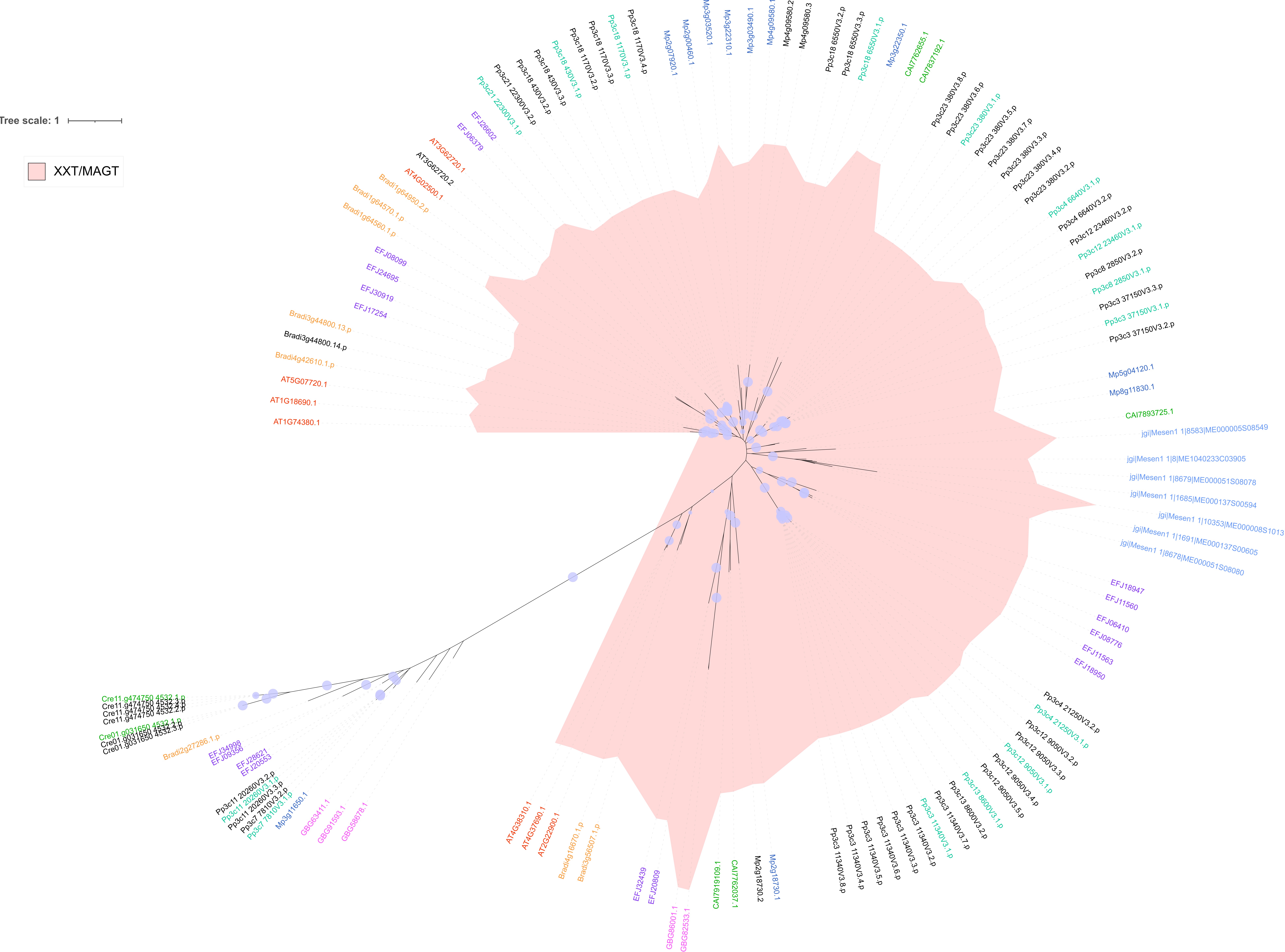
