## Supplementary material for "The Building Blocks of Early Land Plants: Glycosyltransferases and Cell Wall Architecture in the model liverwort *Marchantia polymorpha*": Figure S1

Figure S1. Monosaccharide abundance (mol%) deduced from the glycosyl linkage data (Table S2).

mol%

|  | Dormant Gemma | 8d Gemmaling | Mature Thallus | Antheridiophore | Archegoniophore Stem | Archegoniophore | Sporophyte | 7d Cultured Sporellings |
| --- | --- | --- | --- | --- | --- | --- | --- | --- |
| Total Glc | 49.2 | 49.6 | 58.5 | 62.2 | 63.7 | 59.3 | 48.5 | 51 |

mol%

| Monosaccharide | Dormant Gemma | 8d Gemmaling | Mature Thallus | Antheridiophore | Archegoniophore Stem | Archegoniophore | Sporophyte | 7d Cultured Sporellings |
| --- | --- | --- | --- | --- | --- | --- | --- | --- |
| Total Ara | 7.8 | 9.8 | 12.6 | 6.2 | 4.8 | 5.6 | 20.6 | 7.8 |
| Total Xyl | 9.9 | 8.3 | 7.5 | 5.8 | 5.2 | 11.4 | 15.3 | 7.2 |
| Total Rha | 1.7 | 0.6 | 0.8 | 0.8 | 0.6 | 0.7 | 1.3 | 2.1 |
| Total Fuc | 0.3 | 0.2 | 1.7 | 1.3 | 0.6 | 0.9 | 0.4 | 0.1 |
| Total Man | 5.4 | 8.2 | 6.5 | 10.3 | 14.3 | 10.5 | 3.8 | 4.7 |
| Total Glc A | 1.4 | 0.8 | 0 | 0 | 0.1 | 0 | 0 | 0.6 |
| Total Gal | 22.4 | 19.4 | 11.3 | 11.8 | 9.7 | 10.1 | 8.4 | 23.1 |
| Total Gal A | 2.8 | 3.6 | 1 | 1.6 | 1 | 1.3 | 1.9 | 4.2 |
